# Identification of novel prognostic targets in coronary artery disease and related complications using bioinformatics and next generation sequencing data analysis

**DOI:** 10.1101/2023.02.22.529500

**Authors:** Basavaraj Vastrad, Chanabasayya Vastrad

**Affiliations:** Department of Pharmaceutical Chemistry, K.L.E. College of Pharmacy, Gadag, Karnataka 582101, India; Biostatistics and Bioinformatics, Chanabasava Nilaya, Bharthinagar, Dharwad 580001, Karnataka, India

**Author notes:** Chanabasayya Vastrad, Chanabasava Nilaya, Bharthinagar, Dharwad 580001, Karanataka, India.

**Keywords:** Coronary artery disease, Bioinformatics analysis, Differentially expressed genes, Biomarkers, Protein-protein network analysis

## Abstract

Coronary artery disease (CAD) is the most common cardiovascular disorder and the leading cause of heart-related deaths in world. Increasing molecular targets have been discovered for CAD and CAD - related complications prognosis and therapy. However, there is still an urgent need to identify novel biomarkers. Therefore, we evaluated biomarkers that might help the diagnosis and treatment of CAD and CAD - related complications. We searched next generation sequencing (NGS) dataset (GSE202625) and identified differentially expressed genes (DEGs) by comparing CAD and normal control samples using DESeq2. Gene ontology (GO) and pathway enrichment analyses of the DEGs were performed using the g:Profiler online database. The protein-protein interaction (PPI) network was plotted with IMEx interactome and visualized using Cytoscape. Module analysis of the PPI network was done using PEWCC1. MiRNA - hub gene regulatory network and TF - hub gene regulatory network analysis was performed to identify the hub genes, miRNAs and TFs. Receiver operating characteristic (ROC) curve analysis was used to predict the diagnostic effectiveness of the hub genes. A total of 118 DEGs (479 up regulated genes and 479 down regulated genes) were detected. The GO enrichment analysis indicated that the DEGs most significantly enriched in cellular response to stimulus and biosynthetic process. The REACTOME pathway enrichment analysis revealed that the DEGs were most significantly enriched in immune system and eukaryotic translation elongation. PPI network, modules, miRNA - hub gene regulatory network and TF - hub gene regulatory network analysis demonstrated that EGR1, SIRT1, STAT1, LRRK2, HIF1A, CSNK2B, RPS3, RPS2, RPS4X and HDAC11 were the hub genes. On the whole, the findings of this study enhance our understanding of the potential molecular mechanisms of CAD and CAD-related complications, and provide potential targets for further research.

## Introduction

Coronary artery disease (CAD) refers to the complete obstruction of blood flow through the coronary arteries occurs resulting in insufficient oxygen and nutrient delivery to the heart muscles, potentially causing the death of the heart muscle cells and subsequent dysfunction of a portion of the heart muscle itself [1]. CAD and CAD-related complications have become key causes of mortality and morbidity through the world [2]. Incidence of CAD accounts for 8.14 million deaths (16.8%) globally [3]. Myocardial infarction [4], atherosclerosis [5], arrhythmias [6], heart failure [7], atrial fibrillation [8] and dilated cardiomyopathy [9] are associated with CAD. Insulin resistance [10], type 1 and type 2 diabetes mellitus [11–12], non-alcoholic fatty liver disease [13], hypertension [14], obesity [15] and hypercholesterolemia [16] are thought to be the contributing complications to CAD. However, the molecular mechanism underlying many CAD cases remains unclear, resulting in a lack of effective treatment [17]. The higher pervasiveness and finite treatments of CAD lead to considerable public health and economic hardship [18]. Therefore, it is necessary to improve our understanding of CAD molecular pathogenesis and to advance better screening methods for CAD.

In recent years, scholars worldwide have made remarkable achievements in biomarkers and aberrant molecular signaling pathways are involved in the CAD. Data identified several signaling pathways include NF-κB signaling pathway [19], RhoA/ROCK-1 signaling pathway [20], Wnt signaling pathway [21], TLR4 signaling pathway [22] and transforming growth factor β signaling pathway [23], and genes include SCARB1 [24], TP53 [25], RRM1, RRM2 and ERCC2 [26], IL17RA [27] and COX2 [28] that were significantly associated with CAD. In particular, the novel molecular characteristics can be tested in initial risk assessment, the recognition of better specific biomarkers, and the advancement of clinic treatment and survival.

The fast development of next generation sequencing (NGS) technology and bioinformatics analysis based on high-throughput data provide new tactics to identify differentially expressed genes (DEGs) and discover therapeutic targets for the initiation and evolution of CAD and CAD-related complications [29]. However, the differentially expressed genes (DEGs) identified with NGS data large samples and unknown confounding factors. To identify new important DEGs that remained representative and accurate, we used integrated bioinformatics methods genes obtained from NGS data.

In the current investigation, we downloaded NGS dataset, namely, GSE202625 [30] from the NCBI Gene Expression Omnibus (GEO) database (https://www.ncbi.nlm.nih.gov/geo/) [31], including NGS data. We identified DEGs using the DESeq2 package in R bioconductor. The Gene ontology (GO) and pathway enrichment analyses were performed via g:Profiler. Subsequently, we integrated the DEG protein-protein interaction (PPI) network with module screening to identify hub genes in CAD. Subsequently we constructed miRNA-hub gene regulatory network and TF-hub gene regulatory network to identify hub genes, miRNA and TFs. Finally, we validated the hub genes by receiver operating characteristic (ROC) curve analysis. Identifying key genes and their enriched functions and signaling pathways might help provide a convenient and effective approach for clinical prevention, treatment and management of CAD and CAD-related complications.

## Materials and Methods

### Next generation sequencing data source

NGS dataset was downloaded from the GEO database. GSE202625 [30] dataset includes 76 CAD samples and 62 normal control samples. The GSE202625 dataset based on GPL23934 Ion Torrent S5 (Homo sapiens).

### Identification of DEGs

DESeq2 [32] an R package from Bioconductor was used to identify DEGs between samples from patients with CAD and normal controls. We defined DEGs as a fold change (FC) > 0.333 for up regulated genes and fold change (FC) < -0.2803 for down regulated genes. The Benjamini and Hochberg (BH) method was accomplished to adjust P value to cut down the false positive error [33]. A P value□<□0.05 was considered statistically significant. ggplot2 and gplot packages of R software was applied to generate volcano plot and heatmap.

### GO and pathway enrichment analyses of DEGs

GO enrichment analysis (GO, http://www.geneontology.org) [34] which means gene ontology analysis now is prevalently used to define genes and it is an actual useful method for our daily extensive scale of functional enrichment study. GO enrichment analysis can be classified into three different gene functions includes biological process (BP), molecular function (MF), and cellular component (CC). REACTOME (https://reactome.org/) [35] pathway database keeps a lot of data about biological pathways, diseases, and chemical substances. In our study, we used the database for g:Profiler (http://biit.cs.ut.ee/gprofiler/) [36] to analyze DEG enrichment of BP, CC, and MF and the REACTOME pathways. P < 0.05 was considered statistically significant.

### Construction of the PPI network and module analysis

In order to find the hub genes in CAD, we integrated the cross correlation of DEGs through the IMEx interactome (https://www.imexconsortium.org/) [37] database. Later, with the help of the Cytoscape (http://www.cytoscape.org/) [38] software, a PPI network of DEGs was constructed, which was helpful for systematic study on the molecular mechanisms of diseases and therapeutic targets. Finally, for the sake of screening out the hub genes from the PPI network according to the degree [39], betweenness [40], stress [41] and closeness [42] values in the Cytoscape plugin Network Analyzer. The PEWCC1 [43] of Cytoscape was carried out to module analyze and visualize the result of PPI network.

### Construction of the miRNA-hub gene regulatory network

miRNAs can play a role in maintaining physiological stability by regulating the expression of hub genes. We used miRNet database (https://www.mirnet.ca/) [44] to find miRNAs regulating hub genes, taking the intersection of the results of this database. In addition, we visualized miRNA-hub gene regulatory network with Cytoscape software [38].

### Construction of the TF-hub gene regulatory network

TFs can play a role in maintaining physiological stability by regulating the expression of hub genes. We used NetworkAnalyst database (https://www.networkanalyst.ca/) [45] database to find TFs regulating hub genes, taking the intersection of the results of this database. In addition, we visualized TF-hub gene regulatory network with Cytoscape software [38].

### Receiver operating characteristic curve (ROC) analysis

ROC analysis was performed to predict the diagnostic effectiveness of biomarkers for CAD by pROC package in R software [46]. The area under the ROC curve (AUC) value was utilized to determine the diagnostic effectiveness in discriminating CAD from normal control samples in the GSE64566 dataset.

## Results

### Identification of DEGs

Comparing the genes of CAD samples with normal control samples in GSE202625, we found 958 DEGs, including 479 up regulated and 479 down regulated genes (Table 1). The volcano plot of the distribution of DEGs in GSE202625 is shown in Fig. 1. Hierarchical clustering analysis revealed a clear distinction of DEGs between patients with CAD and normal controls (Fig. 2).

**Fig. 1.**
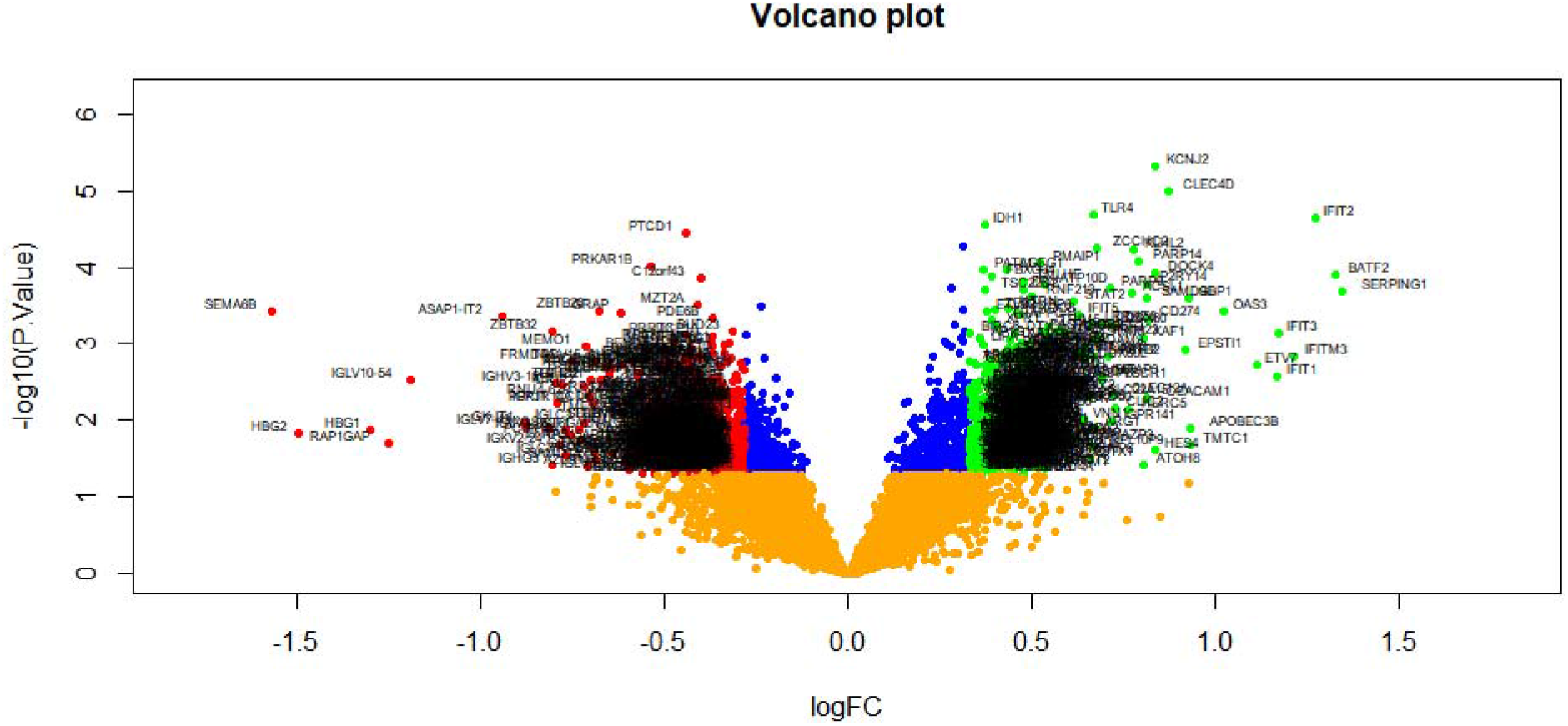
Volcano plot of differentially expressed genes. Genes with a significant change of more than two-fold were selected. Green dot represented up regulated significant genes and red dot represented down regulated significant genes.

**Fig. 2.**
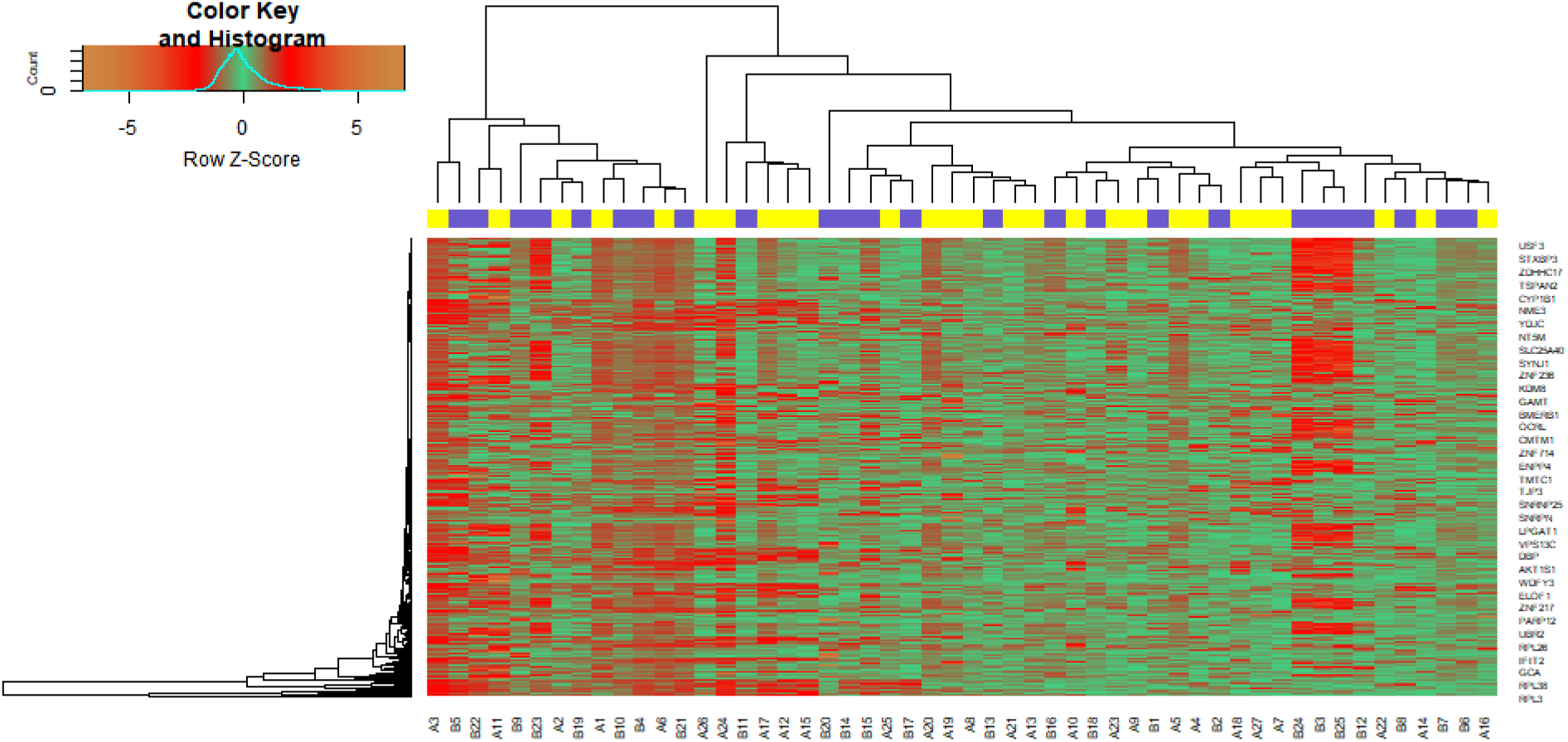
Heat map of differentially expressed genes. Legend on the top left indicate log fold change of genes. (A1 – A76 = CAD samples; B1 – B62 = normal control samples)

**Table 1.**
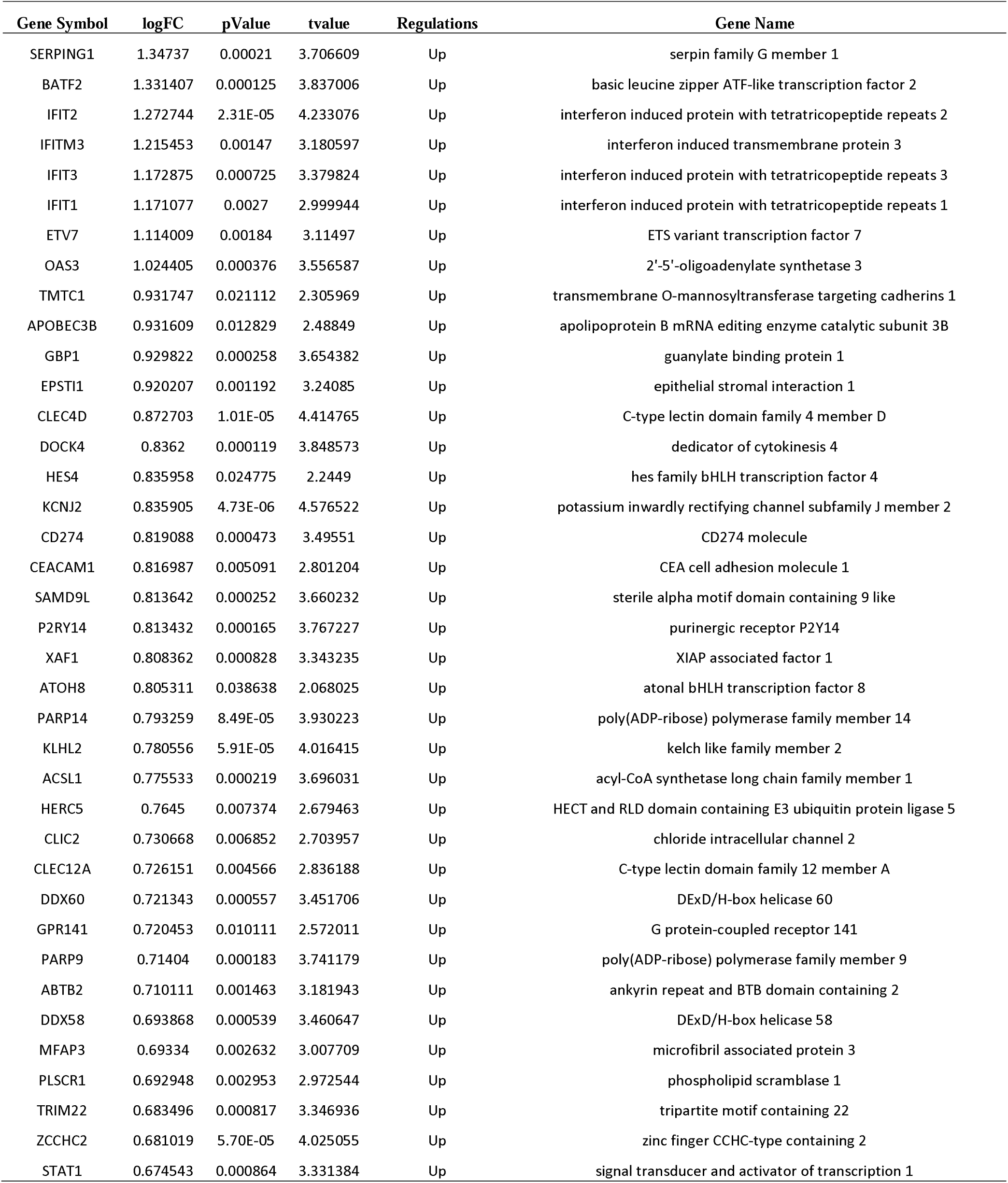
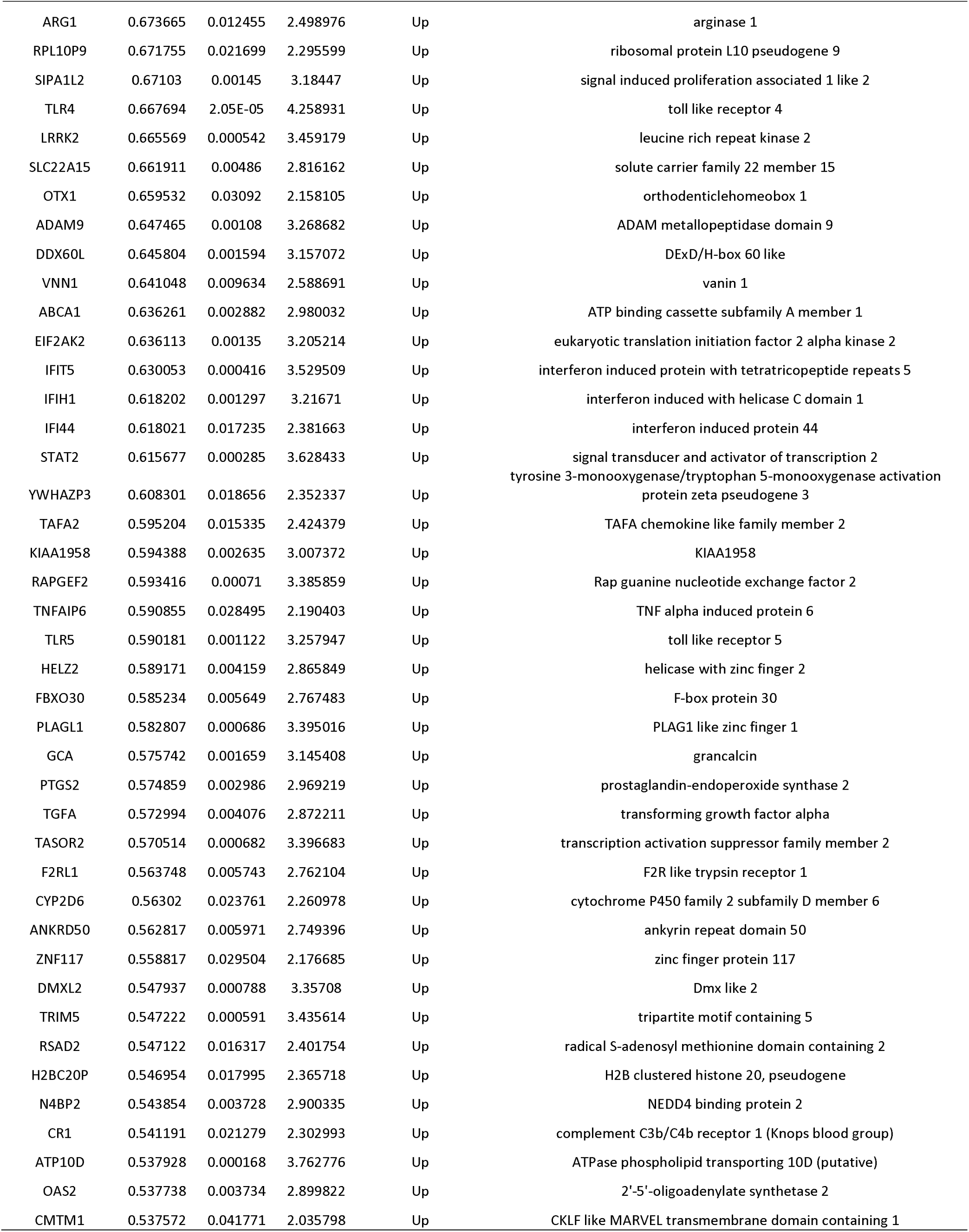
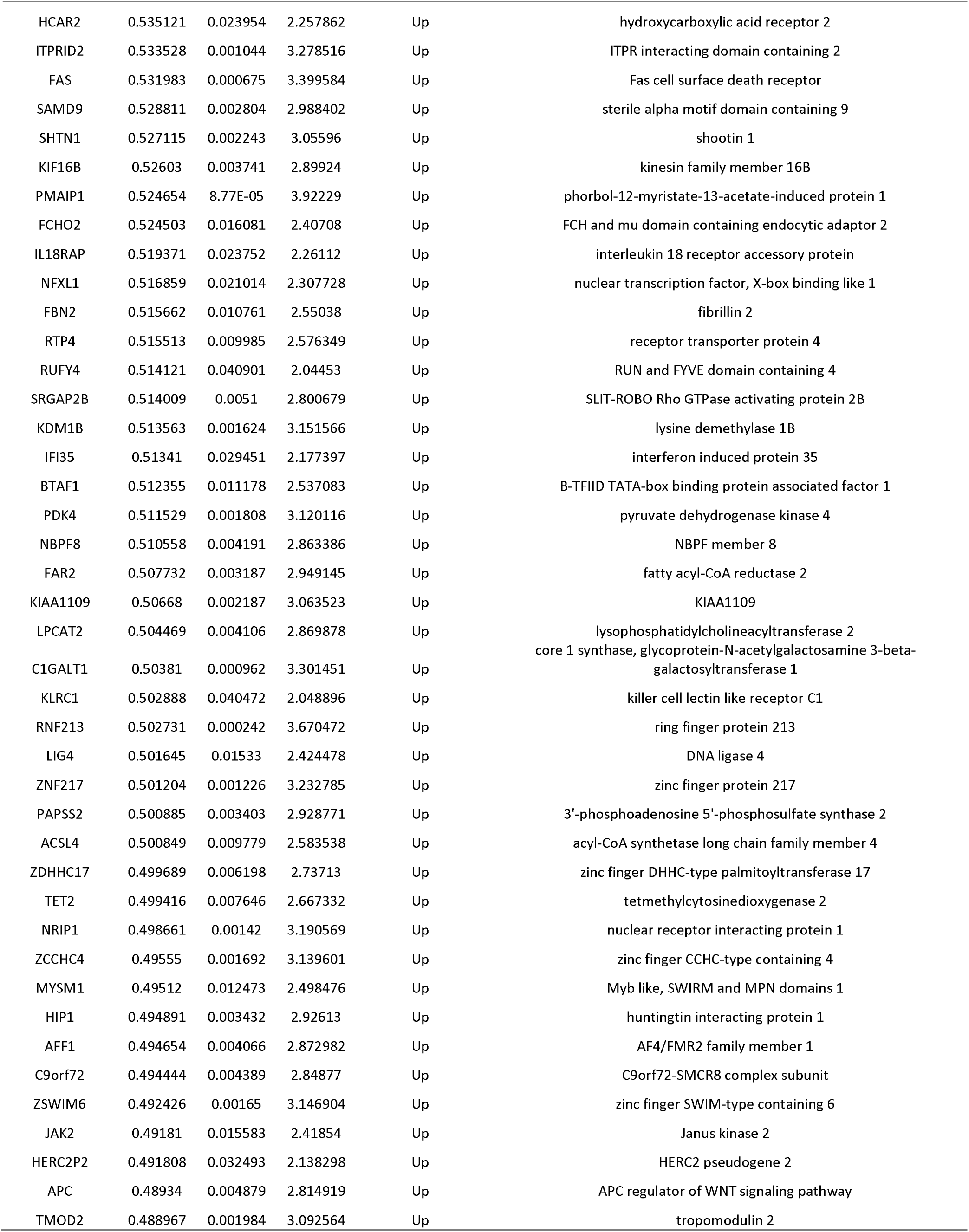
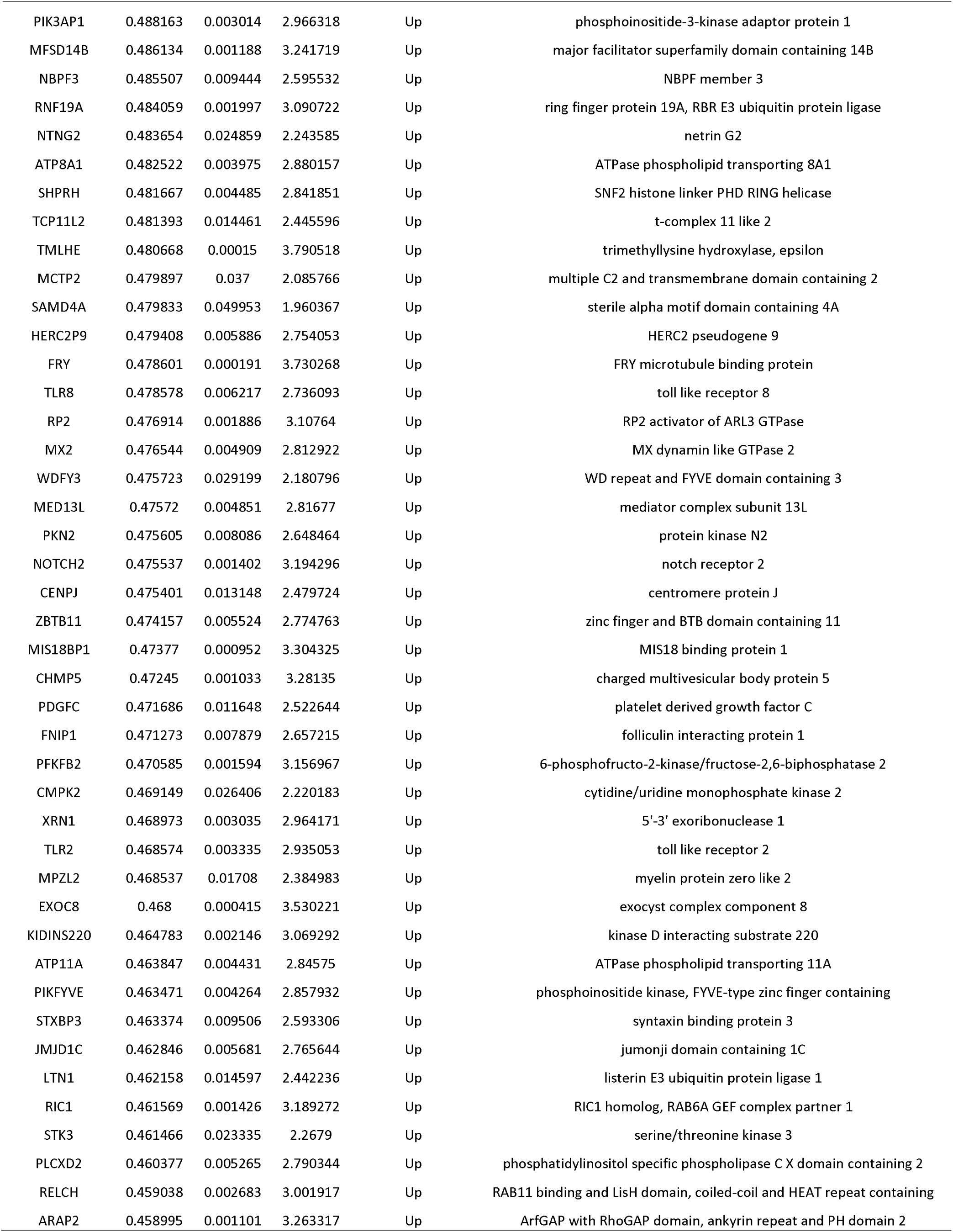
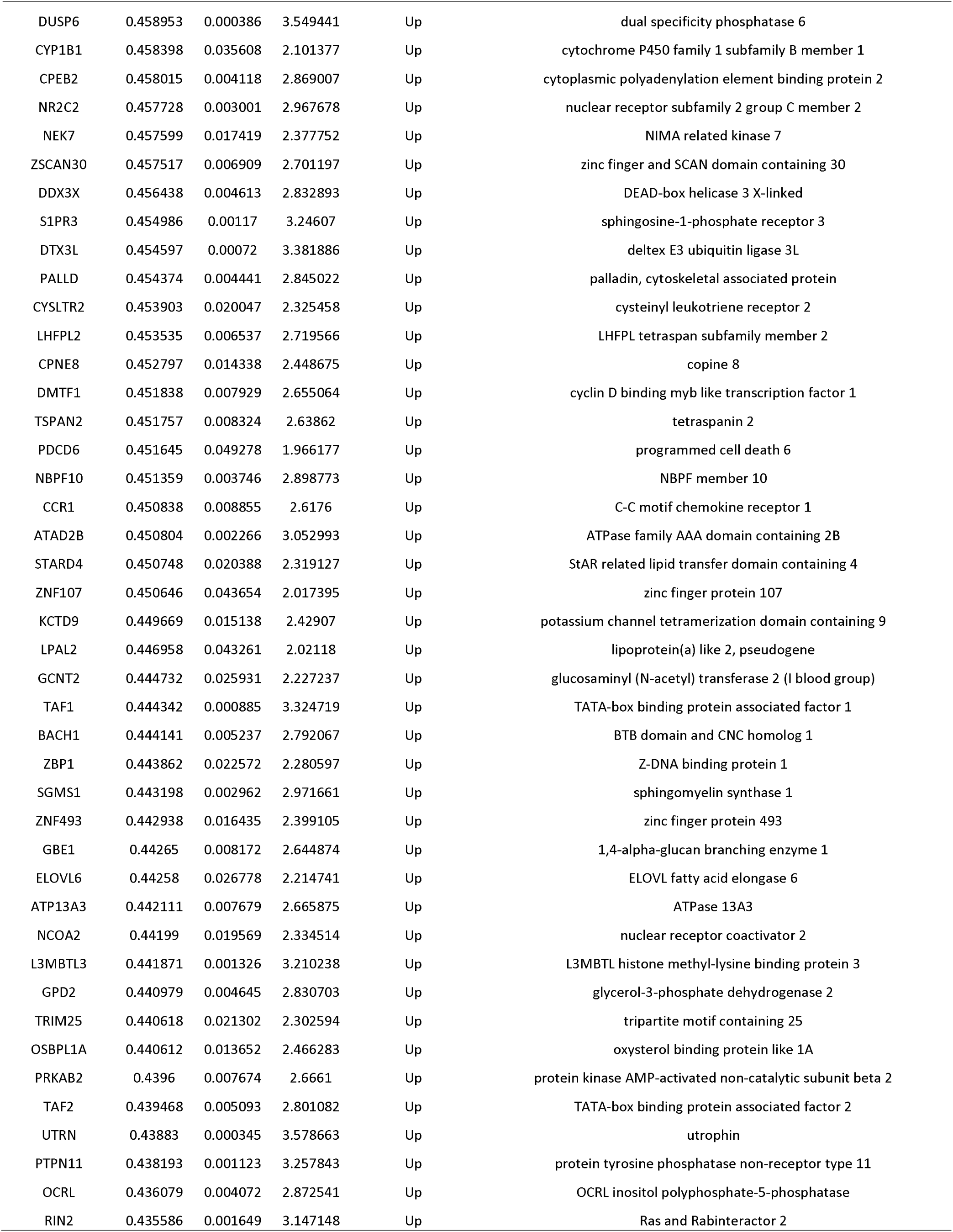
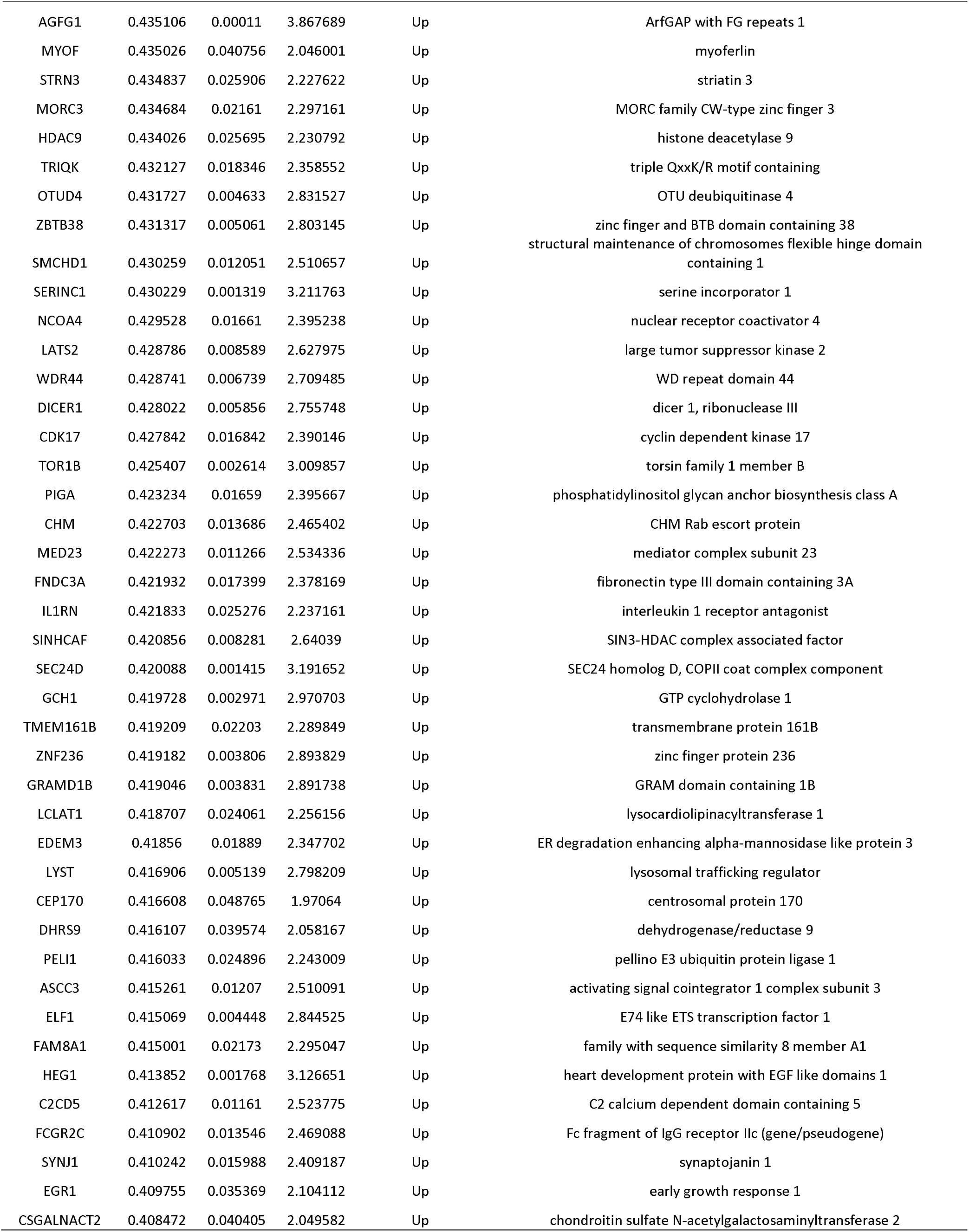
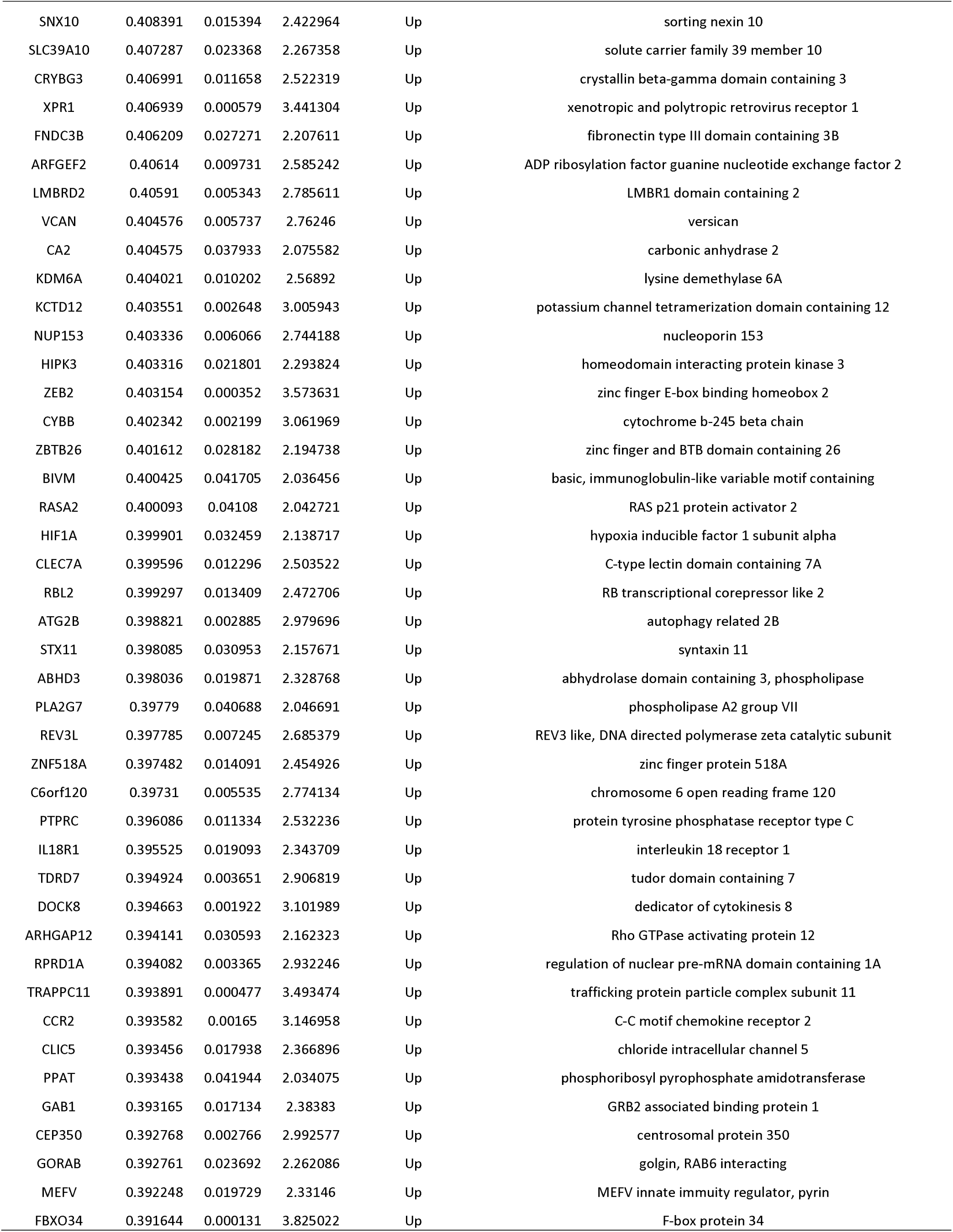
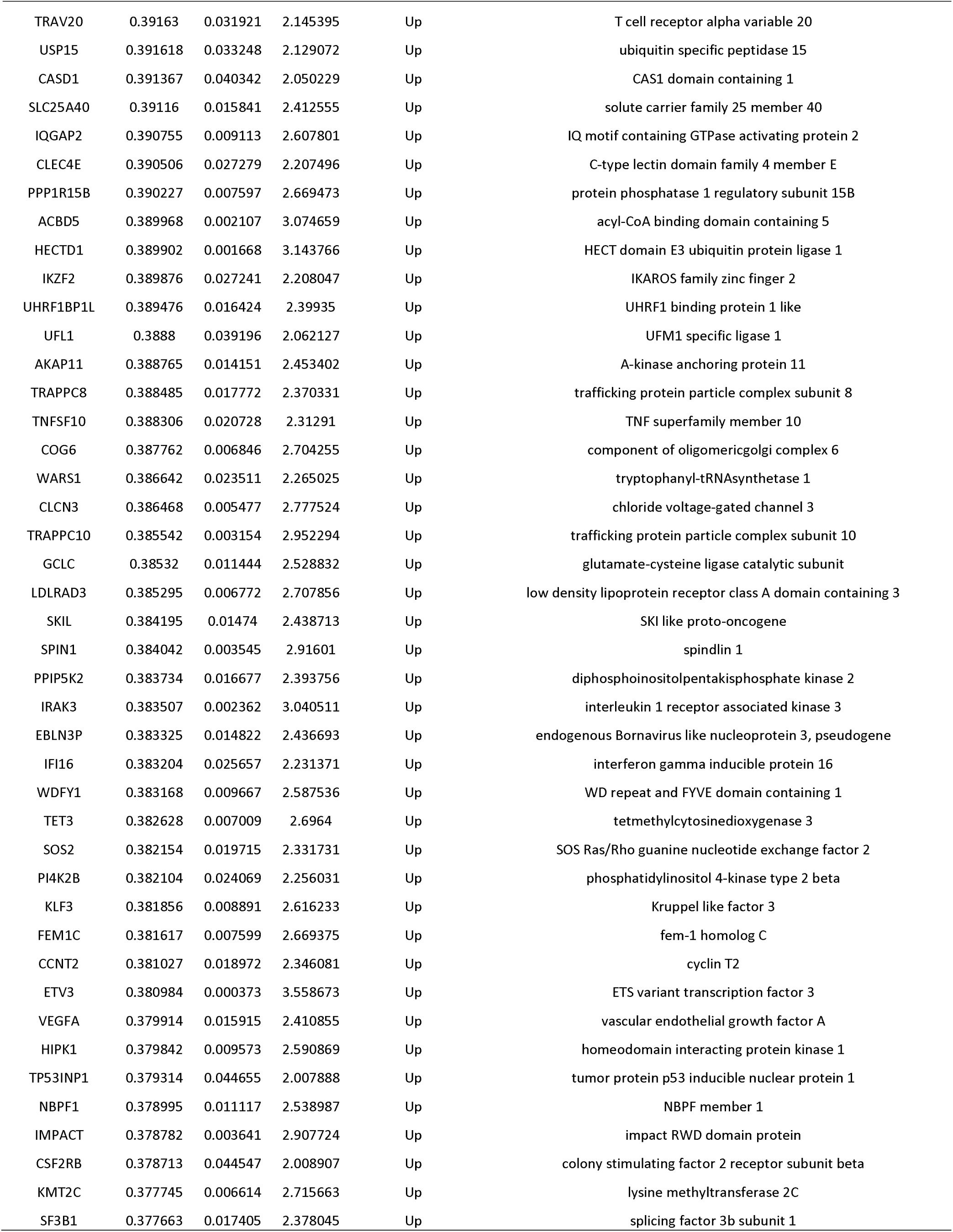
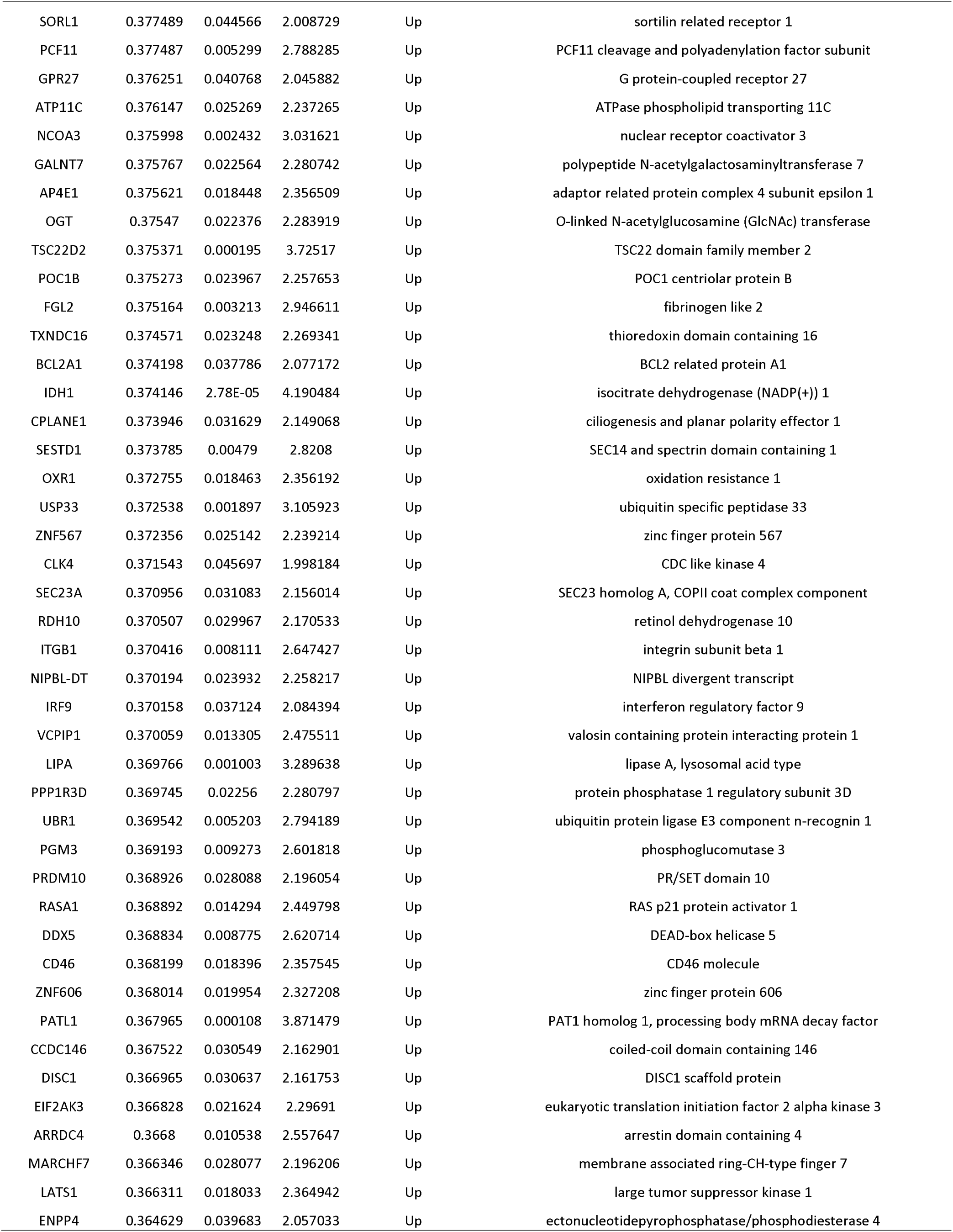
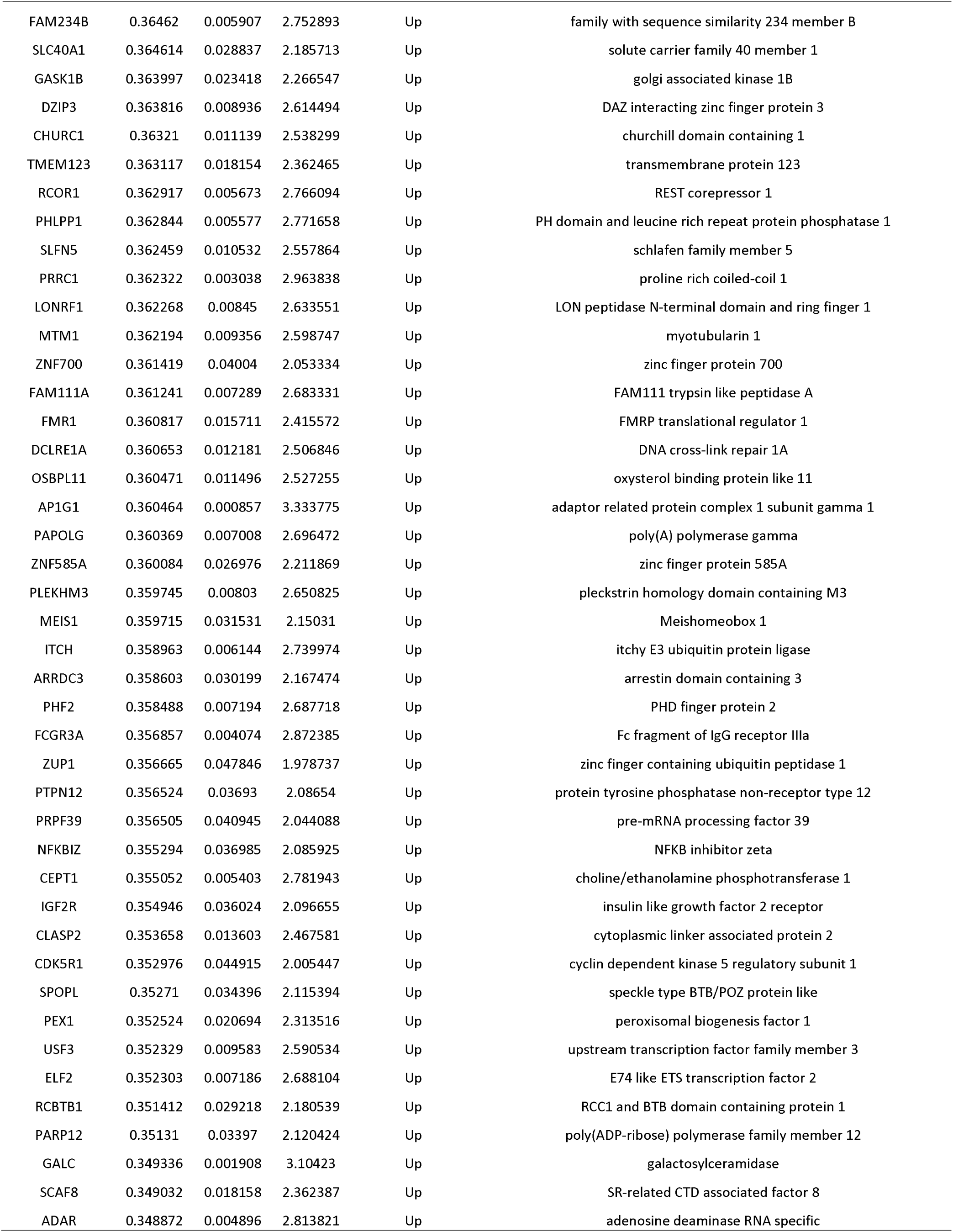
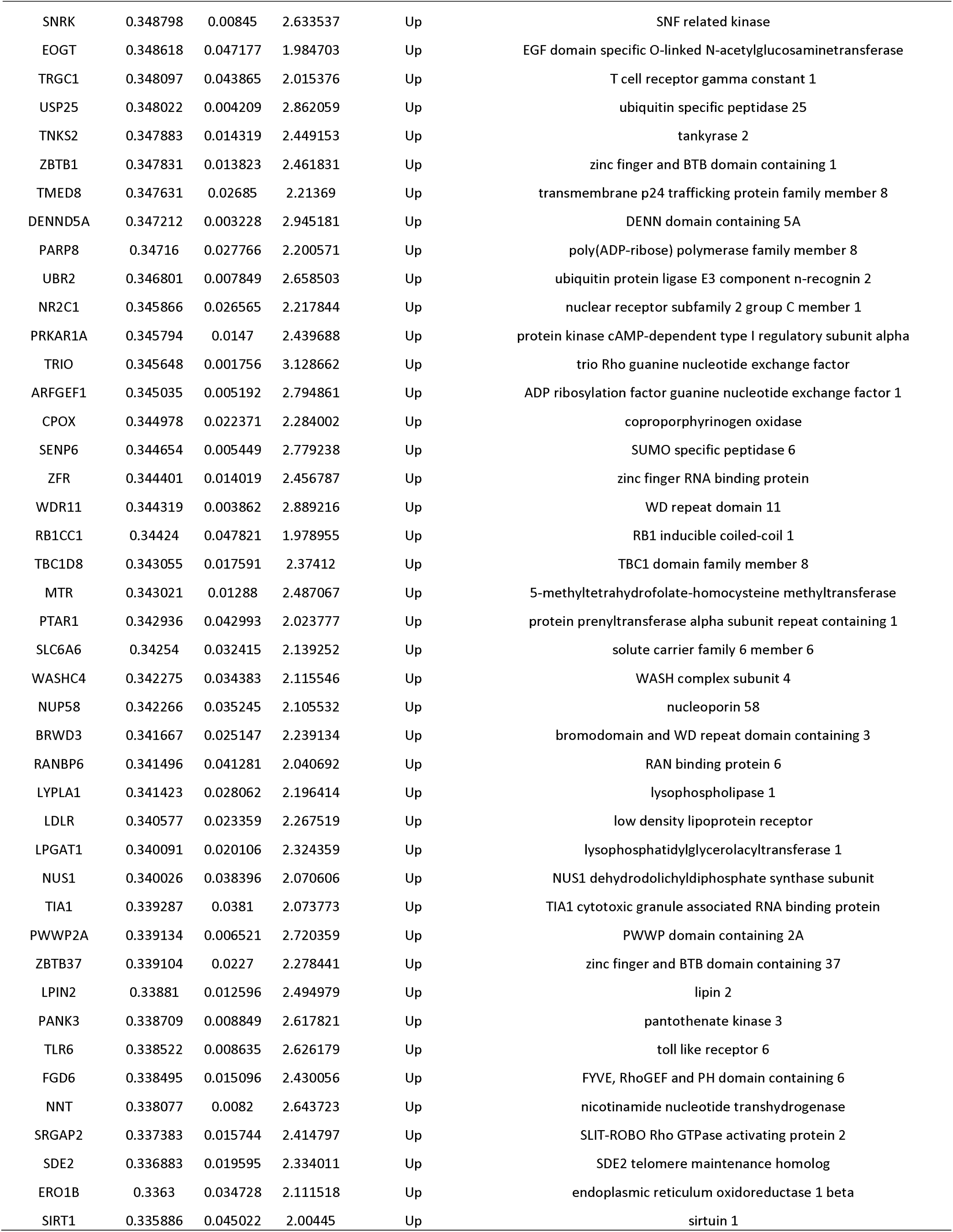
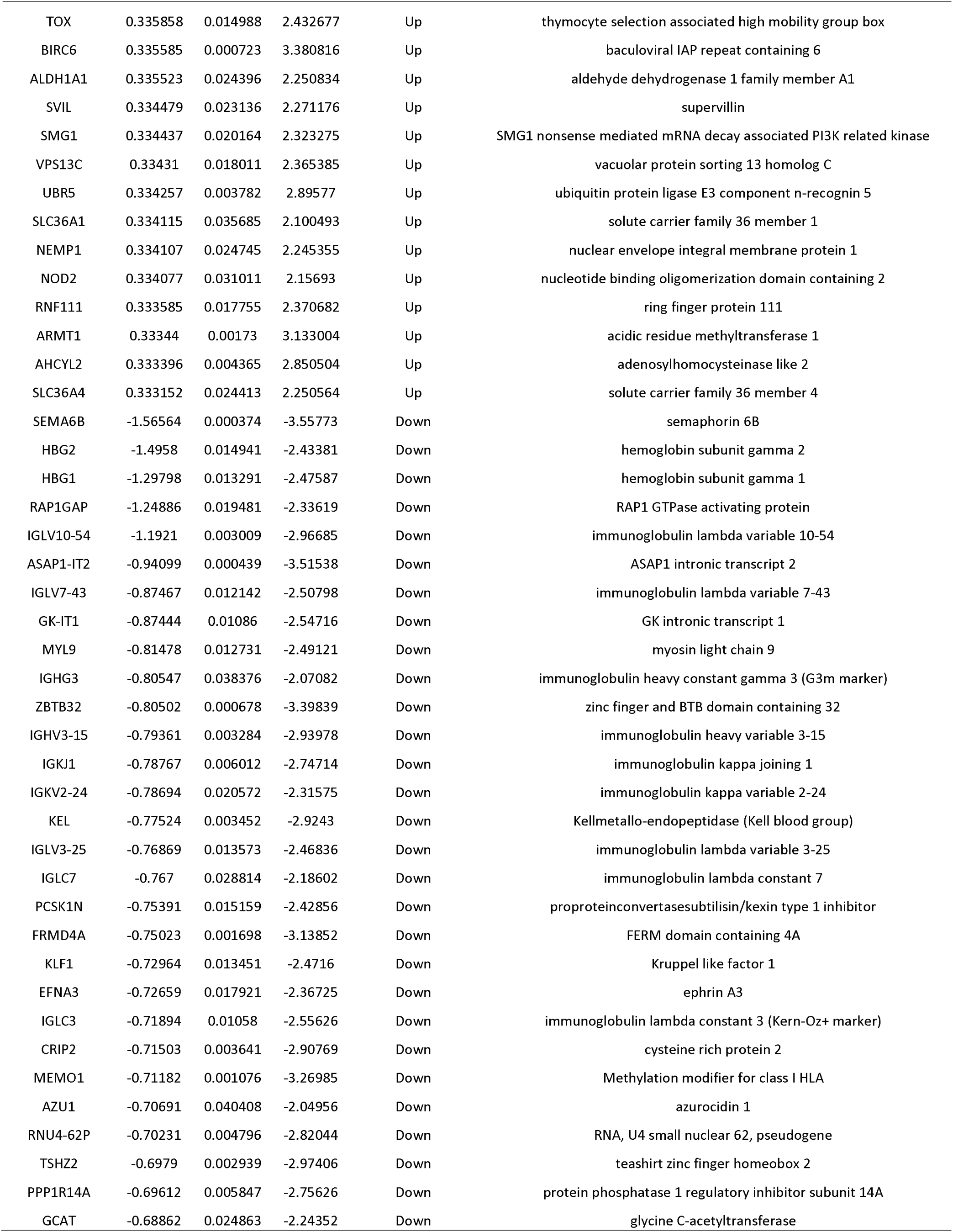
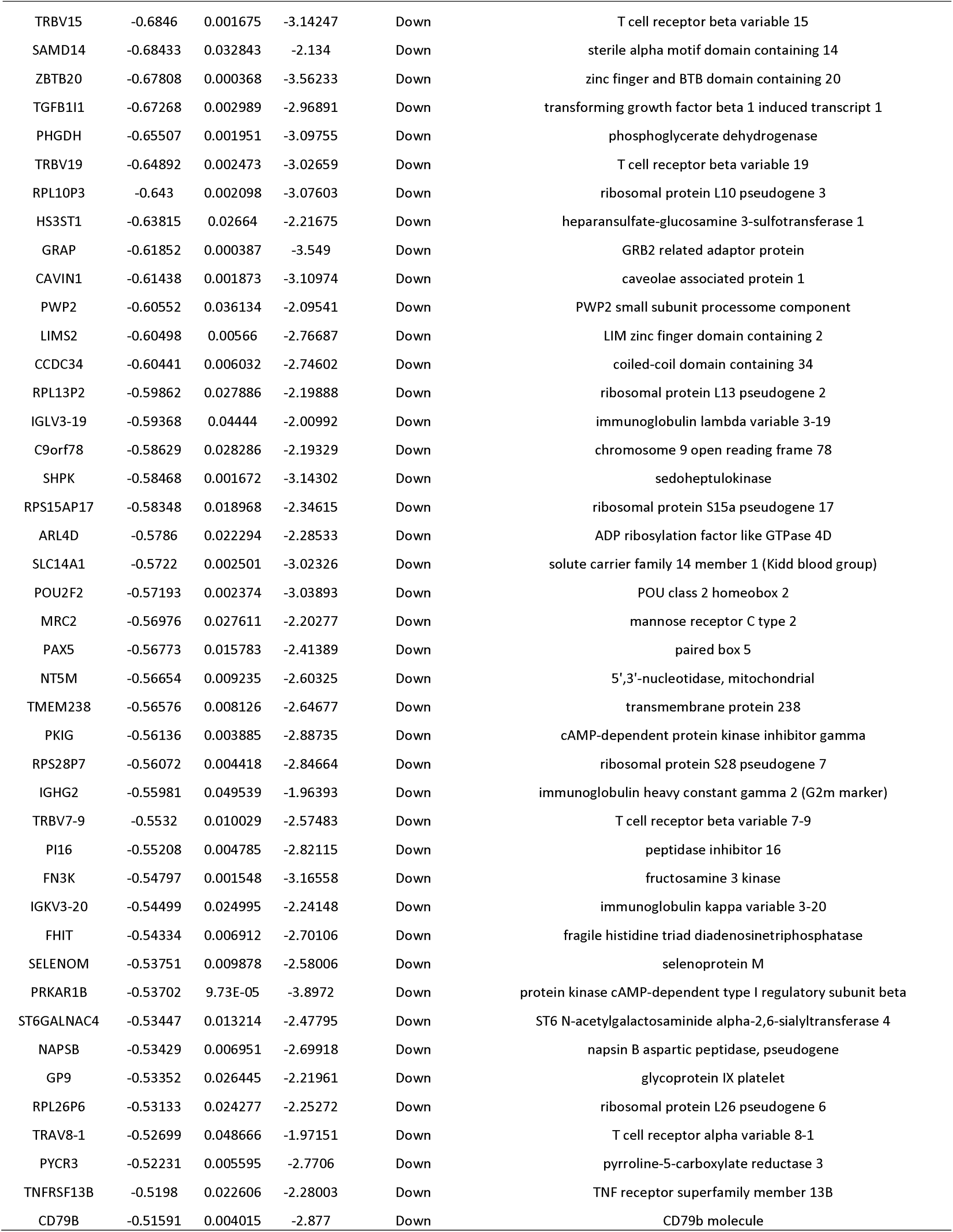
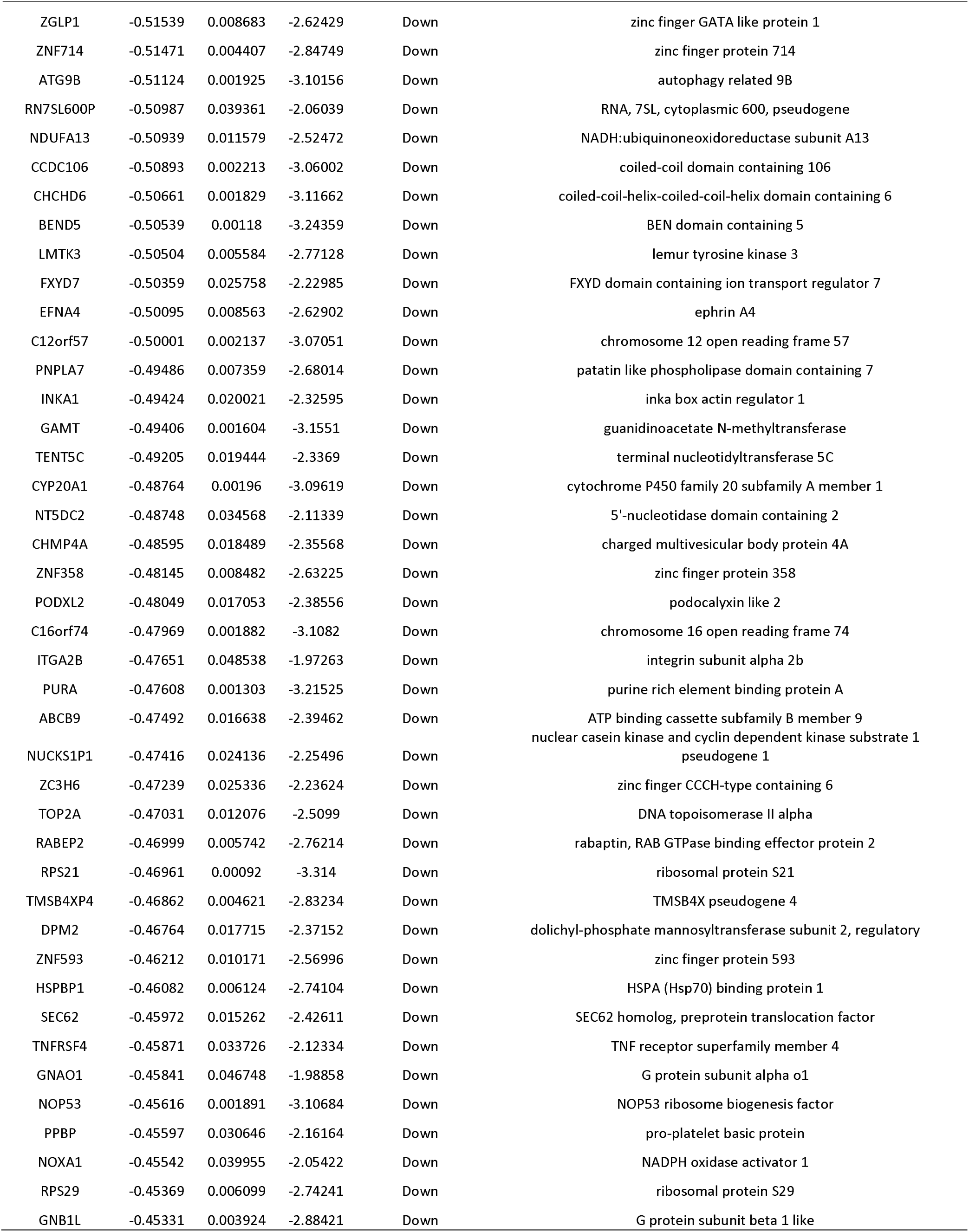
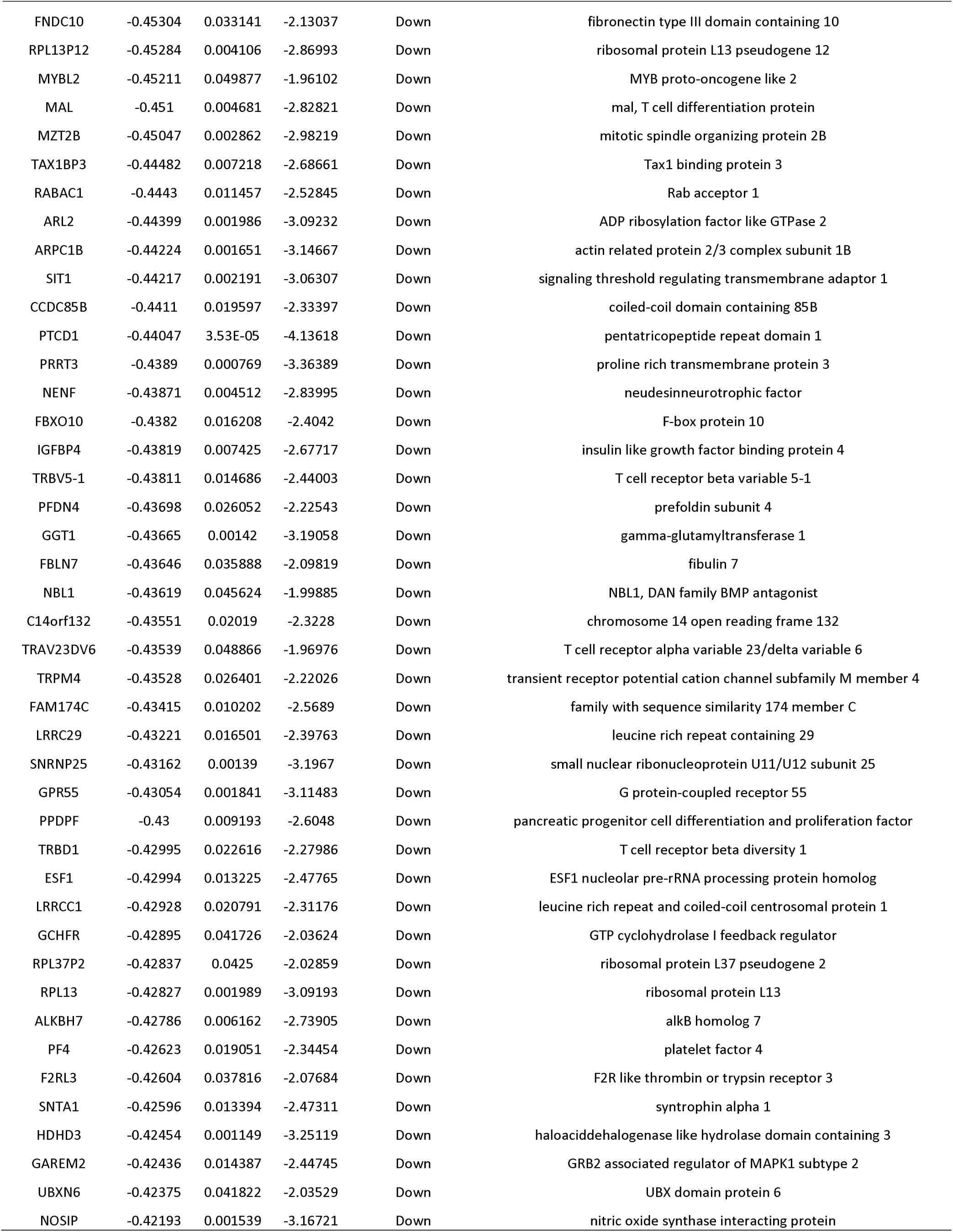
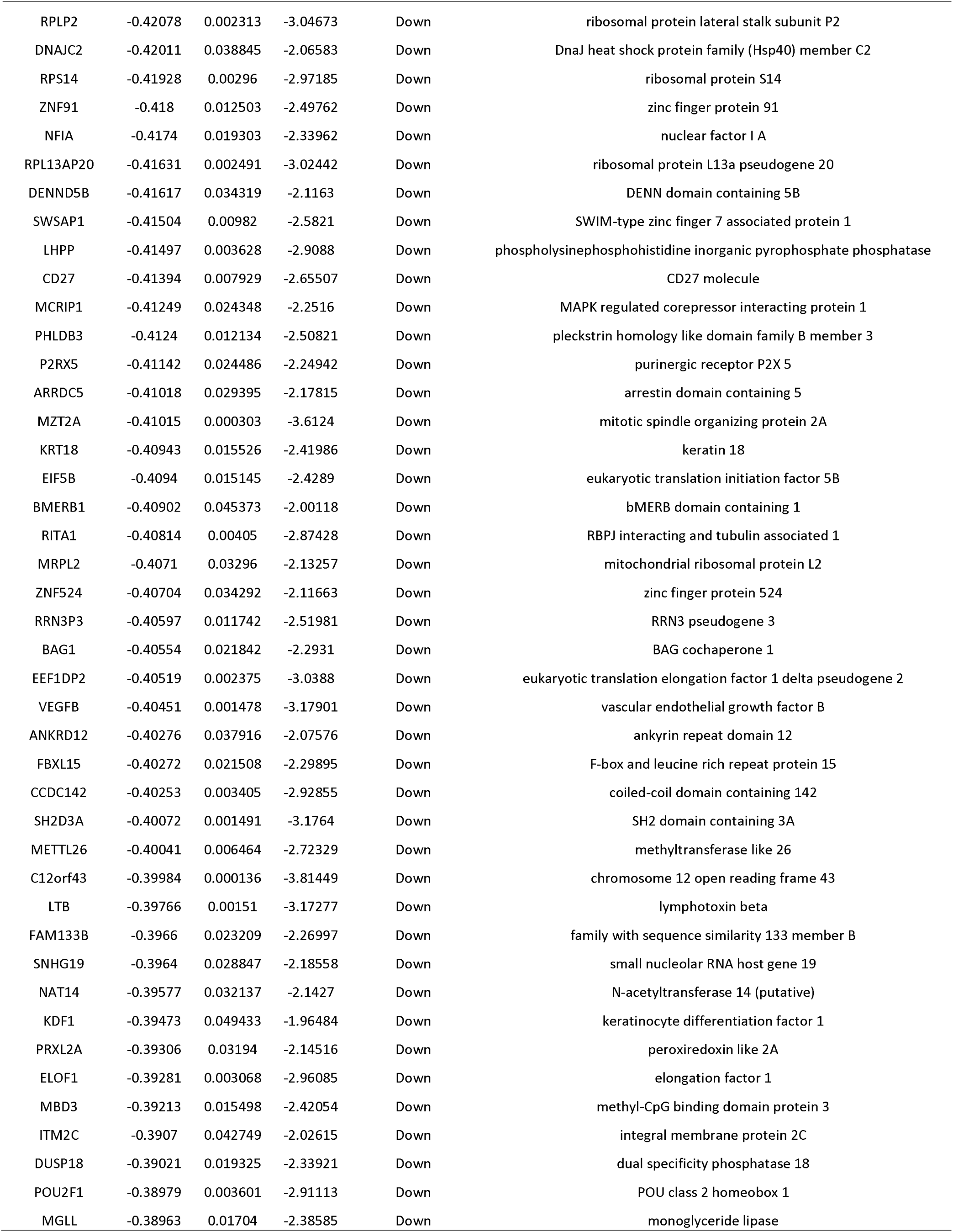
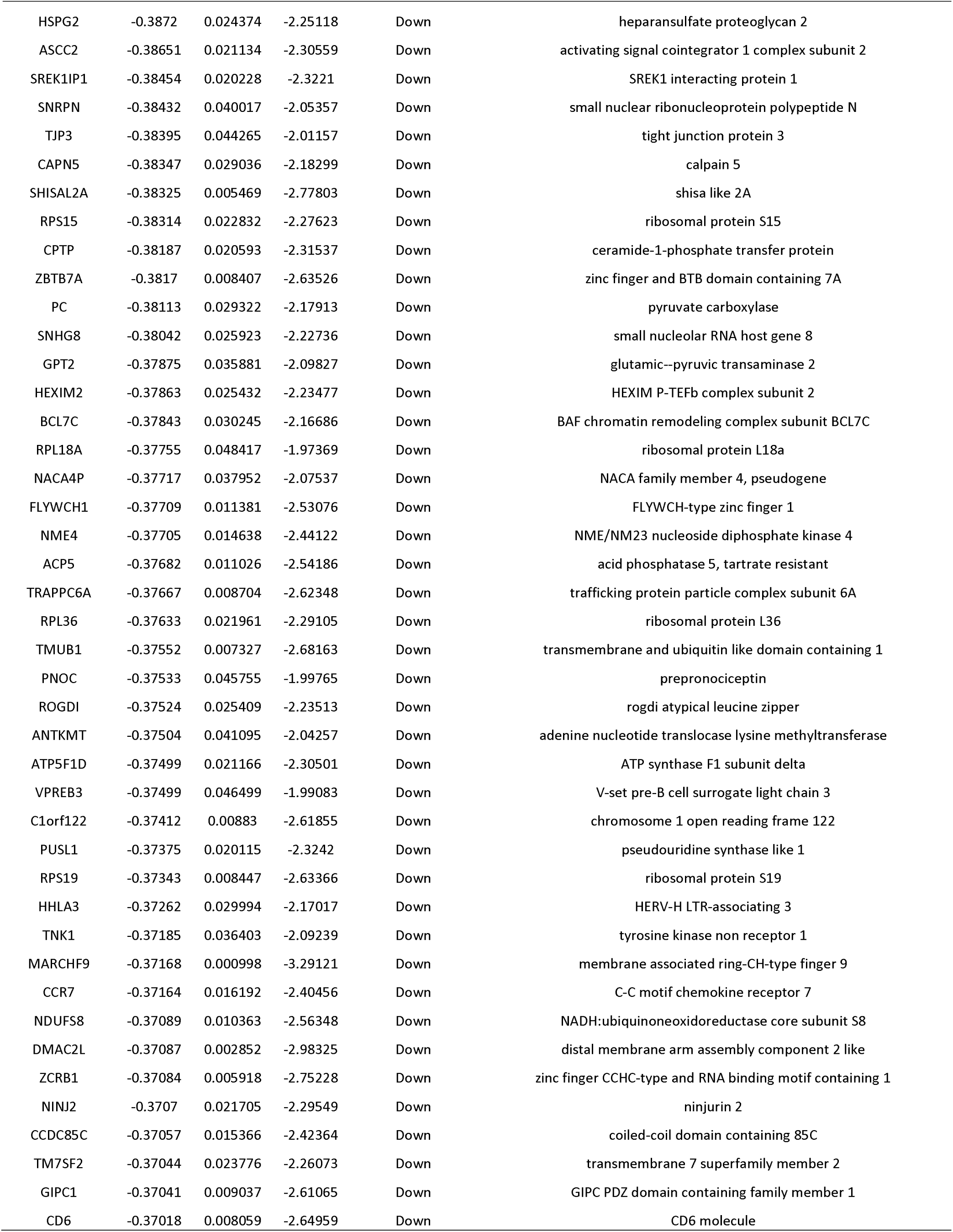
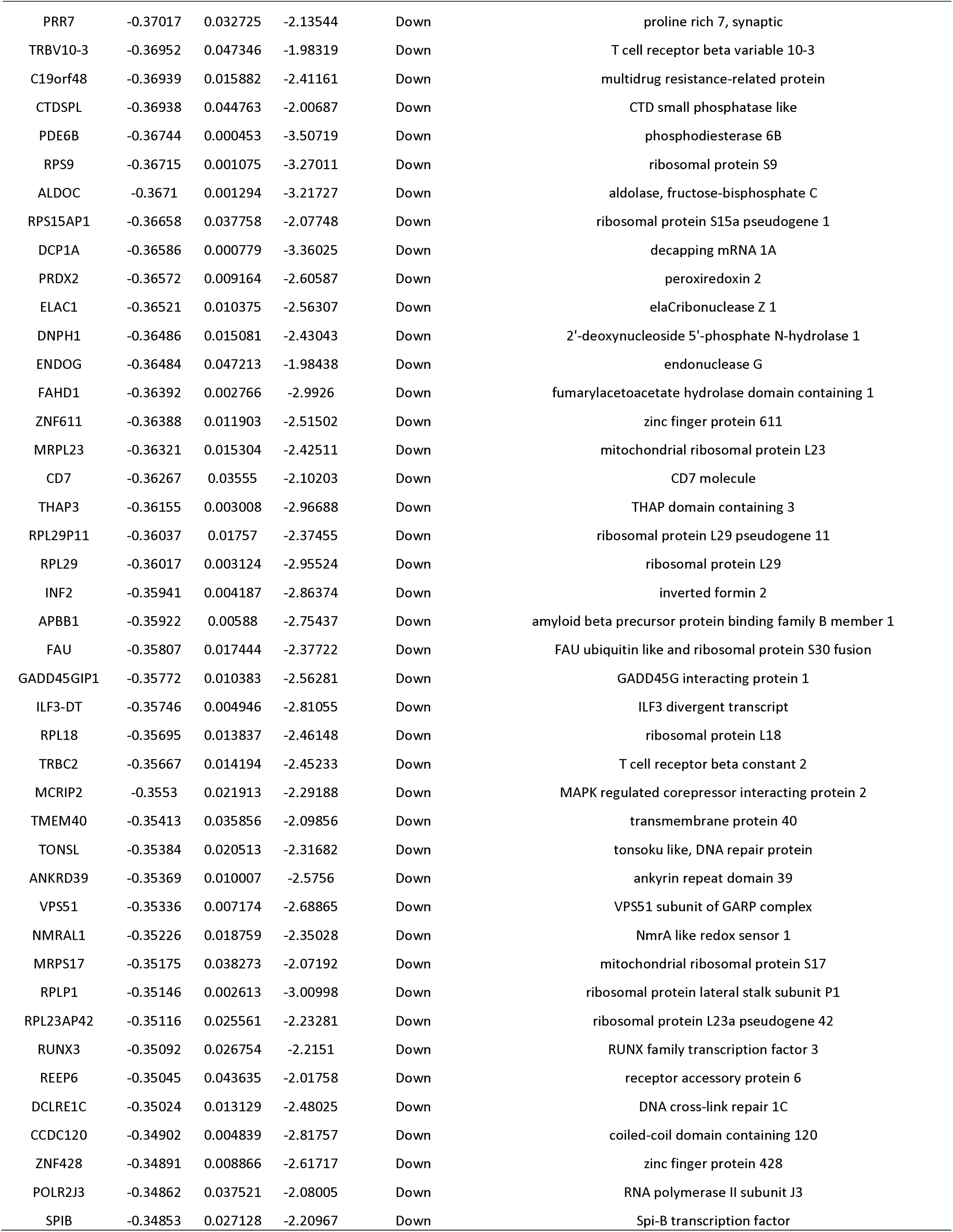
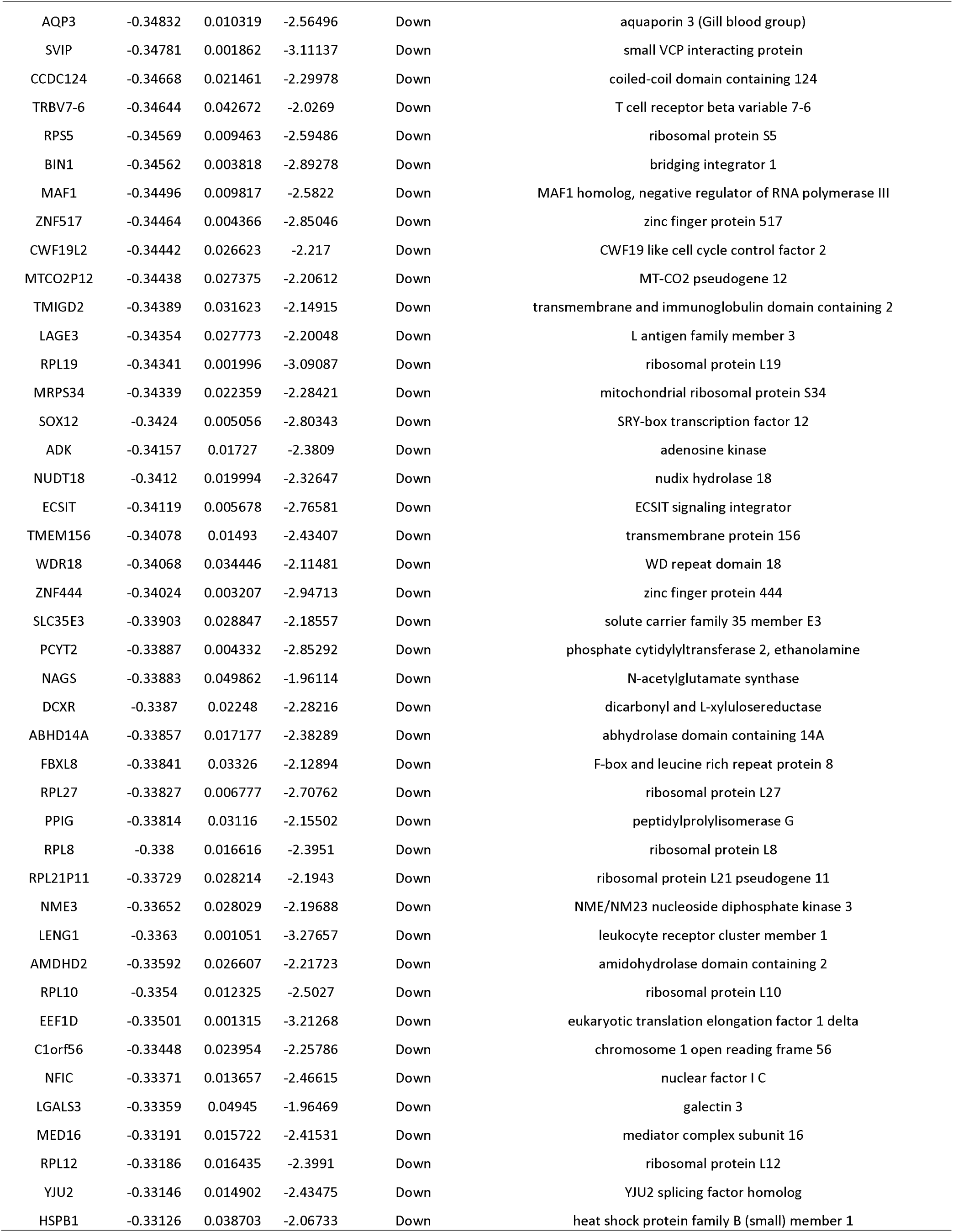
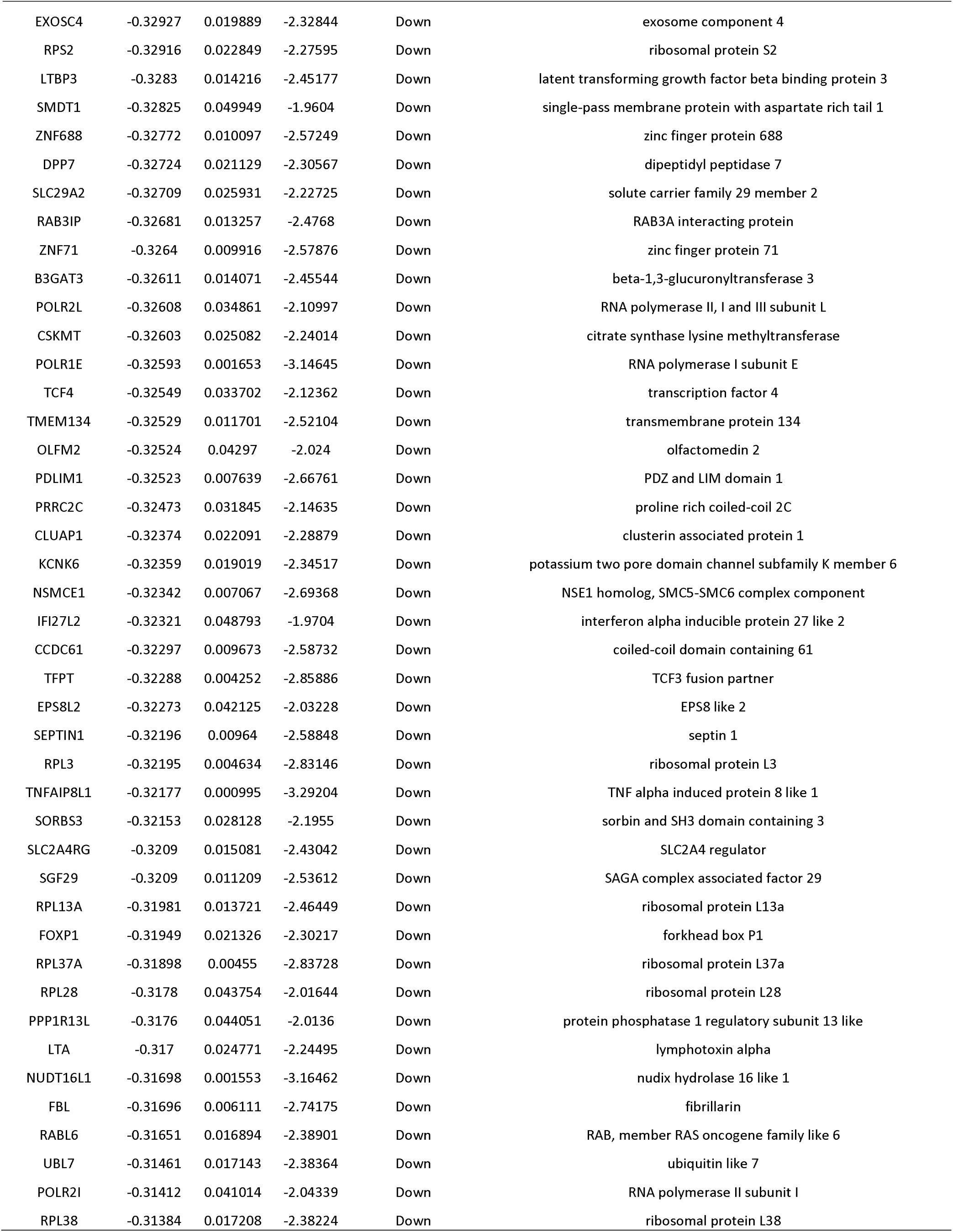
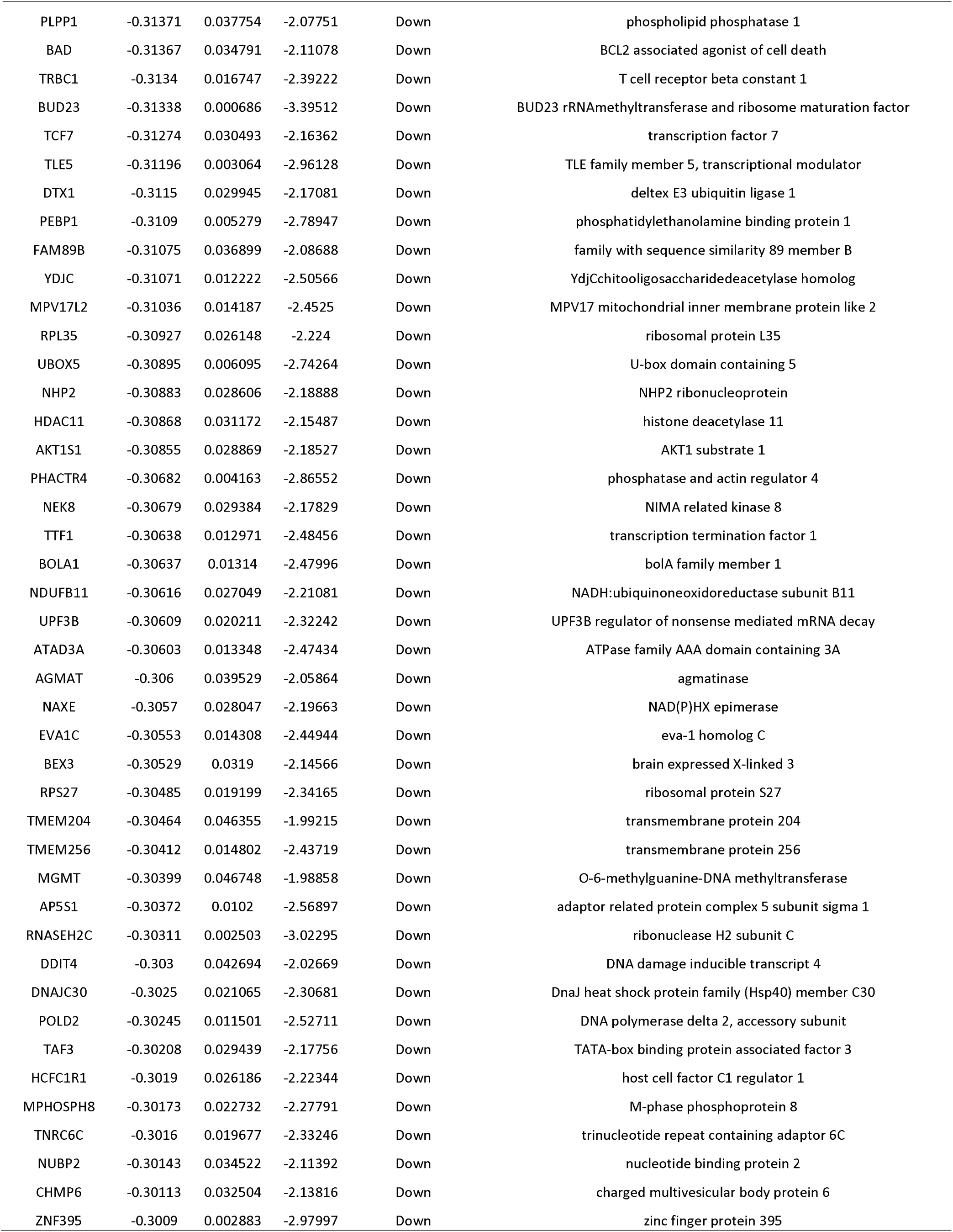
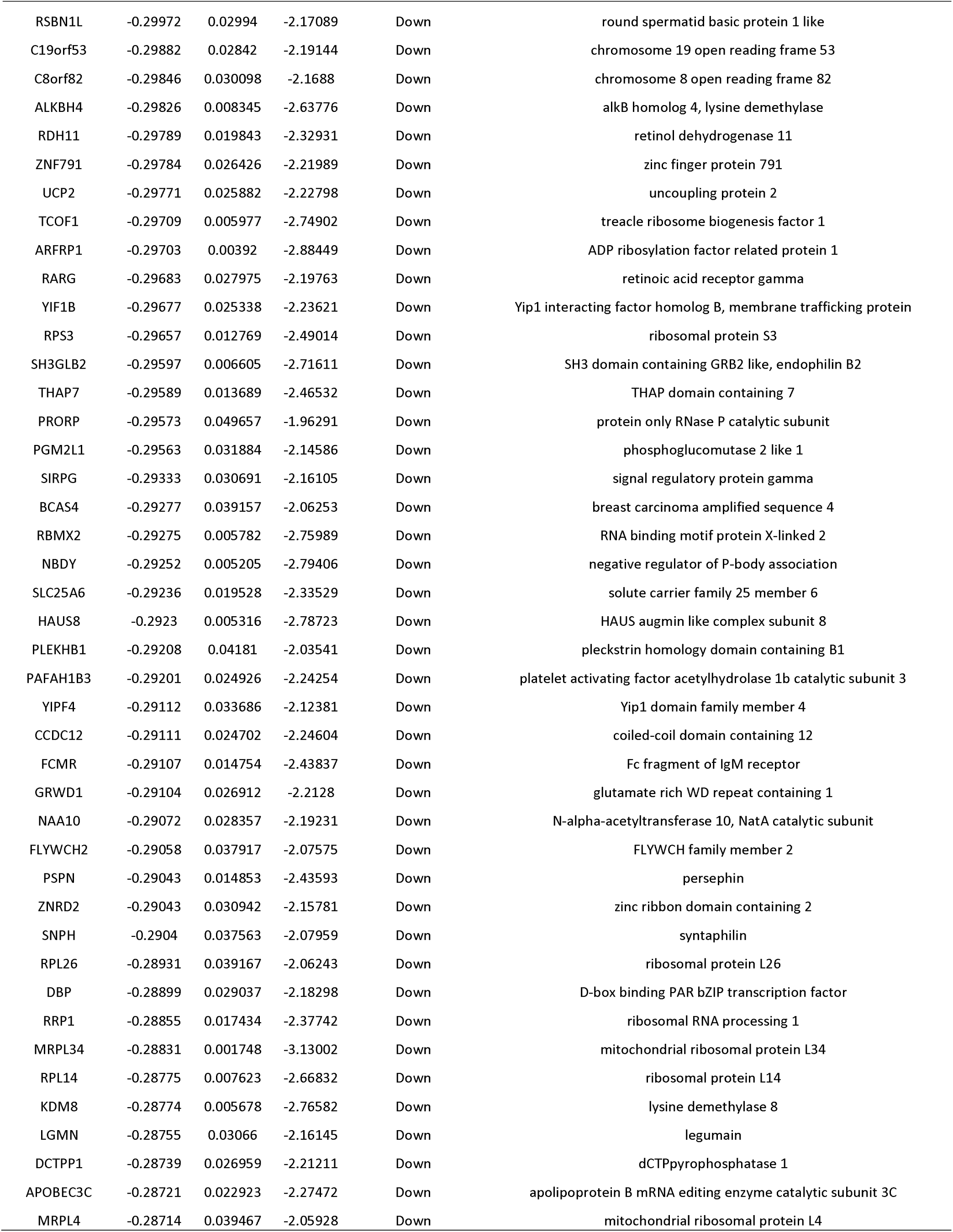
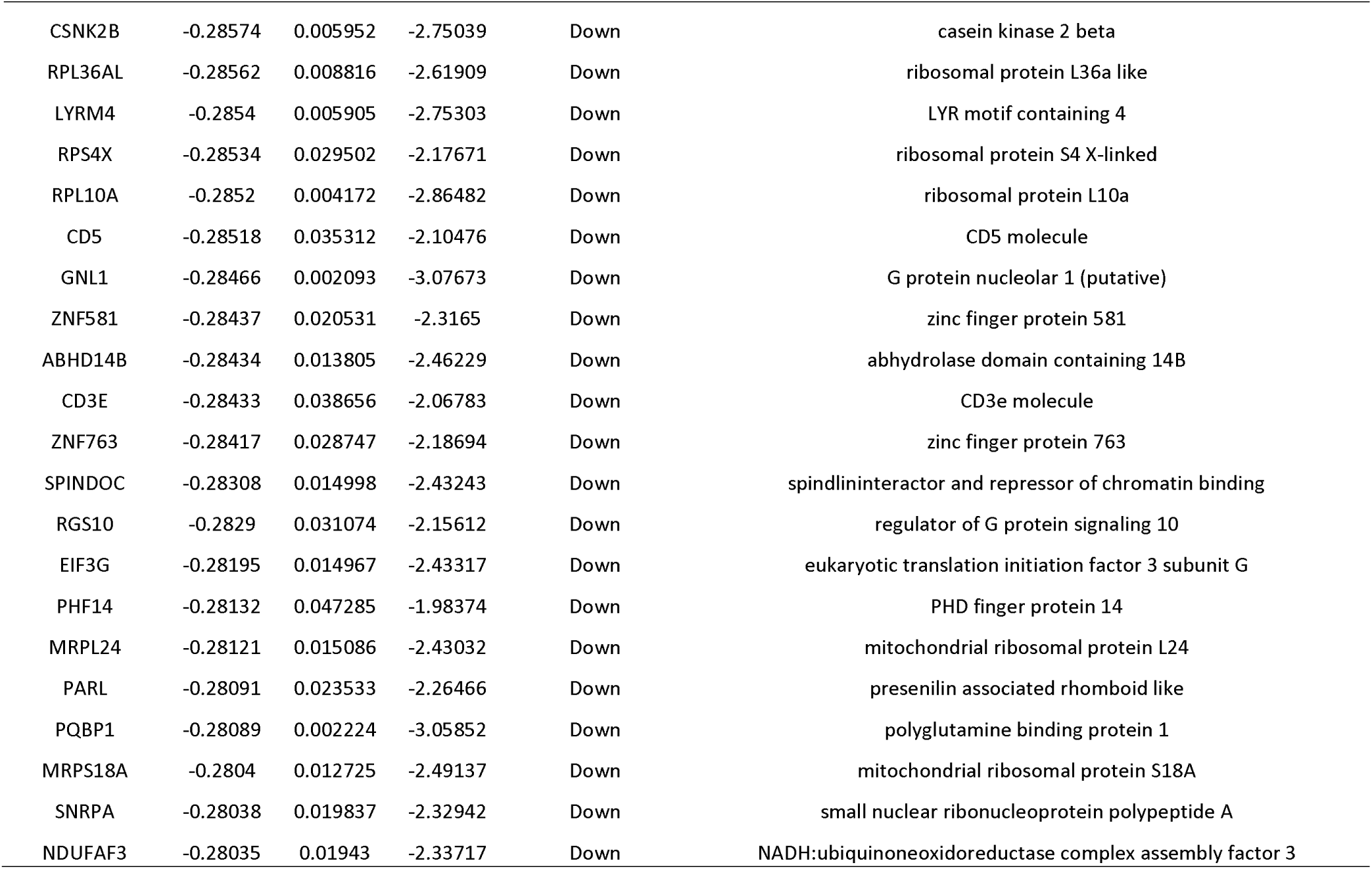
The statistical metrics for key differentially expressed genes (DEGs)

### GO and pathway enrichment analyses of DEGs

GO and REACTOME enrichment analysis of DEGs between CAD and normal controls groups were performed (Table 2 and Table 3). In GO BP analysis, up regulated genes were primarily rich in cellular response to stimulus and biological regulation, while down regulated genes were primarily rich in biosynthetic process and organonitrogen compound metabolic process. In GO CC analysis, up regulated genes were mainly rich in endomembrane system and intracellular anatomical structure, while down regulated genes were primarily rich in intracellular organelle lumen and cytoplasm. In GO MF analysis, up regulated genes were mainly rich in ion binding and enzyme binding, while down regulated genes were mainly rich in structural molecule activity and nucleic acid binding. In REACTOME pathway enrichment analysis, up regulated genes were rich in immune system and cytokine signaling in immune system, while down regulated genes were rich in eukaryotic translation elongation and disease.

**Table 2.**
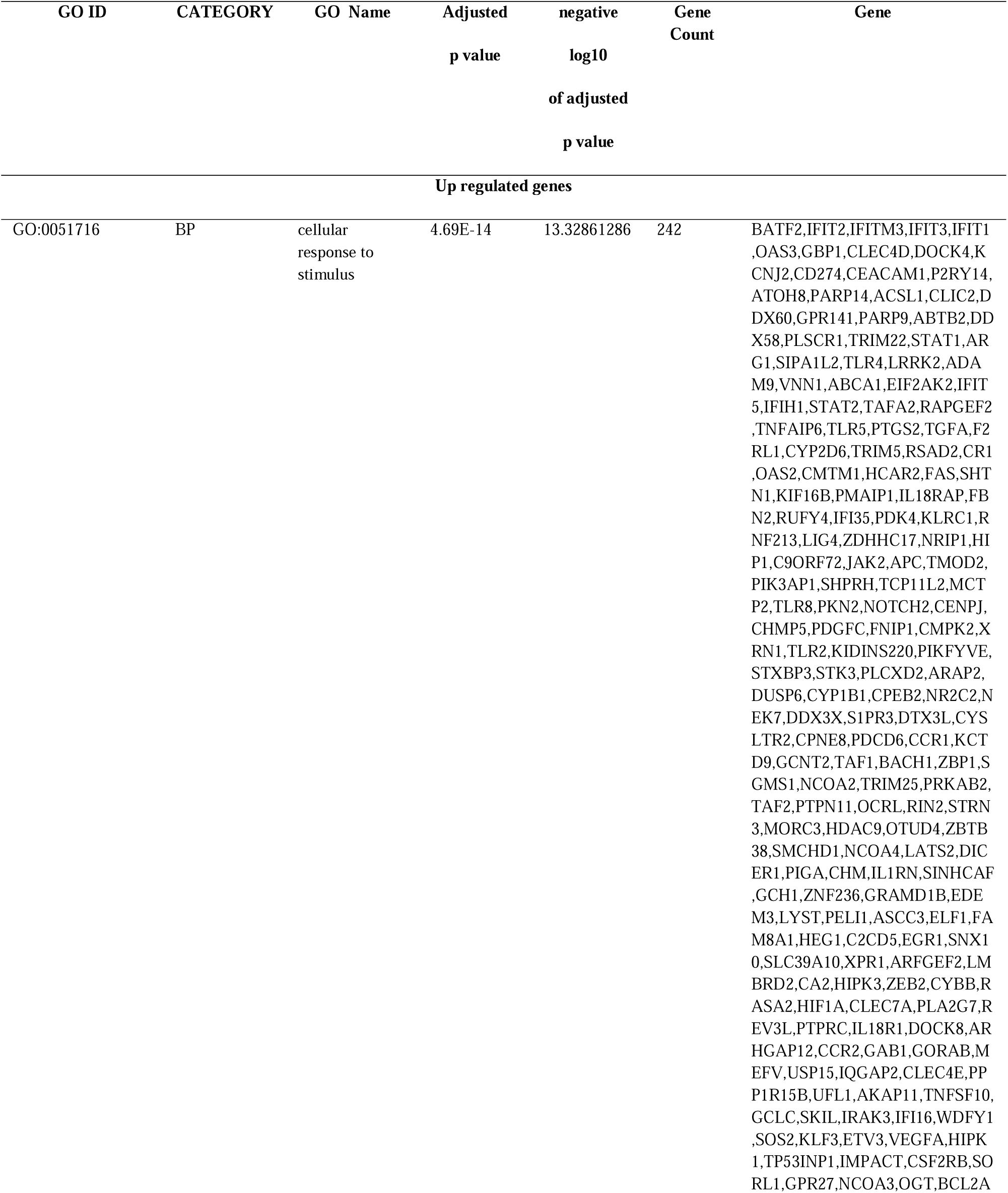
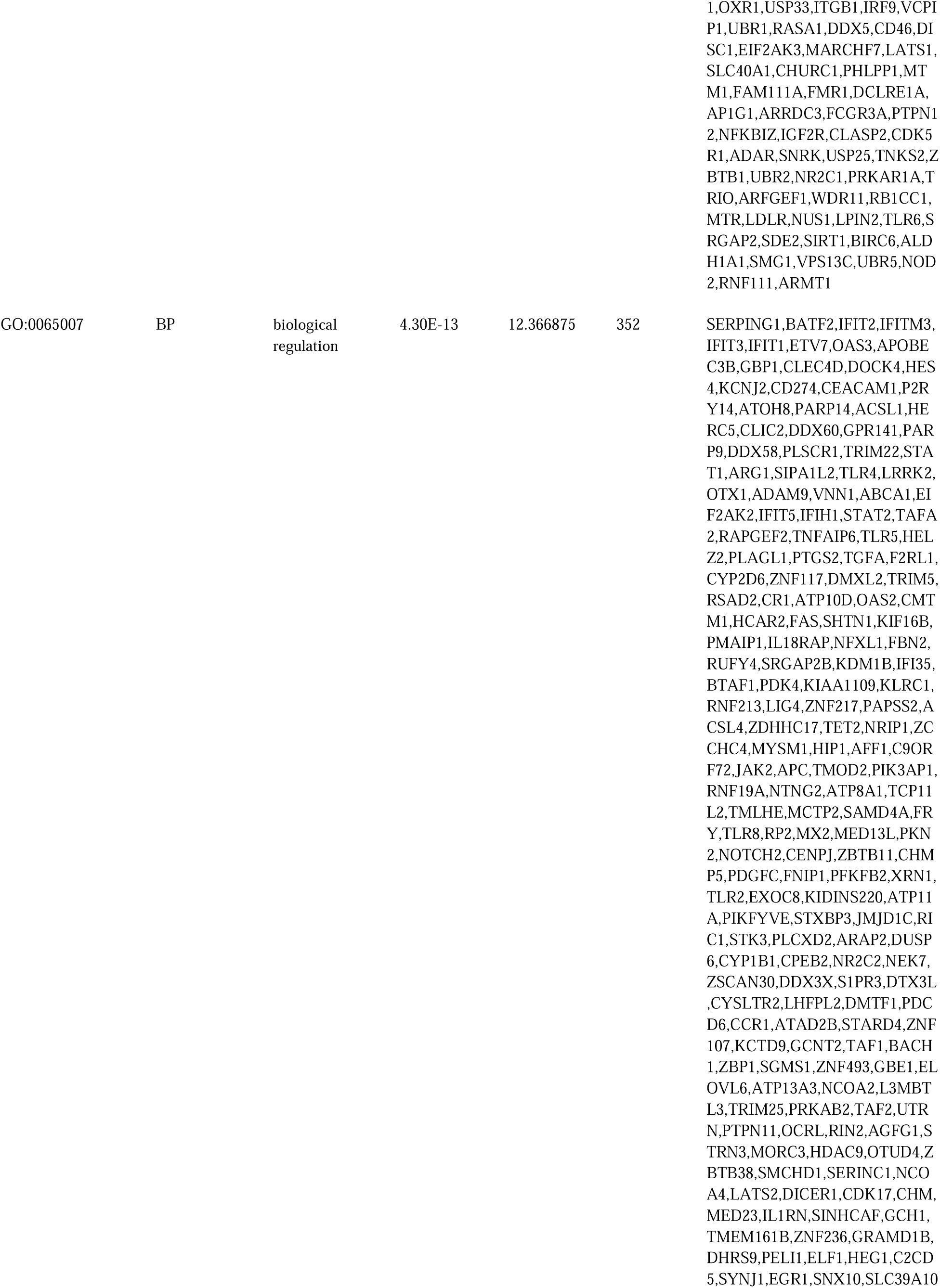
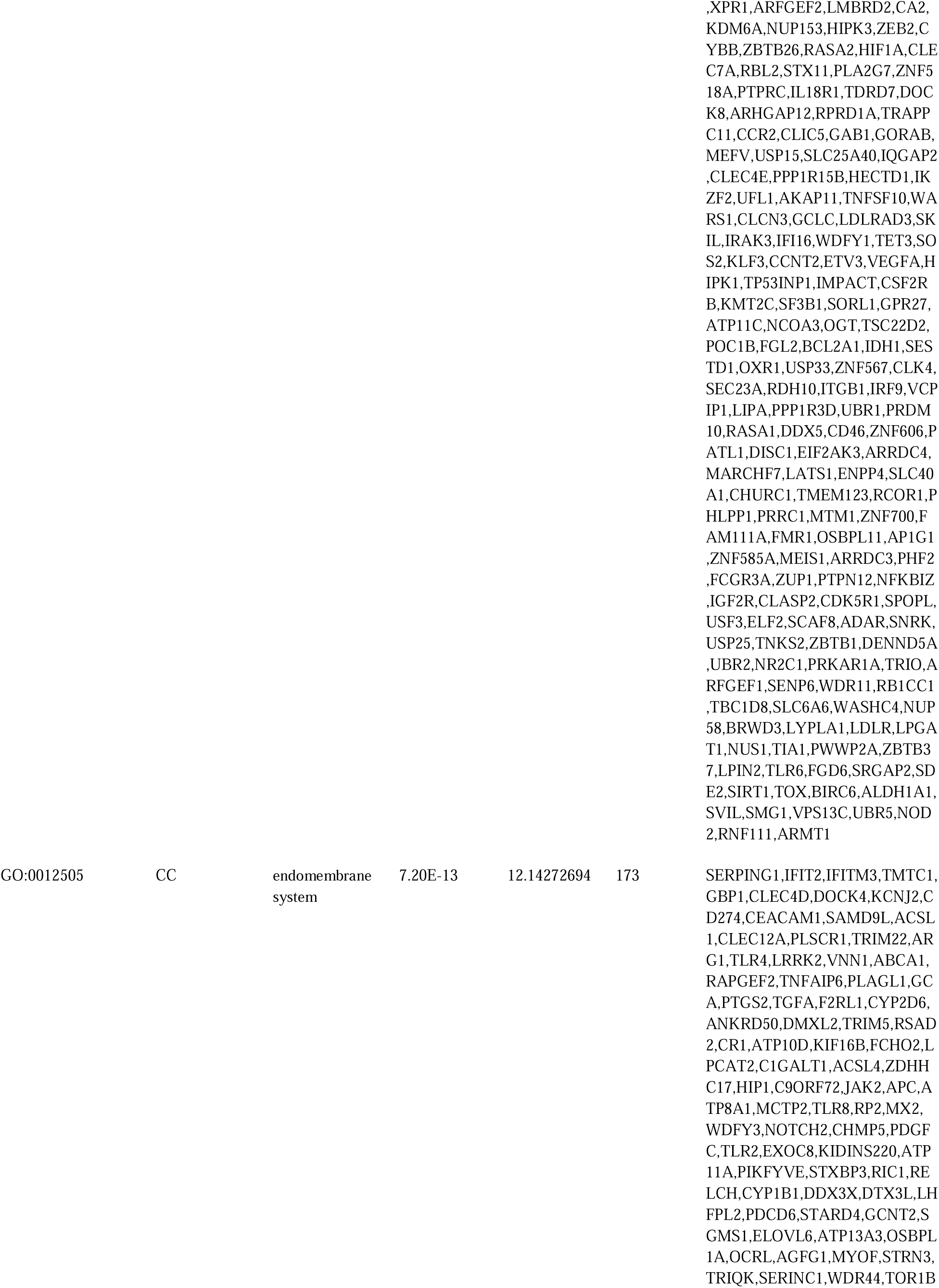
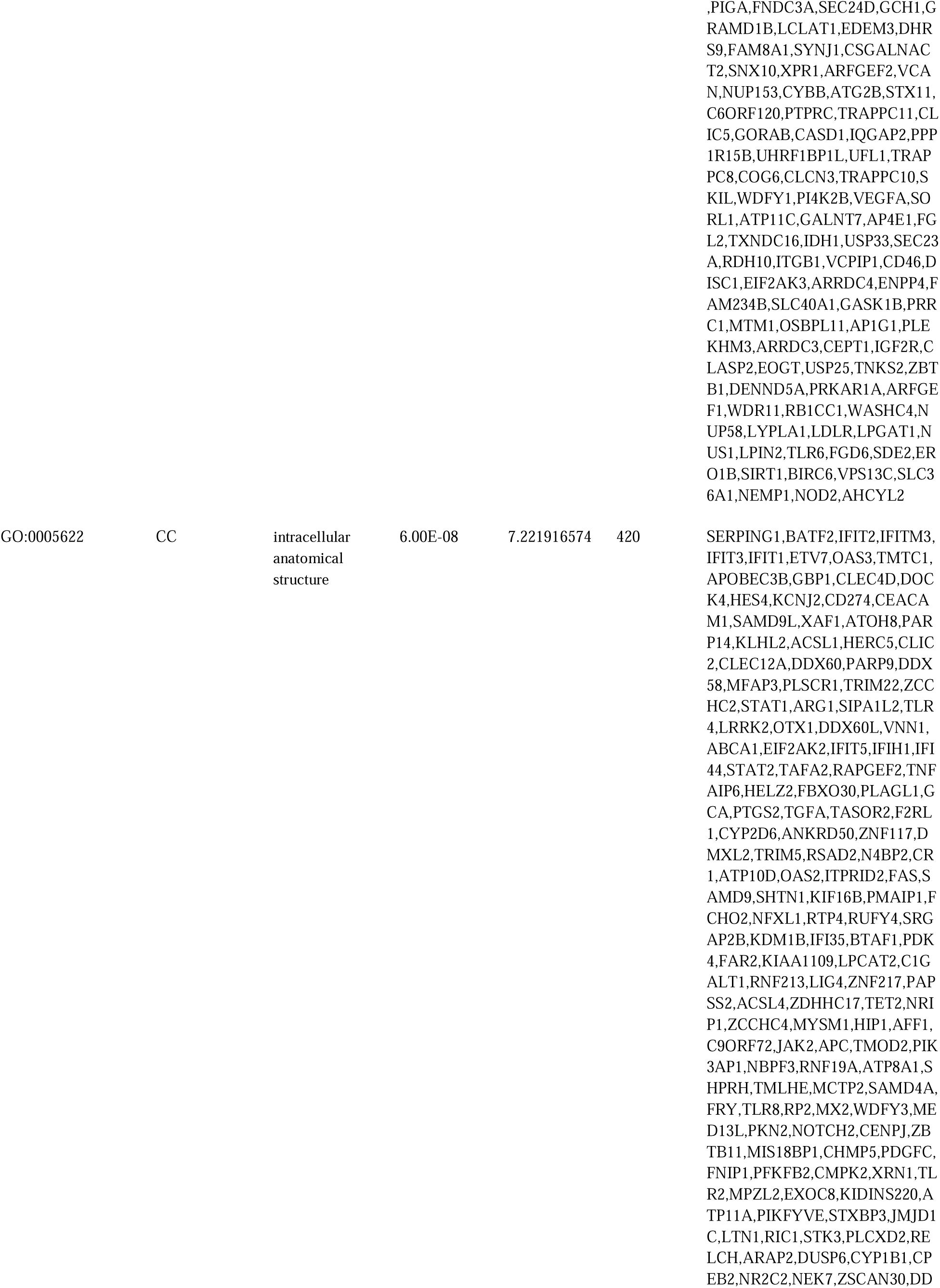
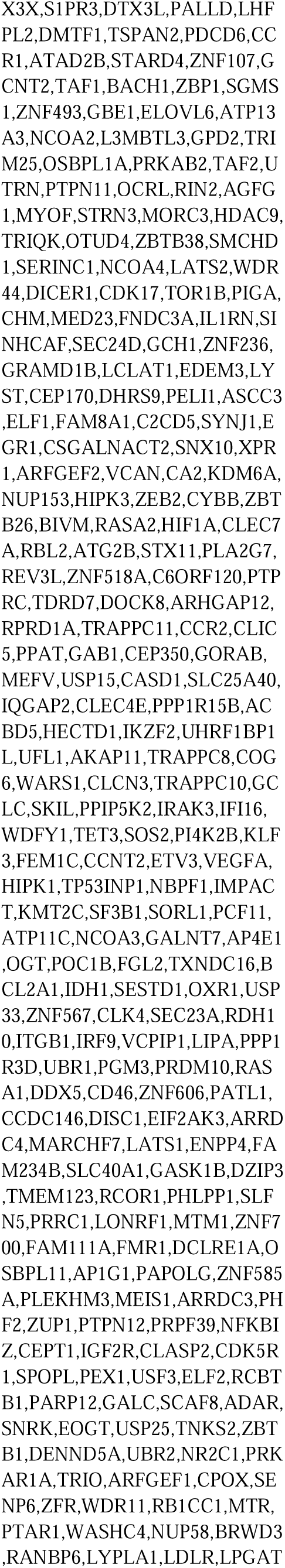
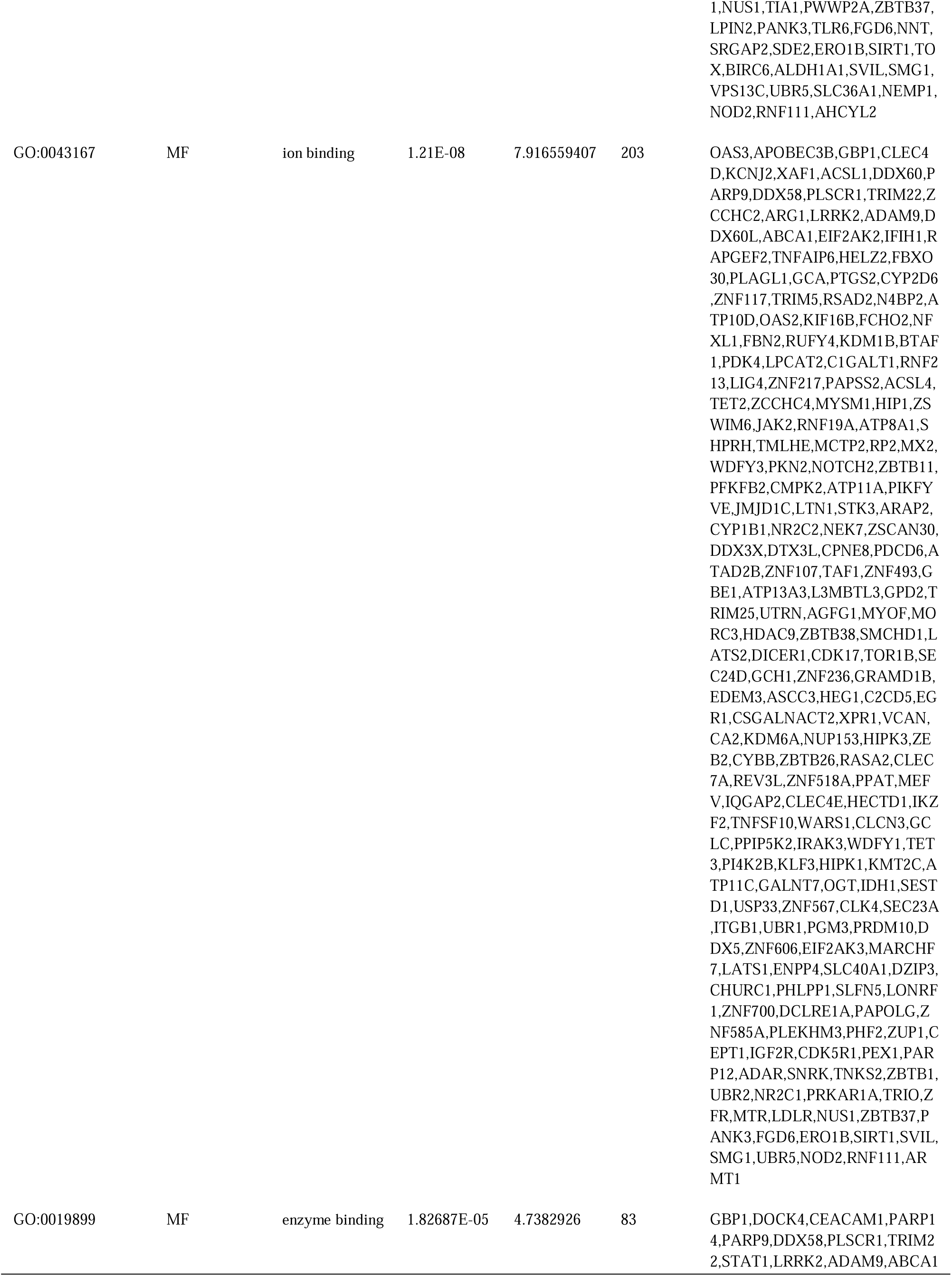
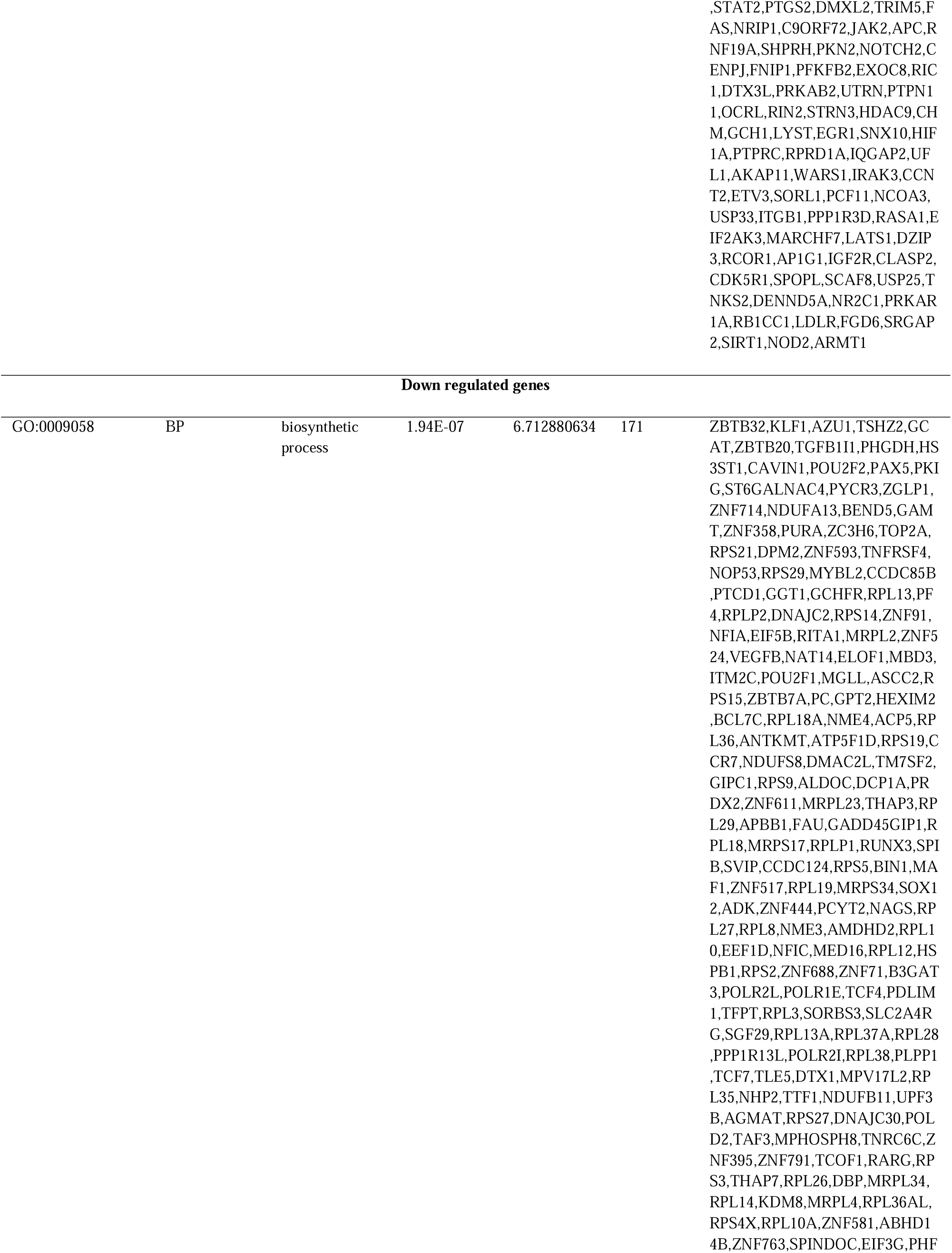
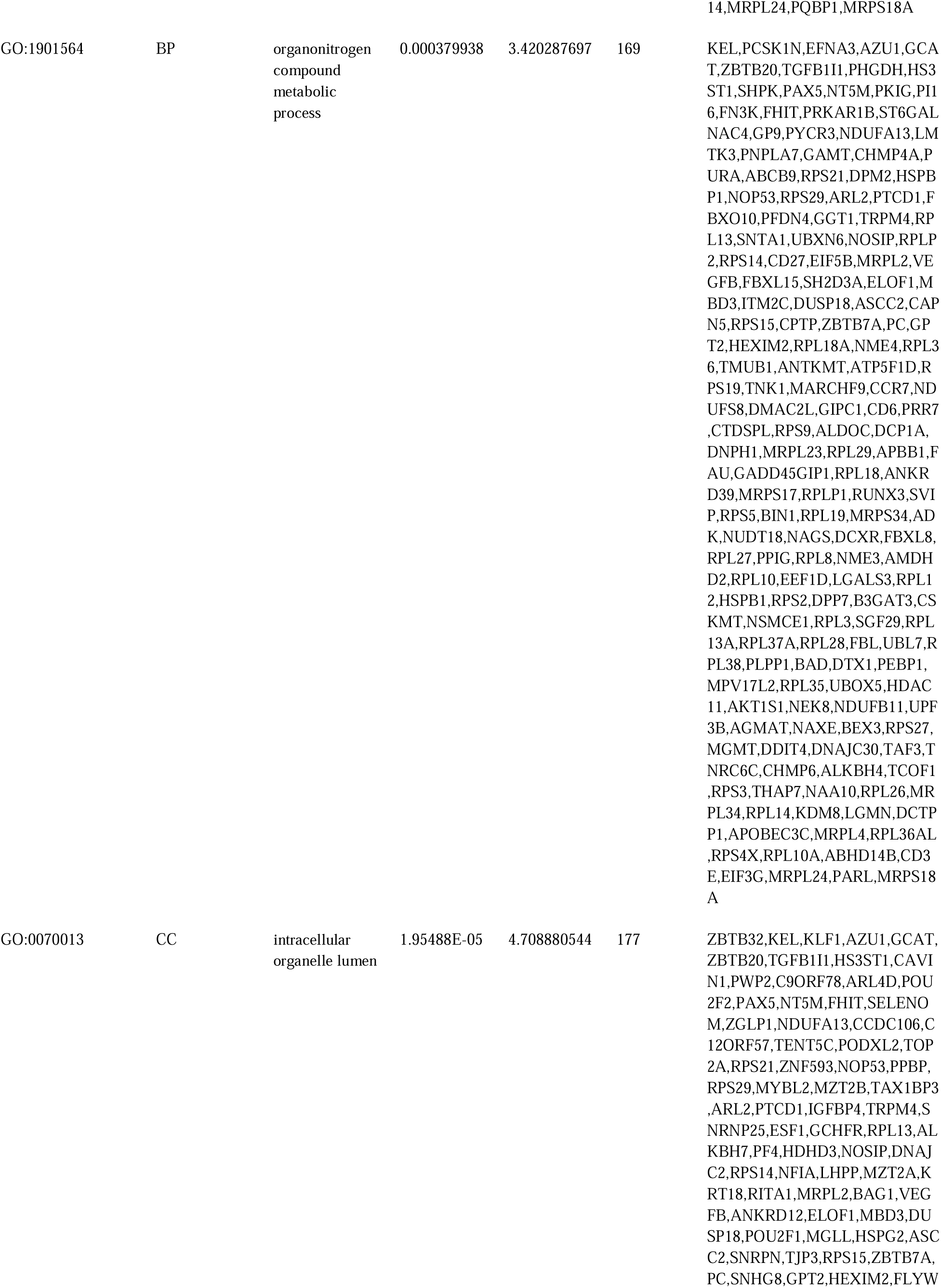
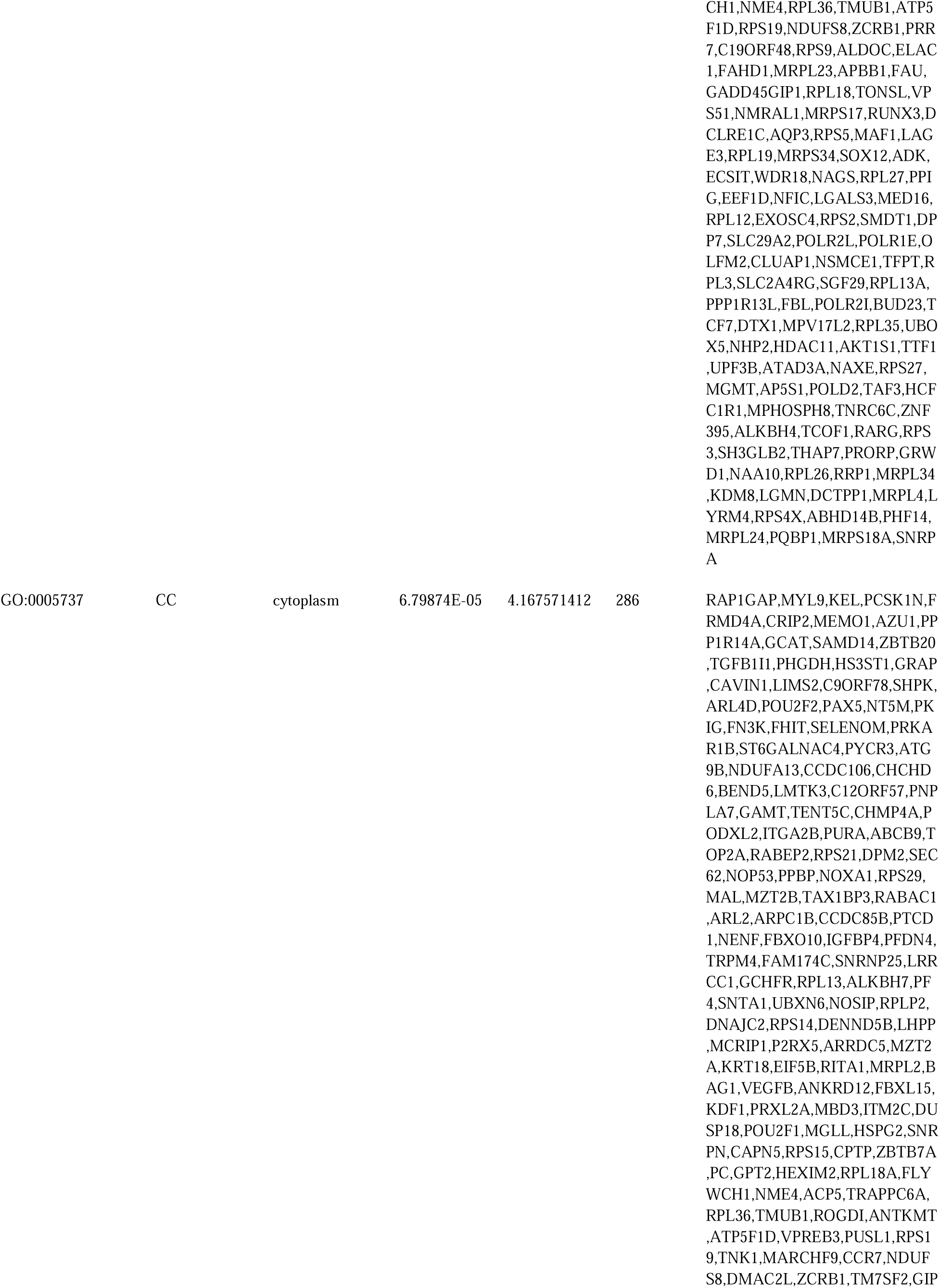
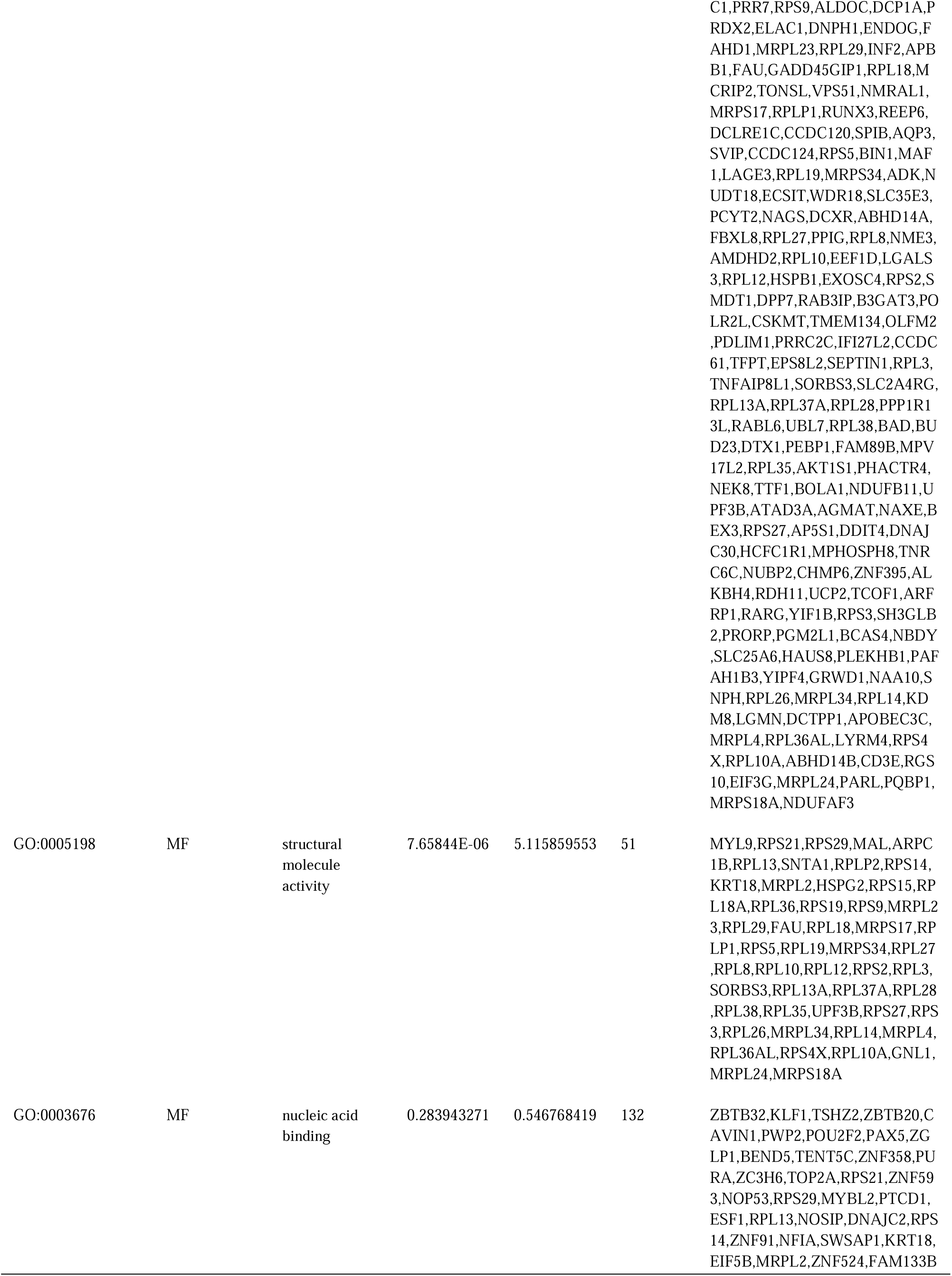
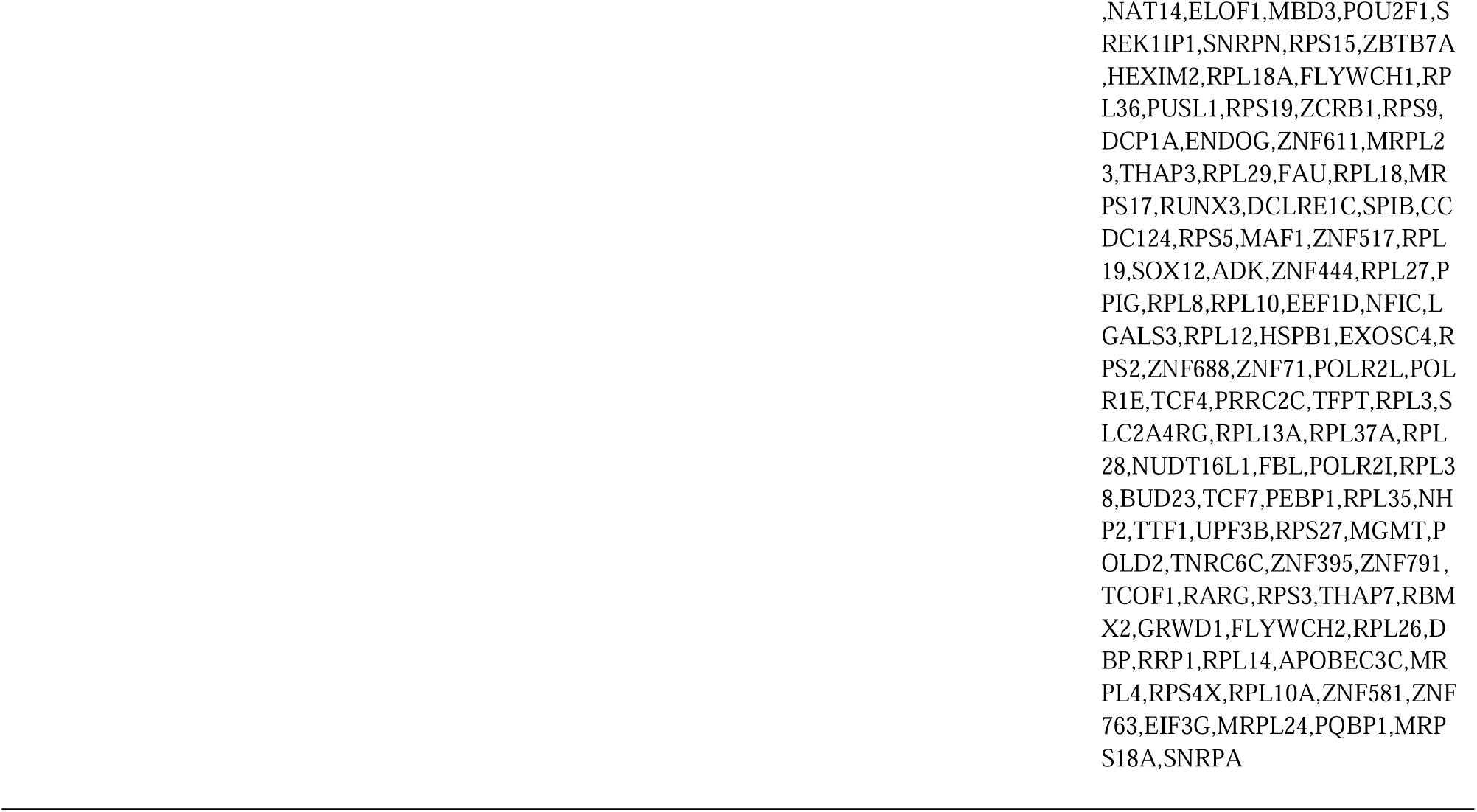
The enriched GO terms of the up and down regulated differentially expressed genes

**Table 3.**
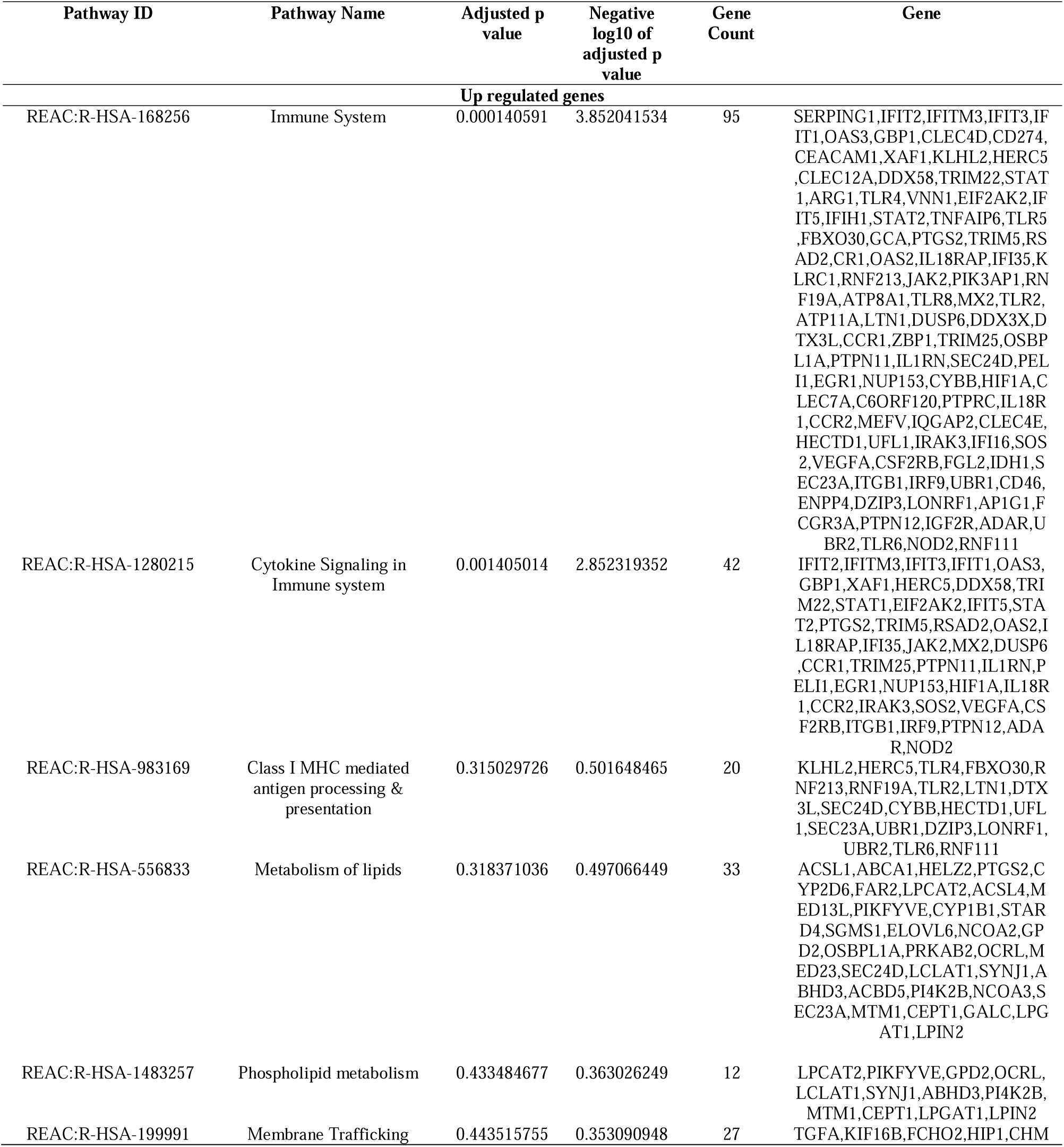
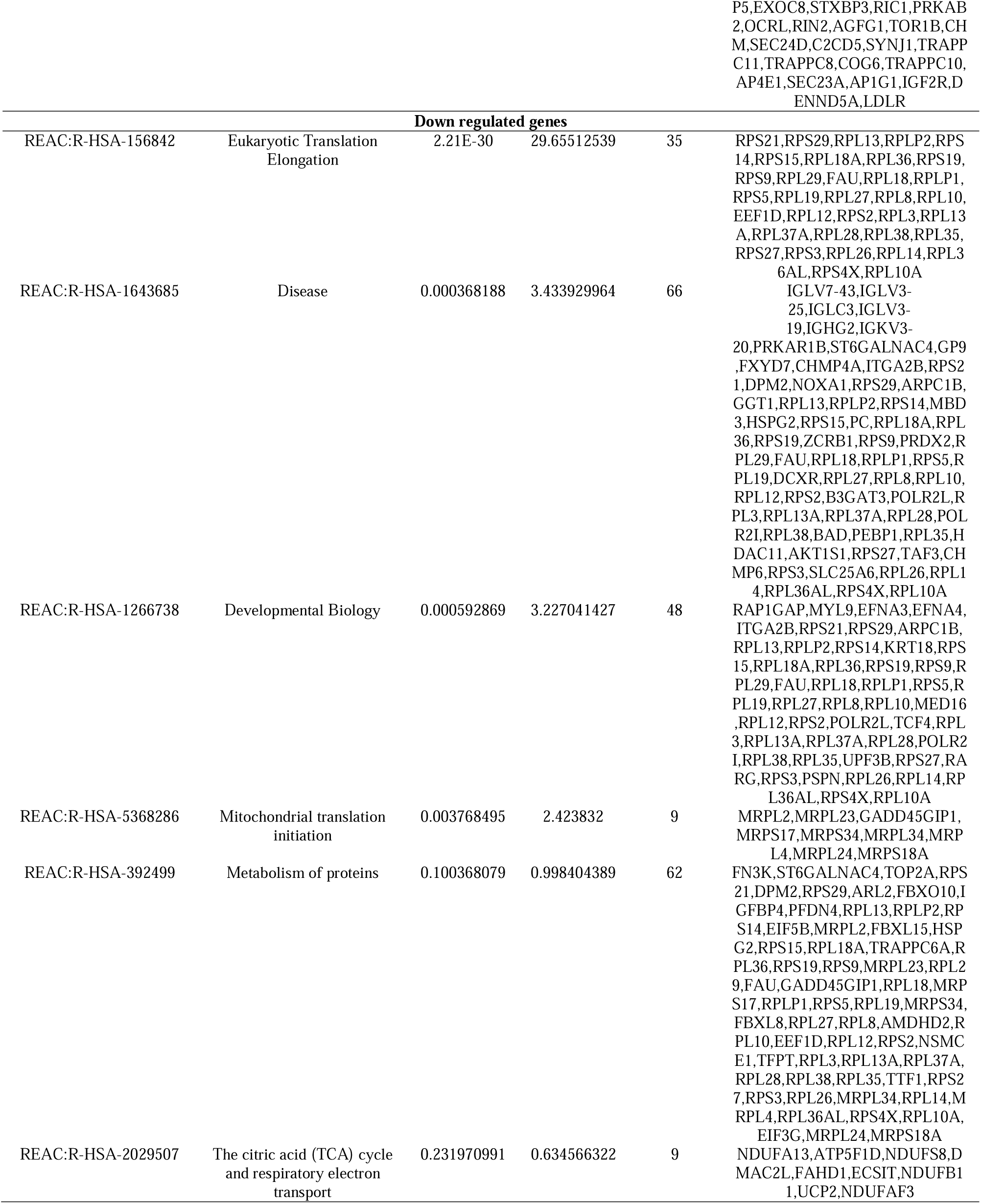
The enriched pathway terms of the up and down regulated differentially expressed genes

### Construction of the PPI network and module analysis

The PPI network of the DEGs was constructed using IMEx interactome. In the PPI network, there were 4863 nodes and 13037 interactions (Fig 3). The hub nodes were EGR1, SIRT1, STAT1, LRRK2, HIF1A, CSNK2B, RPS3, RPS2, RPS4X and HDAC11 (Table 4). The PEWCC1 plug-in was used to screen functional modules of the PPI network, which show a sub-network of highly interconnected proteins. The module 1 contained 26 nodes and 78 interactions (Fig. 4A). Module 1 was linked with immune system, cytokine signaling in immune system, cellular response to stimulus, biological regulation, intracellular anatomical structure and endomembrane system. The module 2 contained 29 nodes and 230 interactions (Fig. 4B). Module 2 was linked with eukaryotic translation elongation, disease, developmental biology, metabolism of proteins, biosynthetic process, organonitrogen compound metabolic process, intracellular organelle lumen, structural molecule activity and nucleic acid binding.

**Fig. 3.**
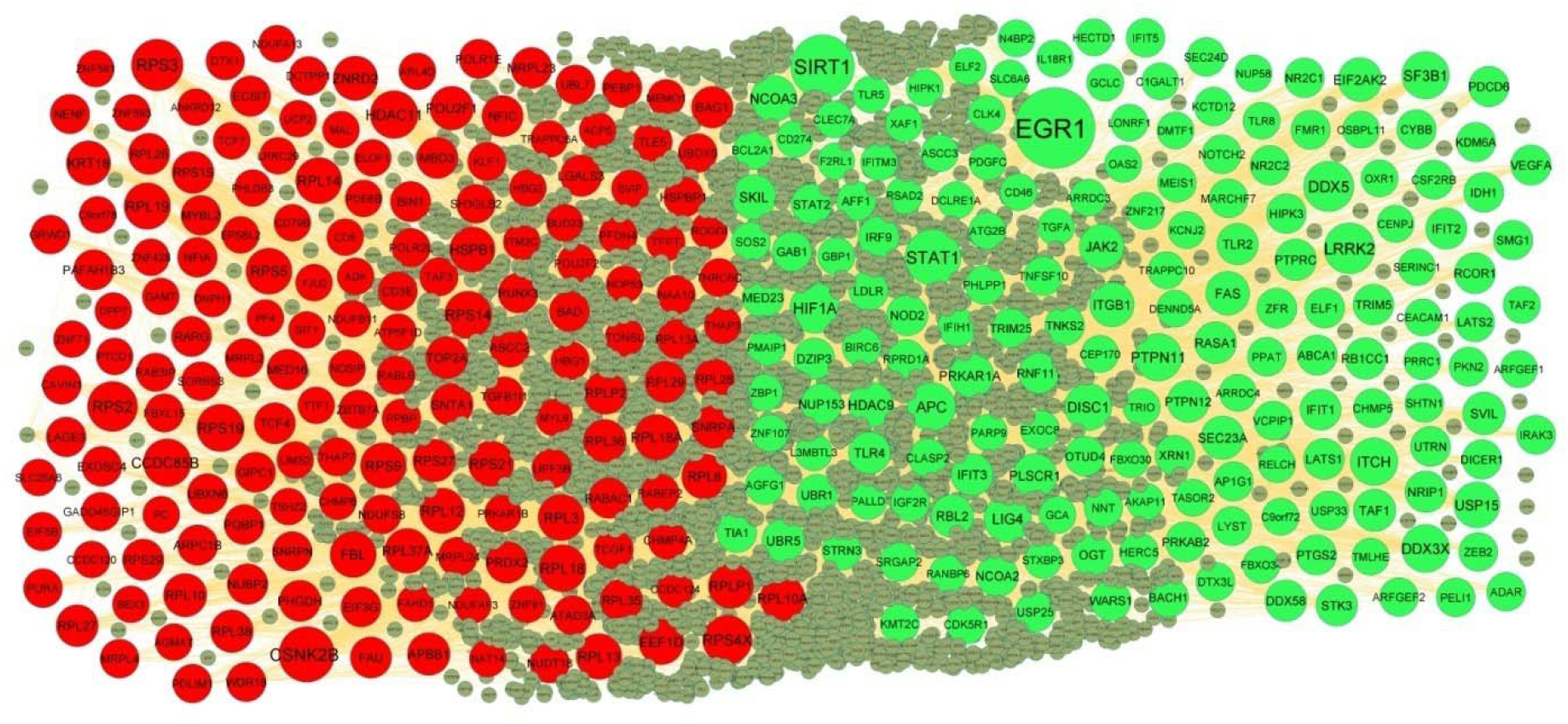
PPI network of DEGs. Up regulated genes are marked in green; down regulated genes are marked in red

**Fig. 4.**
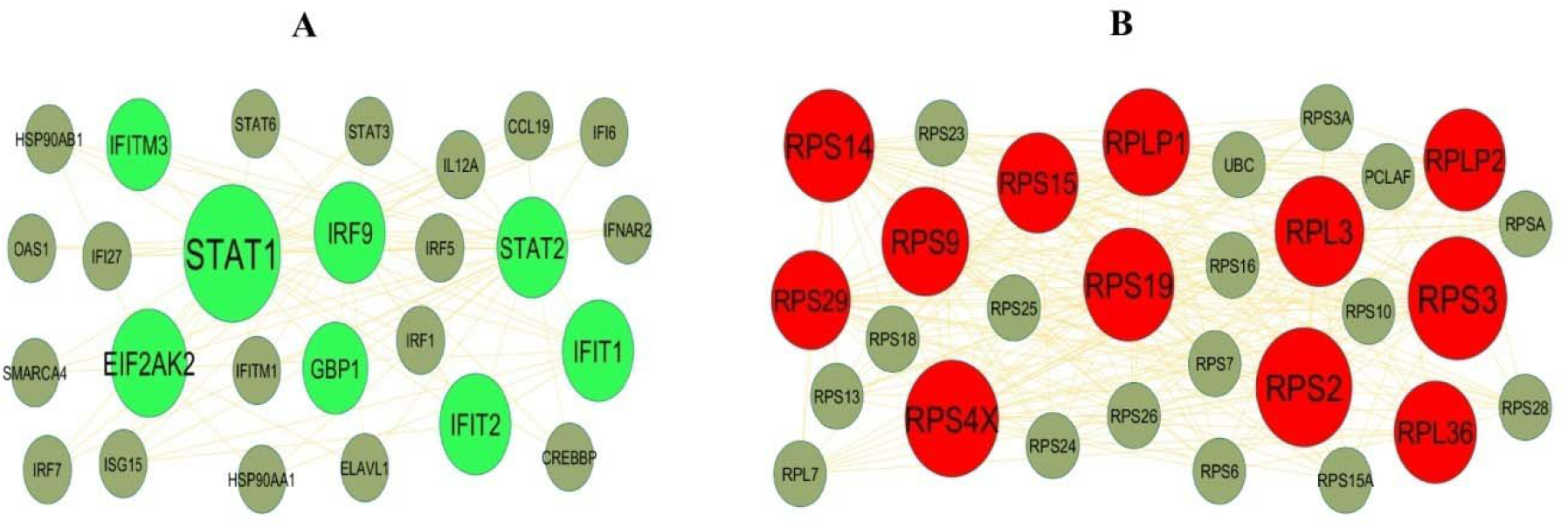
Modules selected from the DEG PPI between patients with FSGS and normal controls. (A) The most significant module was obtained from PPI network with 26 nodes and 78 edges for up regulated genes (B) The most significant module was obtained from PPI network with 29 nodes and 230 edges for down regulated genes. Up regulated genes are marked in green; down regulated genes are marked in red

**Table 4.** Topology table for up and down regulated genes

| Regulation | Node | Degree | Betweenness | Stress | Closeness |
| --- | --- | --- | --- | --- | --- |
| Up | EGR1 | 518 | 0.155691 | 51097908 | 0.373483 |
| Up | SIRT1 | 314 | 0.080189 | 32134000 | 0.36686 |
| Up | STAT1 | 223 | 0.049509 | 22461952 | 0.360228 |
| Up | LRRK2 | 164 | 0.030545 | 13490182 | 0.347112 |
| Up | HIF1A | 156 | 0.031431 | 15278362 | 0.358132 |
| Up | DDX5 | 156 | 0.021312 | 13124370 | 0.353754 |
| Up | APC | 134 | 0.03062 | 8157278 | 0.352549 |
| Up | DDX3X | 129 | 0.01346 | 7508184 | 0.353317 |
| Up | SF3B1 | 125 | 0.017765 | 7636820 | 0.356112 |
| Up | ITCH | 125 | 0.022984 | 9612406 | 0.341769 |
| Up | LIG4 | 123 | 0.028987 | 6646986 | 0.322992 |
| Up | PTPN11 | 122 | 0.025132 | 7810708 | 0.344261 |
| Up | DISC1 | 117 | 0.024269 | 5646774 | 0.319364 |
| Up | NCOA3 | 108 | 0.014326 | 8210920 | 0.34684 |
| Up | USP15 | 106 | 0.021378 | 8044676 | 0.339028 |
| Up | HDAC9 | 96 | 0.01859 | 4353434 | 0.345214 |
| Up | JAK2 | 92 | 0.01367 | 4674124 | 0.345337 |
| Up | SKIL | 90 | 0.019069 | 4779812 | 0.343216 |
| Up | ITGB1 | 88 | 0.020231 | 6343496 | 0.339146 |
| Up | PRKAR1A | 85 | 0.015974 | 5349082 | 0.343168 |
| Up | PLSCR1 | 84 | 0.021078 | 4243846 | 0.350339 |
| Up | EIF2AK2 | 81 | 0.011589 | 4309902 | 0.345779 |
| Up | FAS | 79 | 0.012899 | 4058128 | 0.354192 |
| Up | SEC23A | 76 | 0.013578 | 5658738 | 0.33792 |
| Up | UBR5 | 69 | 0.009193 | 3786410 | 0.341529 |
| Up | RBL2 | 69 | 0.011976 | 4158150 | 0.340548 |
| Up | SVIL | 67 | 0.011621 | 3322418 | 0.340167 |
| Up | TAF1 | 63 | 0.011788 | 3175596 | 0.341002 |
| Up | IFIT3 | 60 | 0.004025 | 1299464 | 0.320205 |
| Up | RASA1 | 59 | 0.009402 | 3115094 | 0.336774 |
| Up | TLR4 | 58 | 0.010616 | 3186120 | 0.335056 |
| Up | NCOA2 | 58 | 0.006941 | 2732602 | 0.341433 |
| Up | PTPN12 | 57 | 0.010002 | 2421710 | 0.338061 |
| Up | DDX58 | 56 | 0.006503 | 1834648 | 0.337498 |
| Up | RB1CC1 | 56 | 0.008398 | 2355384 | 0.340667 |
| Up | STK3 | 55 | 0.009833 | 1712866 | 0.340238 |
| Up | NRIP1 | 54 | 0.00827 | 3272078 | 0.342105 |
| Up | PTPRC | 53 | 0.008794 | 2245892 | 0.332854 |
| Up | TLR2 | 52 | 0.005513 | 2069040 | 0.319406 |
| Up | MED23 | 52 | 0.00655 | 2458958 | 0.334918 |
| Up | NOD2 | 50 | 0.007945 | 2448312 | 0.33501 |
| Up | OGT | 50 | 0.005936 | 2454148 | 0.335033 |
| Up | IFIT2 | 49 | 0.00407 | 1266226 | 0.315325 |
| Up | IFIT1 | 49 | 0.004249 | 1046646 | 0.323529 |
| Up | DZIP3 | 49 | 0.005721 | 1898762 | 0.331832 |
| Up | NUP153 | 48 | 0.007755 | 2222394 | 0.352089 |
| Up | RCOR1 | 47 | 0.005941 | 1928750 | 0.336331 |
| Up | IRF9 | 47 | 0.004901 | 1168868 | 0.323982 |
| Up | LATS1 | 45 | 0.006286 | 1648222 | 0.338933 |
| Up | STAT2 | 45 | 0.004441 | 1404884 | 0.344188 |
| Up | UBR1 | 43 | 0.006883 | 1966348 | 0.330254 |
| Up | TRIM5 | 43 | 0.00492 | 1419914 | 0.336354 |
| Up | RNF111 | 42 | 0.003349 | 1439100 | 0.334756 |
| Up | PDCD6 | 40 | 0.004532 | 1868828 | 0.334641 |
| Up | PRKAB2 | 39 | 0.009131 | 1791642 | 0.331492 |
| Up | VEGFA | 39 | 0.004753 | 1575996 | 0.337803 |
| Up | PTGS2 | 39 | 0.003263 | 1647510 | 0.334964 |
| Up | WARS1 | 38 | 0.007643 | 1386458 | 0.343531 |
| Up | ABCA1 | 37 | 0.009943 | 1770442 | 0.350794 |
| Up | STRN3 | 36 | 0.00712 | 1163202 | 0.340024 |
| Up | TRIM25 | 36 | 0.005366 | 1043530 | 0.340262 |
| Up | ZFR | 36 | 0.004606 | 1607878 | 0.335287 |
| Up | UTRN | 35 | 0.009868 | 1586706 | 0.340548 |
| Up | ELF1 | 35 | 0.005117 | 1379662 | 0.332717 |
| Up | IDH1 | 34 | 0.005267 | 1446080 | 0.330771 |
| Up | OTUD4 | 34 | 0.005666 | 1056004 | 0.334434 |
| Up | TIA1 | 33 | 0.003322 | 1355074 | 0.333883 |
| Up | USP25 | 33 | 0.005798 | 1286746 | 0.337522 |
| Up | GAB1 | 32 | 0.003575 | 736882 | 0.306809 |
| Up | HIPK3 | 32 | 0.004394 | 1067342 | 0.337007 |
| Up | TNKS2 | 32 | 0.007932 | 1127420 | 0.333448 |
| Up | XRN1 | 32 | 0.00463 | 1166800 | 0.332308 |
| Up | NR2C1 | 32 | 0.002237 | 881234 | 0.333906 |
| Up | NR2C2 | 31 | 0.002628 | 842738 | 0.315223 |
| Up | BACH1 | 30 | 0.003883 | 1156962 | 0.330479 |
| Up | DICER1 | 29 | 0.003258 | 1048798 | 0.328914 |
| Up | SMG1 | 29 | 0.003881 | 944060 | 0.333562 |
| Up | LATS2 | 29 | 0.002866 | 932064 | 0.334964 |
| Up | AFF1 | 28 | 0.004083 | 854336 | 0.344921 |
| Up | AP1G1 | 28 | 0.007669 | 984640 | 0.330748 |
| Up | LDLR | 27 | 0.004527 | 823592 | 0.345288 |
| Up | CYBB | 27 | 0.003654 | 730344 | 0.338933 |
| Up | KDM6A | 27 | 0.003959 | 815862 | 0.331267 |
| Up | BCL2A1 | 26 | 0.003065 | 536256 | 0.304046 |
| Up | CSF2RB | 26 | 0.002156 | 669768 | 0.337498 |
| Up | USP33 | 26 | 0.002994 | 730842 | 0.333402 |
| Up | CDK5R1 | 26 | 0.004459 | 939900 | 0.345386 |
| Up | FMR1 | 26 | 0.003111 | 963508 | 0.332263 |
| Up | EXOC8 | 26 | 0.006786 | 1054730 | 0.338226 |
| Up | AGFG1 | 25 | 0.002709 | 802666 | 0.332626 |
| Up | PHLPP1 | 25 | 0.00394 | 865598 | 0.331425 |
| Up | IGF2R | 25 | 0.00464 | 1190522 | 0.331628 |
| Up | PKN2 | 25 | 0.002459 | 733000 | 0.333768 |
| Up | TNFSF10 | 24 | 0.00253 | 533778 | 0.299292 |
| Up | VCPIP1 | 24 | 0.003847 | 626002 | 0.331041 |
| Up | KMT2C | 24 | 0.002698 | 821542 | 0.331832 |
| Up | BIRC6 | 24 | 0.003247 | 830228 | 0.330187 |
| Up | IRAK3 | 24 | 0.004322 | 1008336 | 0.329784 |
| Up | LYST | 24 | 0.00515 | 1052670 | 0.329493 |
| Up | NOTCH2 | 23 | 0.003516 | 733162 | 0.32858 |
| Up | CHMP5 | 23 | 0.00412 | 696206 | 0.325217 |
| Up | IFIH1 | 22 | 0.001947 | 527982 | 0.33386 |
| Up | CLASP2 | 22 | 0.00263 | 721240 | 0.332922 |
| Up | ZEB2 | 22 | 0.001047 | 587914 | 0.331289 |
| Up | HERC5 | 22 | 0.001864 | 592046 | 0.344237 |
| Up | DTX3L | 22 | 0.001464 | 651102 | 0.325544 |
| Up | MARCHF7 | 22 | 0.00212 | 515132 | 0.323465 |
| Up | MEIS1 | 21 | 0.003315 | 2580846 | 0.284877 |
| Up | PELI1 | 21 | 0.001105 | 479568 | 0.32927 |
| Up | SOS2 | 21 | 9.67E-04 | 345366 | 0.30029 |
| Up | KCTD12 | 21 | 0.001914 | 677242 | 0.328669 |
| Up | PPAT | 21 | 0.002572 | 660286 | 0.328758 |
| Up | SEC24D | 21 | 0.00203 | 616138 | 0.324826 |
| Up | ADAR | 21 | 0.001844 | 835666 | 0.348481 |
| Up | SRGAP2 | 20 | 0.002579 | 564926 | 0.333562 |
| Up | RPRD1A | 20 | 0.002845 | 718856 | 0.331357 |
| Up | SHTN1 | 20 | 0.002265 | 377498 | 0.340667 |
| Up | IFIT5 | 20 | 0.001655 | 410358 | 0.334227 |
| Up | ARFGEF2 | 20 | 0.003033 | 463622 | 0.334388 |
| Up | NNT | 19 | 0.00332 | 653754 | 0.330479 |
| Up | ARFGEF1 | 3 | 2.70E-05 | 11908 | 0.263209 |
| Up | IFITM3 | 3 | 0 | 0 | 0.267246 |
| Up | GBP1 | 3 | 0 | 0 | 0.267246 |
| Up | PMAIP1 | 2 | 1.31E-05 | 2106 | 0.240063 |
| Up | TAF2 | 2 | 3.21E-05 | 21792 | 0.272381 |
| Up | RSAD2 | 2 | 0 | 0 | 0.266076 |
| Up | STXBP3 | 2 | 6.84E-05 | 61250 | 0.282691 |
| Up | CEP170 | 2 | 0 | 0 | 0.255317 |
| Up | PALLD | 2 | 2.08E-04 | 29096 | 0.260209 |
| Up | F2RL1 | 1 | 0 | 0 | 0.250981 |
| Up | N4BP2 | 1 | 0 | 0 | 0.248848 |
| Up | CLEC7A | 1 | 0 | 0 | 0.242095 |
| Up | AKAP11 | 1 | 0 | 0 | 0.255505 |
| Up | ELF2 | 1 | 0 | 0 | 0.271939 |
| Up | SLC6A6 | 1 | 0 | 0 | 0.271939 |
| Up | C1GALT1 | 1 | 0 | 0 | 0.271939 |
| Up | PDGFC | 1 | 0 | 0 | 0.271939 |
| Up | DMTF1 | 1 | 0 | 0 | 0.271939 |
| Up | NUP58 | 1 | 0 | 0 | 0.260669 |
| Up | TLR8 | 1 | 0 | 0 | 0.264844 |
| Up | XAF1 | 1 | 0 | 0 | 0.264844 |
| Up | CD274 | 1 | 0 | 0 | 0.25611 |
| Up | CEACAM1 | 1 | 0 | 0 | 0.25611 |
| Up | ZBP1 | 1 | 0 | 0 | 0.24941 |
| Up | TGFA | 1 | 0 | 0 | 0.230985 |
| Up | KCNJ2 | 1 | 0 | 0 | 0.230985 |
| Up | TASOR2 | 1 | 0 | 0 | 0.241951 |
| Up | PARP9 | 1 | 0 | 0 | 0.245605 |
| Up | RANBP6 | 1 | 0 | 0 | 0.24738 |
| Up | ARRDC4 | 1 | 0 | 0 | 0.254728 |
| Up | GCA | 1 | 0 | 0 | 0.254728 |
| Up | ARRDC3 | 1 | 0 | 0 | 0.254728 |
| Up | CENPJ | 1 | 0 | 0 | 0.247846 |
| Up | ZNF107 | 1 | 0 | 0 | 0.252074 |
| Up | LONRF1 | 1 | 0 | 0 | 0.24442 |
| Up | TRIO | 1 | 0 | 0 | 0.242071 |
| Up | TLR5 | 1 | 0 | 0 | 0.250981 |
| Up | ZNF217 | 1 | 0 | 0 | 0.251695 |
| Up | CD46 | 1 | 0 | 0 | 0.253269 |
| Up | OAS2 | 1 | 0 | 0 | 0.250955 |
| Up | GCLC | 1 | 0 | 0 | 0.271939 |
| Up | TRAPPC10 | 1 | 0 | 0 | 0.188559 |
| Up | IL18R1 | 1 | 0 | 0 | 0.271939 |
| Up | DENND5A | 1 | 0 | 0 | 0.265814 |
| Up | DCLRE1A | 1 | 0 | 0 | 0.268411 |
| Up | HIPK1 | 1 | 0 | 0 | 0.255532 |
| Up | ASCC3 | 1 | 0 | 0 | 0.255532 |
| Up | OXR1 | 1 | 0 | 0 | 0.24415 |
| Up | FBXO30 | 1 | 0 | 0 | 0.260669 |
| Up | ATG2B | 1 | 0 | 0 | 0.260042 |
| Up | L3MBTL3 | 1 | 0 | 0 | 0.257685 |
| Up | C9orf72 | 1 | 0 | 0 | 0.254116 |
| Up | PRRC1 | 1 | 0 | 0 | 0.248556 |
| Up | OSBPL11 | 1 | 0 | 0 | 0.248556 |
| Up | CLK4 | 1 | 0 | 0 | 0.271939 |
| Up | SERINC1 | 1 | 0 | 0 | 0.249052 |
| Up | TMLHE | 1 | 0 | 0 | 0.25405 |
| Up | FBXO34 | 1 | 0 | 0 | 0.256638 |
| Up | RELCH | 1 | 0 | 0 | 0.252362 |
| Up | HECTD1 | 1 | 0 | 0 | 0.260669 |
| Down | CSNK2B | 222 | 0.056174 | 17631706 | 0.362025 |
| Down | RPS3 | 176 | 0.012295 | 9257168 | 0.355799 |
| Down | RPS2 | 156 | 0.009378 | 5766964 | 0.357054 |
| Down | RPS4X | 145 | 0.005665 | 5003802 | 0.351962 |
| Down | HDAC11 | 130 | 0.030131 | 5932312 | 0.351402 |
| Down | CCDC85B | 129 | 0.032481 | 5281170 | 0.319155 |
| Down | RPS19 | 126 | 0.008243 | 5318220 | 0.350768 |
| Down | RPS14 | 123 | 0.005871 | 5083400 | 0.348056 |
| Down | RPL18 | 122 | 0.003792 | 3647522 | 0.353189 |
| Down | RPL19 | 115 | 0.0038 | 2987980 | 0.351199 |
| Down | RPL3 | 112 | 0.006259 | 4717518 | 0.348655 |
| Down | RPL14 | 111 | 0.003611 | 2989780 | 0.351047 |
| Down | RPS5 | 110 | 0.004451 | 3267918 | 0.34863 |
| Down | FBL | 109 | 0.011851 | 5023504 | 0.34868 |
| Down | RPL13 | 106 | 0.006492 | 4256224 | 0.360015 |
| Down | RPS9 | 104 | 0.00379 | 3744910 | 0.350794 |
| Down | RPL18A | 103 | 0.004506 | 2954966 | 0.348156 |
| Down | RPL12 | 101 | 0.003356 | 2961874 | 0.349031 |
| Down | RPS21 | 100 | 0.006374 | 2945134 | 0.350515 |
| Down | HSPB1 | 100 | 0.01647 | 7946014 | 0.342033 |
| Down | RPL8 | 99 | 0.00578 | 3226670 | 0.346124 |
| Down | EEF1D | 99 | 0.014903 | 6249736 | 0.341697 |
| Down | RPL10A | 98 | 0.003901 | 2869994 | 0.354373 |
| Down | RPLP1 | 97 | 0.009168 | 4520058 | 0.348956 |
| Down | RPL37A | 96 | 0.003973 | 2574564 | 0.34519 |
| Down | ZNRD2 | 91 | 0.016749 | 5308632 | 0.341361 |
| Down | KRT18 | 89 | 0.019649 | 7044288 | 0.348505 |
| Down | POU2F1 | 82 | 0.013443 | 5840974 | 0.339834 |
| Down | RPL10 | 78 | 0.003095 | 2523704 | 0.34573 |
| Down | RPL36 | 72 | 0.002491 | 2063476 | 0.343507 |
| Down | RPLP2 | 72 | 0.00291 | 2184006 | 0.343507 |
| Down | RPL38 | 71 | 0.004592 | 2141760 | 0.34295 |
| Down | RPS27 | 71 | 0.006415 | 2530570 | 0.344066 |
| Down | SNTA1 | 69 | 0.018872 | 2462930 | 0.300346 |
| Down | APBB1 | 69 | 0.009676 | 2780082 | 0.343289 |
| Down | SNRPA | 68 | 0.008238 | 3614266 | 0.342853 |
| Down | RPL35 | 67 | 0.003092 | 2148100 | 0.341937 |
| Down | TOP2A | 64 | 0.005513 | 4072480 | 0.341409 |
| Down | RPS15 | 62 | 0.004076 | 1866096 | 0.336494 |
| Down | RPL27 | 61 | 0.002978 | 1678152 | 0.338485 |
| Down | PRDX2 | 58 | 0.008991 | 3958342 | 0.336657 |
| Down | RPL29 | 57 | 0.002603 | 1419924 | 0.339478 |
| Down | BAG1 | 57 | 0.007676 | 3434558 | 0.336238 |
| Down | LGALS3 | 56 | 0.00845 | 1865196 | 0.324544 |
| Down | RPS29 | 55 | 0.002489 | 832996 | 0.340024 |
| Down | PHGDH | 54 | 0.005938 | 2441924 | 0.343434 |
| Down | RPL26 | 53 | 0.002942 | 1758546 | 0.339952 |
| Down | MBD3 | 53 | 0.006462 | 1870582 | 0.339549 |
| Down | FAU | 53 | 0.001131 | 1036104 | 0.34225 |
| Down | NFIC | 52 | 0.010634 | 3092564 | 0.347783 |
| Down | TCF4 | 51 | 0.008284 | 1807164 | 0.349633 |
| Down | MRPL23 | 50 | 0.007556 | 1265756 | 0.336447 |
| Down | NUBP2 | 46 | 0.006086 | 1674116 | 0.330636 |
| Down | MED16 | 46 | 0.006184 | 1909986 | 0.333906 |
| Down | ECSIT | 44 | 0.006864 | 1020928 | 0.308934 |
| Down | RABAC1 | 43 | 0.01234 | 2137826 | 0.335982 |
| Down | BIN1 | 43 | 0.0067 | 2582058 | 0.308698 |
| Down | PAFAH1B3 | 43 | 0.006128 | 2093888 | 0.338391 |
| Down | ARPC1B | 42 | 0.005643 | 1431032 | 0.337288 |
| Down | UBXN6 | 42 | 0.007469 | 1807124 | 0.333425 |
| Down | GIPC1 | 41 | 0.009058 | 1572624 | 0.330951 |
| Down | RARG | 41 | 0.008772 | 1169788 | 0.334664 |
| Down | EIF3G | 40 | 0.004085 | 1322146 | 0.333516 |
| Down | RPL28 | 40 | 0.001195 | 931888 | 0.338344 |
| Down | BAD | 39 | 0.004502 | 1531146 | 0.32188 |
| Down | EXOSC4 | 39 | 0.009295 | 1821656 | 0.326134 |
| Down | ASCC2 | 38 | 0.007002 | 1168692 | 0.337662 |
| Down | HSPBP1 | 38 | 0.003444 | 1325910 | 0.3329 |
| Down | PQBP1 | 38 | 0.006776 | 1400710 | 0.33785 |
| Down | MYBL2 | 36 | 0.004044 | 3335808 | 0.306075 |
| Down | NFIA | 35 | 0.002543 | 824542 | 0.294756 |
| Down | LAGE3 | 35 | 0.006065 | 1324068 | 0.325566 |
| Down | NUDT18 | 35 | 0.007306 | 684202 | 0.279892 |
| Down | RUNX3 | 35 | 0.00354 | 1330910 | 0.334319 |
| Down | UPF3B | 33 | 0.003086 | 1140422 | 0.329694 |
| Down | PEBP1 | 33 | 0.00281 | 1236570 | 0.330726 |
| Down | TLE5 | 32 | 0.00485 | 1046816 | 0.334388 |
| Down | SNRPN | 32 | 0.002554 | 849514 | 0.336984 |
| Down | TCOF1 | 32 | 0.00472 | 1623208 | 0.332399 |
| Down | NENF | 31 | 0.003779 | 689502 | 0.33538 |
| Down | RPL13A | 31 | 0.003028 | 1650092 | 0.325304 |
| Down | POLR1E | 30 | 0.004456 | 818072 | 0.333837 |
| Down | ATAD3A | 30 | 0.001638 | 1033330 | 0.324978 |
| Down | CHMP4A | 30 | 0.004822 | 904830 | 0.333997 |
| Down | WDR18 | 30 | 0.002192 | 817698 | 0.332195 |
| Down | CD3E | 28 | 0.006746 | 1235934 | 0.333676 |
| Down | UBL7 | 28 | 0.003366 | 798026 | 0.332149 |
| Down | NDUFS8 | 27 | 0.005599 | 1024420 | 0.32188 |
| Down | TGFB1I1 | 27 | 0.003891 | 793770 | 0.335079 |
| Down | SORBS3 | 26 | 0.003948 | 3906396 | 0.273361 |
| Down | TONSL | 26 | 0.003067 | 864156 | 0.33138 |
| Down | MRPL24 | 26 | 0.004476 | 853098 | 0.326835 |
| Down | PURA | 25 | 0.003175 | 1040344 | 0.348481 |
| Down | CAVIN1 | 25 | 0.002887 | 661460 | 0.335264 |
| Down | GRWD1 | 25 | 0.001986 | 1011652 | 0.324891 |
| Down | UBOX5 | 25 | 0.00154 | 616704 | 0.331109 |
| Down | ZBTB7A | 24 | 0.001803 | 493312 | 0.323637 |
| Down | POU2F2 | 24 | 0.001553 | 739578 | 0.332786 |
| Down | PPBP | 24 | 0.0049 | 805220 | 0.293386 |
| Down | MRPL4 | 24 | 0.003024 | 868942 | 0.325609 |
| Down | SH3GLB2 | 24 | 0.004486 | 768154 | 0.324588 |
| Down | MRPL2 | 24 | 0.002496 | 599756 | 0.325719 |
| Down | GADD45GIP1 | 23 | 0.002025 | 652996 | 0.338179 |
| Down | TFPT | 23 | 0.004884 | 2734812 | 0.296445 |
| Down | PDLIM1 | 22 | 0.00278 | 722376 | 0.324566 |
| Down | DCTPP1 | 22 | 0.002215 | 760632 | 0.326791 |
| Down | EIF5B | 22 | 0.001181 | 612952 | 0.331719 |
| Down | DTX1 | 21 | 0.001238 | 435252 | 0.334802 |
| Down | POLR2L | 21 | 0.00407 | 851400 | 0.327121 |
| Down | NAA10 | 21 | 0.002348 | 526556 | 0.339312 |
| Down | TAF3 | 20 | 0.001056 | 228098 | 0.286708 |
| Down | KLF1 | 20 | 0.001571 | 516082 | 0.334802 |
| Down | THAP7 | 20 | 0.002718 | 952806 | 0.294025 |
| Down | RAB3IP | 20 | 0.004665 | 3742314 | 0.232365 |
| Down | FBXL15 | 19 | 0.002406 | 562466 | 0.330187 |
| Down | TNRC6C | 19 | 0.002762 | 4923668 | 0.276407 |
| Down | RABEP2 | 19 | 0.001975 | 540152 | 0.32838 |
| Down | RABL6 | 19 | 0.003071 | 2190512 | 0.281398 |
| Down | HBG1 | 19 | 0.001005 | 288998 | 0.307838 |
| Down | PFDN4 | 18 | 0.003287 | 554480 | 0.328114 |
| Down | TCF7 | 3 | 2.27E-04 | 42330 | 0.276958 |
| Down | NDUFA13 | 3 | 1.25E-04 | 23576 | 0.258686 |
| Down | PF4 | 2 | 1.02E-05 | 16122 | 0.257617 |
| Down | TRAPPC6A | 2 | 3.62E-04 | 307310 | 0.255036 |
| Down | ARL4D | 2 | 6.90E-06 | 2180 | 0.25389 |
| Down | MEMO1 | 2 | 8.81E-05 | 20782 | 0.264642 |
| Down | MAL | 1 | 0 | 0 | 0.250981 |
| Down | NDUFAF3 | 1 | 0 | 0 | 0.243514 |
| Down | SLC25A6 | 1 | 0 | 0 | 0.259459 |
| Down | EPS8L2 | 1 | 0 | 0 | 0.250206 |
| Down | CD79B | 1 | 0 | 0 | 0.249743 |
| Down | PRKAR1B | 1 | 0 | 0 | 0.255505 |
| Down | YJU2 | 1 | 0 | 0 | 0.271939 |
| Down | ELOF1 | 1 | 0 | 0 | 0.271939 |
| Down | LRRC29 | 1 | 0 | 0 | 0.271939 |
| Down | PDE6B | 1 | 0 | 0 | 0.271939 |
| Down | ANKRD12 | 1 | 0 | 0 | 0.254688 |
| Down | ACP5 | 1 | 0 | 0 | 0.230361 |
| Down | GAMT | 1 | 0 | 0 | 0.265814 |
| Down | ZNF71 | 1 | 0 | 0 | 0.265814 |
| Down | HBG2 | 1 | 0 | 0 | 0.250838 |
| Down | DNPH1 | 1 | 0 | 0 | 0.248277 |
| Down | CHMP6 | 1 | 0 | 0 | 0.250386 |
| Down | FAHD1 | 1 | 0 | 0 | 0.245941 |
| Down | ROGDI | 1 | 0 | 0 | 0.242071 |
| Down | C9orf78 | 1 | 0 | 0 | 0.248492 |
| Down | ZNF593 | 1 | 0 | 0 | 0.248277 |
| Down | BUD23 | 1 | 0 | 0 | 0.248938 |
| Down | PTCD1 | 1 | 0 | 0 | 0.248403 |
| Down | CD5 | 1 | 0 | 0 | 0.251943 |
| Down | PC | 1 | 0 | 0 | 0.244457 |
| Down | MYL9 | 1 | 0 | 0 | 0.257685 |
| Down | ATP5F1D | 1 | 0 | 0 | 0.245036 |
| Down | ADK | 1 | 0 | 0 | 0.271939 |
| Down | UCP2 | 1 | 0 | 0 | 0.268411 |
| Down | BEX3 | 1 | 0 | 0 | 0.241951 |
| Down | ZNF91 | 1 | 0 | 0 | 0.245419 |
| Down | TTF1 | 1 | 0 | 0 | 0.251097 |
| Down | PHLDB3 | 1 | 0 | 0 | 0.268411 |
| Down | LIMS2 | 1 | 0 | 0 | 0.252074 |
| Down | SVIP | 1 | 0 | 0 | 0.248721 |
| Down | THAP3 | 1 | 0 | 0 | 0.250968 |
| Down | NAT14 | 1 | 0 | 0 | 0.2515 |
| Down | ZNF428 | 1 | 0 | 0 | 0.24415 |
| Down | CCDC124 | 1 | 0 | 0 | 0.25695 |
| Down | CCDC120 | 1 | 0 | 0 | 0.241951 |
| Down | AGMAT | 1 | 0 | 0 | 0.25533 |
| Down | ITM2C | 1 | 0 | 0 | 0.271939 |
| Down | TSHZ2 | 1 | 0 | 0 | 0.255572 |
| Down | NDUFB11 | 1 | 0 | 0 | 0.226846 |
| Down | NOP53 | 1 | 0 | 0 | 0.25695 |
| Down | ZNF581 | 1 | 0 | 0 | 0.259459 |
| Down | DPP7 | 1 | 0 | 0 | 0.249436 |
| Down | NOSIP | 1 | 0 | 0 | 0.271939 |
| Down | SIT1 | 1 | 0 | 0 | 0.25611 |

### Construction of the miRNA-hub gene regulatory network

The miRNA-hub gene regulatory network included 2694 (miRNA: 2348; gene: 346) nodes and 22561 interactions (Fig 5). We discovered that DDX3X was targeted by 187 miRNAs (ex; hsa-mir-149-3p); DDX5 was targeted by 145 miRNAs (ex; hsa-mir-500b-5p); NCOA3 was targeted by 143 miRNAs (ex; hsa-mir-548y); EGR1 was targeted by 129 miRNAs (ex; hsa-mir-518b); HIF1A was targeted by 101 miRNAs (ex; hsa-mir-16-5p); RPS3 was targeted by 50 miRNAs (ex; hsa-mir-5699-3p); RPS4X was targeted by 40 miRNAs (ex; hsa-mir-532-3p); RPS14 was targeted by 40 miRNAs (ex; hsa-mir-503-5p); FBL was targeted by 39 miRNAs (ex; hsa-mir-31-5p); RPL3 was targeted by 32 miRNAs (ex; hsa-mir-365b-3p) and are listed in Table 5.

**Fig. 5.**
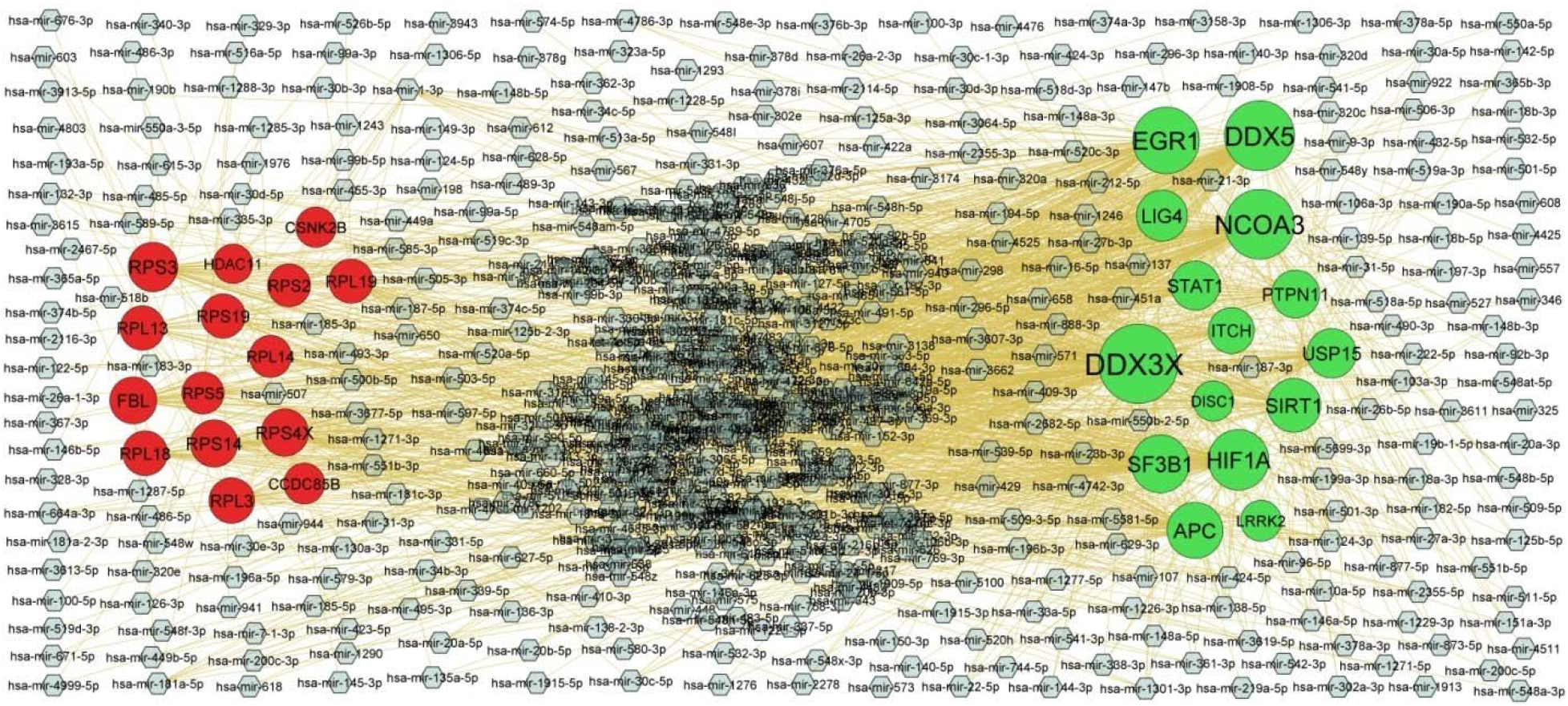
Target gene - miRNA regulatory network between target genes. The blue color diamond nodes represent the key miRNAs; up regulated genes are marked in green; down regulated genes are marked in red.

**Table 5.** MiRNA - hub gene and TF – hub gene topology table

| Regulation | Target Genes | Degree | MicroRNA | Regulation | Target Genes | Degree | TF |
| --- | --- | --- | --- | --- | --- | --- | --- |
| Up | DDX3X | 187 | hsa-mir-149-3p | Up | EGR1 | 53 | LMO2 |
| Up | DDX5 | 145 | hsa-mir-500b-5p | Up | DDX5 | 51 | MYCN |
| Up | NCOA3 | 143 | hsa-mir-548y | Up | NCOA3 | 48 | SIN3B |
| Up | EGR1 | 129 | hsa-mir-518b | Up | STAT1 | 42 | IRF8 |
| Up | HIF1A | 101 | hsa-mir-16-5p | Up | HIF1A | 40 | GATA1 |
| Up | SF3B1 | 89 | hsa-mir-181c-3p | Up | DISC1 | 37 | RUNX1 |
| Up | APC | 80 | hsa-mir-148a-3p | Up | USP15 | 34 | AP1S2 |
| Up | SIRT1 | 70 | hsa-mir-520c-3p | Up | APC | 34 | HOXB4 |
| Up | LIG4 | 57 | hsa-mir-501-3p | Up | SIRT1 | 33 | TBX3 |
| Up | USP15 | 50 | hsa-mir-20a-5p | Up | DDX3X | 32 | PDX1 |
| Up | STAT1 | 48 | hsa-mir-449a | Up | LIG4 | 31 | KDM6A |
| Up | PTPN11 | 43 | hsa-mir-30d-3p | Up | PTPN11 | 29 | TCF3 |
| Up | ITCH | 34 | hsa-mir-27a-3p | Up | SF3B1 | 29 | NANOG |
| Up | LRRK2 | 11 | hsa-mir-4476 | Up | ITCH | 17 | BCL3 |
| Up | DISC1 | 8 | hsa-mir-30d-5p | Up | LRRK2 | 16 | CNOT3 |
| Down | RPS3 | 50 | hsa-mir-5699-3p | Down | RPS14 | 45 | RCOR3 |
| Down | RPS4X | 40 | hsa-mir-532-3p | Down | RPL3 | 44 | DMRT1 |
| Down | RPS14 | 40 | hsa-mir-503-5p | Down | RPS19 | 42 | SREBF1 |
| Down | FBL | 39 | hsa-mir-31-5p | Down | RPL18 | 36 | FOXP3 |
| Down | RPL3 | 32 | hsa-mir-365b-3p | Down | RPS5 | 35 | POU5F1 |
| Down | RPL18 | 29 | hsa-mir-550a-3-5p | Down | CSNK2B | 35 | SOX11 |
| Down | RPL13 | 27 | hsa-mir-495-3p | Down | FBL | 33 | GATA2 |
| Down | RPL19 | 26 | hsa-mir-23b-3p | Down | RPL14 | 29 | FOXO3 |
| Down | RPS19 | 26 | hsa-mir-320e | Down | RPS3 | 27 | AR |
| Down | RPS2 | 20 | hsa-mir-378d | Down | RPL13 | 24 | ESR2 |
| Down | RPL14 | 16 | hsa-mir-424-5p | Down | RPL19 | 24 | RARG |
| Down | RPS5 | 16 | hsa-mir-103a-3p | Down | HDAC11 | 24 | MITF |
| Down | CCDC85B | 11 | hsa-mir-603 | Down | RPS2 | 22 | RELA |
| Down | CSNK2B | 9 | hsa-mir-124-3p | Down | CCDC85B | 19 | PPARD |
| Down | HDAC11 | 4 | hsa-mir-1306-5p | Down | RPS4X | 15 | XRN2 |

### Construction of the TF-hub gene regulatory network

The TF-hub gene regulatory network included 535 (TF: 181; gene: 354) nodes and 9293 interactions (Fig 6). We discovered that that EGR1 was targeted by 53 TFs (ex; LMO2); DDX5 was targeted by 51 TFs (ex; MYCN); NCOA3 was targeted by 48 TFs (ex; SIN3B); STAT1 was targeted by 42 TFs (ex; IRF8); HIF1A was targeted by 40 TFs (ex; GATA1); RPS14 was targeted by 45 TFs (ex; RCOR3); RPL3 was targeted by 44 TFs (ex; DMRT1); RPS19 was targeted by 42 TFs (ex; SREBF1); RPL18 was targeted by 36 TFs (ex; FOXP3); RPS5 was targeted by 35 TFs (ex; POU5F1) and are listed in Table 5.

**Fig. 6.**
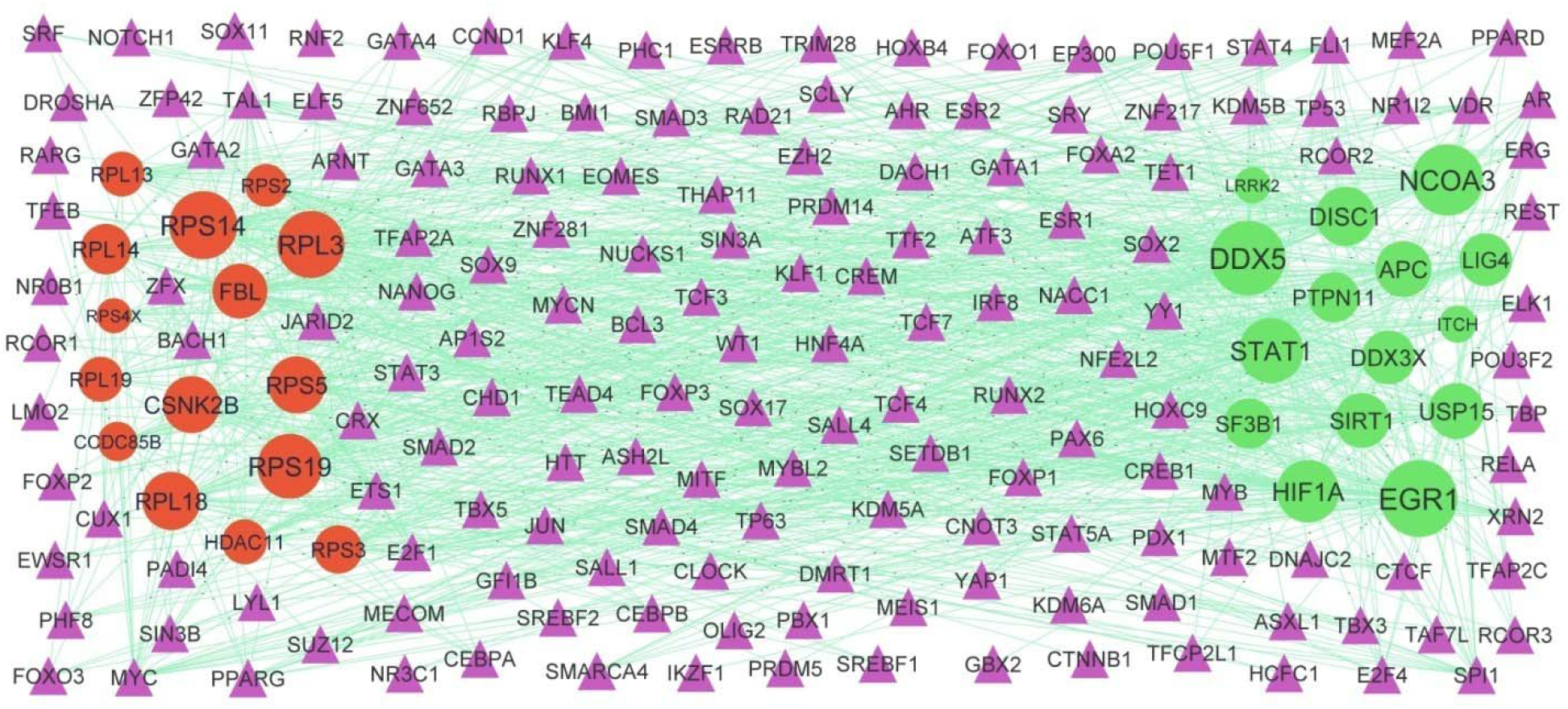
Target gene - TF regulatory network between target genes. The violet color triangle nodes represent the key TFs; up regulated genes are marked in green; down regulated genes are marked in red.

### Receiver operating characteristic curve (ROC) analysis

ROC curve analyses were performed to verify the hub genes, and area under the curve (AUC) values was calculated. Fig 7 shows the AUC for EGR1 was 0.958, AUC for SIRT1 was 0.917, AUC for STAT1 was 0.932, AUC for LRRK2 was 0.922, AUC for HIF1A was 0.912, AUC for CSNK2B was 0.910, AUC for RPS3 was 0.935, AUC for RPS2 was 0.953, AUC for RPS4X was 0.940 and AUC for HDAC11 was 0.907, indicating that the hub genes had high prognostic ability.

**Fig. 7.**
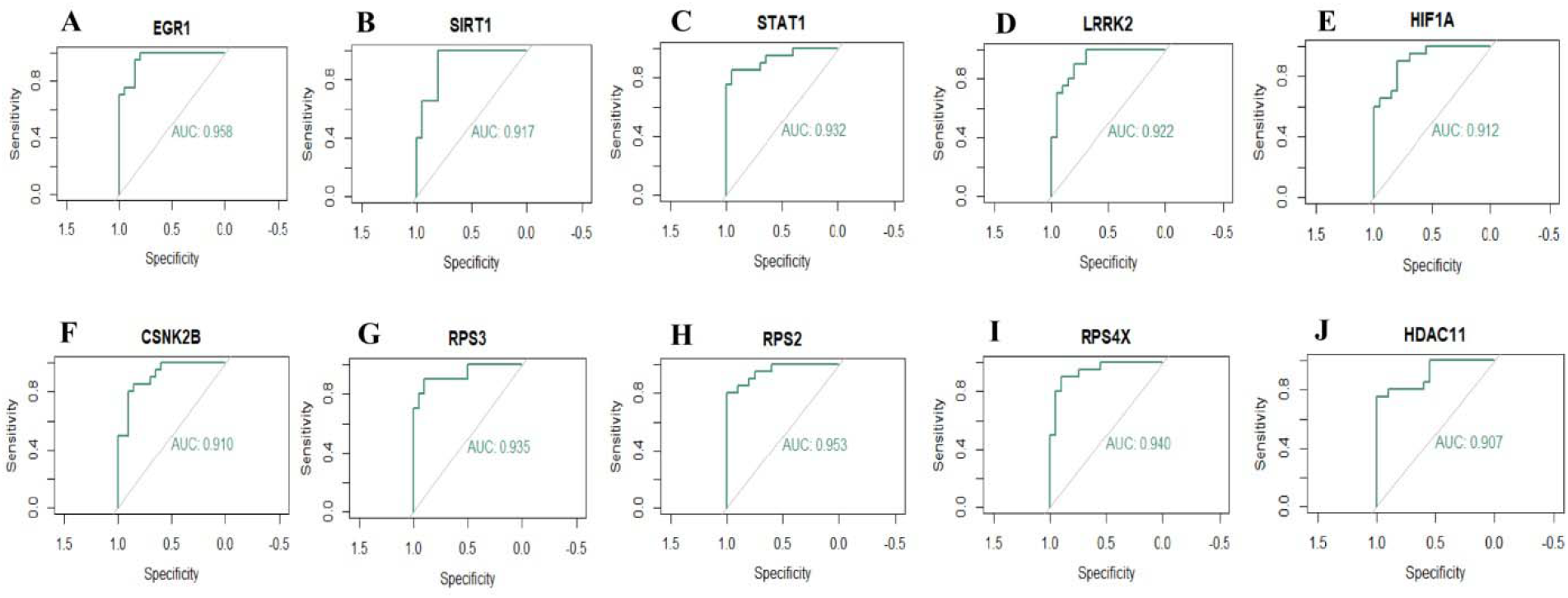
ROC curve analyses of hub genes. A) EGR1 B) SIRT1 C) STAT1 D) LRRK2 E) HIF1A F) CSNK2B G) RPS3 H) RPS2 I) RPS4X J) HDAC11

## Discussion

CAD is a highly frequent disease around world, which remains a dominant cause of cardiovascular death. Therefore, timely and appropriate diagnosis of CAD is essential to enhance the prognosis. In this investigation, we collected NGS dataset from the GEO database, and a total of 958 DEGs, including 479 up regulated genes and 479 down regulated genes, were found. SERPING1 [47] might play an important role in occurrence of development CAD. IFITM3 [48] were a novel key gene in obesity. IFITM3 [49] gene might be related to the pathophysiology of diabetes mellitus. IFITM3 [49] expression is related to the patients with hypercholesterolemia and its progression. Previous studies have shown that IFIT3 [50] might influence the prognosis in myocardial infarction. IFIT1 [51] might be involved in the pathogenesis of atherosclerosis. HBG1 [52] is important in the development of atrial fibrillation. RAP1GAP [53] was associated with cardiac hypertrophy. The results of this investigation suggest that DEGs might play a key role in the pathogenesis of CAD and CAD related complications.

The key DEGs and their relevant GO terms and signaling pathways were screened out based on a bioinformatic analysis. Signaling pathways include immune system [54], metabolism of lipids [55], phospholipid metabolism [56] and disease [57] plays an important role in CAD. DOCK4 [58], CEACAM1 [59], STAT1 [60], ARG1 [61], TLR4 [62], ADAM9 [63], VNN1 [64], ABCA1 [65], STAT2 [66], TLR5 [67], PTGS2 [68], RNF213 [69], ZDHHC17 [70], JAK2 [71], TLR8 [72], NOTCH2 [73], PDGFC (platelet derived growth factor C) [74], CMPK2 [75], TLR2 [76], CYP1B1 [77], CCR1 [78], HDAC9 [79], IL1RN [80], GCH1 [81], LYST (lysosomal trafficking regulator) [82], PELI1 [83], EGR1 [84], SNX10 [85], CA2 [86], ZEB2 [87], HIF1A [88], PLA2G7 [89], CCR2 [90], GAB1 [91], IRAK3 [92], LDLR (low density lipoprotein receptor) [93], TLR6 [94], SIRT1 [95], NOD2 [96], ATP10D [97], ELOVL6 [98], VCAN (versican) [99], TET2 [100], TET3 [101], ZBTB20 [102], HS3ST1 [103], PF4 [104], DNAJC2 [105], NFIA (nuclear factor I A) [106], CCR7 [107], PRDX2 [108], ADK (adenosine kinase) [109], TCF7 [110], LGALS3 [111], OLFM2 [112], HDAC11 [113] and ARPC1B [114] are significantly related to the atherosclerosis. Studies have revealed that KCNJ2 [115], TLR4 [116], JAK2 [117], TLR2 [118], EGR1 [119], GAB1 [120], ZBTB11 [121], BIN1 [122], TCF4 [123], PPP1R13L [124], TRPM4 [125], LGALS3 [126] and SNTA1 [127] plays a key role in cardiac arrhythmia. Recently, increasing evidence demonstrated that KCNJ2 [128], TLR4 [129], CYP2D6 [130], TLR2 [129], SNX10 [131], SIRT1 [132], PF4 [133], PCYT2 [134] and LGALS3 [135] were altered expression in atrial fibrillation. Studies had shown that KCNJ2 [136], ARG1 [137], TLR4 [138], IFIH1 [139], FBN2 [140], PDK4 [141], TLR8 [142], PDGFC (platelet derived growth factor C) [143], FNIP1 [144], TLR2 [145], PTPN11 [146], LATS2 [147], GCH1 [148], ARFGEF2 [149], CA2 [150], PTPRC (protein tyrosine phosphatase receptor type C) [151], CCR2 [152], GAB1 [153], VEGFA (vascular endothelial growth factor A) [154], UBR1 [155], PHLPP1 [156], MTM1 [157], FMR1 [158], SIRT1 [159], NOD2 [160], MYOF (myoferlin) [161], OSBPL11 [162], ZBTB11 [121], UTRN (utrophin) [163], ZNF593 [164], CCR7 [165], PRDX2 [166], BIN1 [167], NFIC (nuclear factor I C) [168], TCF4 [169], PPP1R13L [124], NDUFB11 [170], TAX1BP3 [171], TRPM4 [172], NMRAL1 [173], LGALS3 [126] and NAA10 [174] were associated with cardiomyopathy. CD274 [175], CEACAM1 [176], STAT1 [177], ARG1 [178], TLR4 [179], LRRK2 [180], ABCA1 [181], IFIH1 [182], TLR5 [183], PTGS2 [184], CYP2D6 [185], RNF213 [186], C9ORF72 [187], JAK2 [188], TLR8 [189], NOTCH2 [190], PDGFC (platelet derived growth factor C) [191], TLR2 [192], PRKAB2 [193], HDAC9 [194], NCOA4 [195], LATS2 [196], DICER1 [197], IL1RN [198], GCH1 [148], EGR1 [199], HIPK3 [200], ZEB2 [201], HIF1A [202], PLA2G7 [203], DOCK8 [204], CCR2 [205], PPP1R15B [206], GCLC (glutamate-cysteine ligase catalytic subunit) [207], VEGFA (vascular endothelial growth factor A) [208], SORL1 [209], OGT (O-linked N-acetylglucosamine (GlcNAc) transferase) [210], EIF2AK3 [211], SLC40A1 [212], PHLPP1 [213], IGF2R [214], LPIN2 [215], SIRT1 [216], VPS13C [217], ACSL4 [218], ELOVL6 [219], FGL2 [220], ERO1B [221], XAF1 [222], TET2 [223], TET3 [224], PF4 [225], POU2F1 [226], PC (pyruvate carboxylase) [227], NDUFS8 [228], PRDX2 [229], RUNX3 [230], HSPB1 [231], TCF4 [169], TCF7 [232], IGFBP4 [233], HSPG2 [234], NMRAL1 [173], AQP3 [235] and LGALS3 [236] might be a potential therapeutic target for diabetes mellitus. Studies have found that CEACAM1 [237], STAT1 [238], ARG1 [239], TLR4 [240], LRRK2 [241], ABCA1 [242], PTGS2 [243], CYP2D6 [244], JAK2 [245], TLR2 [246], DUSP6 [247], CYP1B1 [248], CCR1 [249], HDAC9 [250], LATS2 [251], IL1RN [252], GCH1 [253], PELI1 [254], EGR1 [255], HIPK3 [256], CCR2 [257], GCLC (glutamate-cysteine ligase catalytic subunit) [258], KLF3 [259], VEGFA (vascular endothelial growth factor A) [260], ITGB1 [261], RASA1 [262], PTPN12 [263], SNRK (SNF related kinase) [264], PRKAR1A [265], LDLR (low density lipoprotein receptor) [266], SIRT1 [267], NOD2 [268], VCAN (versican) [269], TET2 [270], PFKFB2 [271], ZBTB20 [272], MYBL2 [273], PF4 [274], VEGFB (vascular endothelial growth factor B) [275], CCR7 [276], PRDX2 [277], HSPB1 [278], ZNF791 [279], IGFBP4 [280], ESF1 [281], SNHG8 [282], LGALS3 [283] and LGMN (legumain) [284] are altered expression in myocardial infarction. Altered expression of CEACAM1 [176], ACSL1 [285], STAT1 [286], TLR4 [287], ABCA1 [288], TLR5 [183], F2RL1 [289], CYP2D6 [290], PDK4 [291], RNF213 [186], JAK2 [292], TLR8 [189], NOTCH2 [293], CENPJ (centromere protein J) [294], FNIP1 [295], TLR2 [296], KIDINS220 [297], DUSP6 [298], CYP1B1 [299], S1PR3 [300], NCOA2 [301], HDAC9 [302], PELI1 [303], EGR1 [304], HIF1A [305], CCR2 [306], IRAK3 [307], KLF3 [308], VEGFA (vascular endothelial growth factor A) [309], OGT (O-linked N-acetylglucosamine (GlcNAc) transferase) [310], PHLPP1 [213], FMR1 [311], IGF2R [312], PRKAR1A [313], WDR11 [314], LDLR (low density lipoprotein receptor) [315], TLR6 [316], SIRT1 [317], ALDH1A1 [318], NOD2 [319], ACSL4 [320], ELOVL6 [321], VCAN (versican) [322], CLIC5 [323], CLCN3 [324], OSBPL11 [162], HELZ2 [325], TET2 [326], KDM6A [327], PHF2 [328], PHGDH (phosphoglycerate dehydrogenase) [329], VEGFB (vascular endothelial growth factor B) [330], ZBTB7A [331], PC (pyruvate carboxylase) [332], CCR7 [333], PRDX2 [334], MAF1 [335], HSPB1 [336], ESF1 [281], LGALS3 [337], OLFM2 [338] and HDAC11 [339] had been confirmed in obesity. CEACAM1 [340], ACSL1 [341], TLR4 [342], ABCA1 [343], TLR5 [344], CYP2D6 [345], JAK2 [346], NOTCH2 [347], DDX3X [348], NCOA4 [349], EGR1 [350], IQGAP2 [351], GCLC (glutamate-cysteine ligase catalytic subunit) [352], VEGFA (vascular endothelial growth factor A) [353], ITGB1 [354], LDLR (low density lipoprotein receptor) [355], TLR6 [316], SIRT1 [356], FGL2 [357], TET2 [358], PHF2 [328], VEGFB (vascular endothelial growth factor B) [359], SELENOM (selenoprotein M) [360], TRPM4 [361], OLFM2 [362] and ATAD3A [363] are thought to be involved in non-alcoholic fatty liver disease. Altered expression of ATOH8 [364], STAT1 [365], ARG1 [366], TLR4 [367], VNN1 [368], ABCA1 [369], IFIH1 [370], PTGS2 [371], F2RL1 [289], CYP2D6 [372], PDK4 [373], RNF213 [374], JAK2 [375], NOTCH2 [376], PDGFC (platelet derived growth factor C) [377], TLR2 [378], CYP1B1 [379], IL1RN [380], GCH1 [381], EGR1 [382], HIF1A [383], PLA2G7 [384], CCR2 [385], GAB1 [386], VEGFA (vascular endothelial growth factor A) [387], OGT (O-linked N-acetylglucosamine (GlcNAc) transferase) [388], OXR1 [389], IRF9 [390], FMR1 [391], LDLR (low density lipoprotein receptor) [392], SIRT1 [393], NOD2 [394], ATP13A3 [395], VCAN (versican) [396], FGL2 [397], TET2 [398], KDM6A [399], KLHL2 [400], CAVIN1 [401], TNFRSF4 [402], PF4 [403], VEGFB (vascular endothelial growth factor B) [330], CCR7 [404], PRDX2 [405], HSPB1 [406], TCF4 [407], MRPL4 [408], PHF14 [409], TRPM4 [410], AQP3 [411] and LGMN (legumain) [412] were observed to be associated with the progression of hypertension. Recent studies have shown that altered expression of DDX58 [413], STAT1 [414], TLR4 [415], CYP2D6 [416], JAK2 [417], TLR2 [418], DUSP6 [419], HDAC9 [420], LATS2 [421], CA2 [150], HIPK3 [422], CCR2 [423], GAB1 [120], UFL1 [424], OGT (O-linked N-acetylglucosamine (GlcNAc) transferase) [425], PRKAR1A [265], SIRT1 [426], FGL2 [427], TET2 [428], ASCC2 [429], BIN1 [430], HSPB1 [431], IGFBP4 [432], TRPM4 [433] and LGALS3 [434] might be associated with the progression of heart failure. STAT1 [435], TLR4 [436], ABCA1 [437], PTGS2 [68], CYP2D6 [438], CR1 [439], PDK4 [440], RNF213 [441], ZDHHC17 [70], TLR8 [442], PDGFC (platelet derived growth factor C) [443], TLR2 [444], CYP1B1 [445], HDAC9 [446], IL1RN [447], GCH1 [448], EGR1 [449], ZEB2 [450], PLA2G7 [451], CCR2 [452], GCLC (glutamate-cysteine ligase catalytic subunit) [258], VEGFA (vascular endothelial growth factor A) [453], CD46 [454], NFKBIZ (NFKB inhibitor zeta) [455], LDLR (low density lipoprotein receptor) [456], TLR6 [457], SIRT1 [458], NOD2 [459], FGL2 [460], IDH1 [461], TET2 [462], PFKFB2 [463], KDM6A [464], IKZF2 [465], ZNF606 [466], PF4 [467], CCR7 [468], RUNX3 [469], TCF7 [470], PPBP (pro-platelet basic protein) [471], IGFBP4 [280], HSPG2 [472] and LGALS3 [236] expression might be regarded as an indicator of susceptibility to CAD. STAT1 [473], TLR4 [367], JAK2 [474], PDGFC (platelet derived growth factor C) [143], CYP1B1 [475], HDAC9 [476], ZEB2 [477], GAB1 [478], IRF9 [479], PHLPP1 [480], SIRT1 [481], NOD2 [482], JMJD1C [483], CLK4 [484], CAVIN1 [485], VEGFB (vascular endothelial growth factor B [486], PRDX2 [487], MAF1 [488], TCF7 [489], IGFBP4 [490], TRPM4 [491], NAA10 [174] and SNTA1 [492] had been demonstrated to participate in cardiac hypertrophy. STAT1 [493], JAK2 [494], NOTCH2 [495], PDGFC (platelet derived growth factor C) [143], RASA1 [262], TLR6 [496], NOD2 [482], JMJD1C [483], CAVIN1 [485], POU2F1 [497], RPS5 [498] and LGALS3 [499] are involved in the regulation of cardiac fibrosis. TLR4 [500], ABCA1 [501], CCR2 [502], LDLR (low density lipoprotein receptor) [503], SIRT1 [504] and NOD2 [505] might crucially contribute to the development of hypercholesterolemia. ABCA1 [506], TLR5 [507], PTGS2 [508], TGFA (transforming growth factor alpha) [509], PDK4 [510], JAK2 [511], TLR2 [512], NEK7 [513], CCR1 [514], BACH1 [515], NCOA4 [195], LATS2 [516], PELI1 [517], EGR1 [518], CYBB (cytochrome b-245 beta chain) [519], MEFV (MEFV innate immuity regulator, pyrin) [520], CLEC4E [521], GCLC (glutamate-cysteine ligase catalytic subunit) [522], KLF3 [523], TP53INP1 [524], ITGB1 [525], IRF9 [526], PHLPP1 [527], NOD2 [528], ACSL4 [529], FGL2 [530], PF4 [531], VEGFB (vascular endothelial growth factor B) [532], CCR7 [533], IGFBP4 [534], TRPM4 [535], BAG1 [536], LGALS3 [537] and ATAD3A [538] could induce ischemic heart disease. Therefore, the data suggest that the identified DEGs which enriched in GO terms and pathways might participant in the development of CAD and its associated complications, and contribute to CAD and CAD related complications treatment.

Notably, the PPI network linked to CAD and CAD related complications were composed of functional proteins that interacted with each other to participate in biological signal pathways, gene expression control, energy, and protein, lipid and carbohydrate metabolism. Key modules might strongly contribute to the occurrence and development of CAD and its associated complications, and have high diagnostic value. EGR1 [84], SIRT1 [95], STAT1 [60], HIF1A [88], HDAC11 [113], STAT2 [66] and IFIT1 [51] might play an important role in regulating the genetic network related to the occurrence, development of atherosclerosis. Altered expression of EGR1 [119] might be associated with cardiac arrhythmia. EGR1 [199], SIRT1 [216], STAT1 [177], LRRK2 [180], HIF1A [202] and IFITM3 [49] genes might be related to the pathophysiology of diabetes mellitus and may be one of the markers for the early diagnosis of diabetes mellitus. EGR1 [255], SIRT1 [267], STAT1 [238] and LRRK2 [241] are mainly involved in myocardial infarction. EGR1 [304], SIRT1 [317], STAT1 [286], HIF1A [305], HDAC11 [339] and IFITM3 [48] genes have been demonstrated to be responsible for the progression of obesity. Studies have confirmed that EGR1 [350] and SIRT1 [356] are altered expressed in non-alcoholic fatty liver disease. Previous studies have shown that EGR1 [382], SIRT1 [393], STAT1 [365], HIF1A [383] and IRF9 [390] might influence the prognosis in hypertension. EGR1 [449], SIRT1 [458] and STAT1 [435] are a genes which plays a role in diagnosis of CAD. EGR1 [518] and IRF9 [526] have a significant prognostic potential in ischemic heart disease. SIRT1 [132] is molecular marker for the diagnosis and prognosis of atrial fibrillation. Studies had shown that SIRT1 [159] expression was associated with cardiomyopathy. Altered expression of SIRT1 [426] and STAT1 [414] are associated with clinical prognosis in heart failure. SIRT1 [481], STAT1 [473] and IRF9 [479] were a diagnostic biomarkers of cardiac hypertrophy and could be used as therapeutic targets. Studies had shown that SIRT1 [504] and IFITM3 [49] were associated with hypercholesterolemia. Studies have found that STAT1 [493] is altered expression in cardiac fibrosis, which can be used as a prognostic marker for cardiac fibrosis. Our findings suggested CSNK2B, RPS3, RPS2, RPS4X, EIF2AK2, GBP1, IFIT2, RPS14, RPS29, RPS9, RPS15, RPLP1, RPS19, RPL3, RPLP2 and RPL36 as novel diagnostic biomarkers for CAD and CAD related complications. Therefore, these biomarkers might be used as potential effective candidates for early diagnosis or prognosis of CAD and CAD related complications. Importantly, study on ribosomal proteins might be a significant new direction for the diagnosis, prognosis and treatment of CAD and CAD related complications.

Based on the miRNA-hub gene regulatory network and TF-hub gene regulatory network constructed by the online database miRNet and NetworkAnalyst, we identified hub genes, miRNAs and TFs. Previous study showed that DDX3X [348], EGR1 [350] and SREBF1 [539] were associated with non-alcoholic fatty liver disease. EGR1 [84], HIF1A [88], STAT1 [60], hsa-mir-16-5p [540], hsa-mir-532-3p [541], hsa-mir-503-5p [542], SREBF1 [543] and FOXP3 [544] plays a pivotal role in the atherosclerosis. The results showed that EGR1 [119] expression participate in the progression of cardiac arrhythmia. EGR1 [199], HIF1A [202], STAT1 [177], hsa-mir-31-5p [545], SREBF1 [546] and FOXP3 [547] were observed to be associated with the progression of diabetes mellitus. EGR1 [255], STAT1 [238] and FOXP3 [548] regulates the development of myocardial infarction. EGR1 [304], HIF1A [305], STAT1 [286], hsa-mir-31-5p [549], hsa-mir-365b-3p [550], SREBF1 [551] and FOXP3 [552] might be associated with the prognosis of obesity. EGR1 [382], HIF1A [383], STAT1 [365] and FOXP3 [553] mainly involved in the occurrence of hypertension. Aberrant expression of EGR1 [449], STAT1 [435], hsa-mir-16-5p [554], IRF8 [555] and FOXP3 [556] have been observed in CAD. Studies have found that EGR1 [518] and hsa-mir-16-5p [557] expression is an independent predictor of prognosis in patients with ischemic heart disease. STAT1 [414] and FOXP3 [558] migt be a prognostic biomarkers and potential therapeutic targets for patients with heart failure. Studies have found that STAT1 [493] and RPS5 [498] mRNA expressions are an independent predictor of prognosis in patients with cardiac fibrosis. hsa-mir-16-5p [557] and FOXP3 [559] are associated with the prognosis of cardiomyopathy. FOXP3 [600] was identified as a candidate biomarker for the diagnosis and treatment of hypercholesterolemia. We identified DDX5, NCOA3, RPS3, RPS4X, RPS14, FBL, RPL3, RPS19, RPL18, hsa-mir-149-3p, hsa-mir-500b-5p,’ hsa-mir-548y, hsa-mir-518b, hsa-mir-5699-3p, LMO2, MYCN, SIN3B, GATA1, RCOR3, DMRT1 and POU5F1 that might serve as novel biomarkers for CAD and CAD related complications.

In conclusion, a comprehensive bioinformatics analysis of DEGs and pathways involved in the occurrence and advanacement of CAD and CAD related complications was performed, and we find and obtained essential genes and pathways contributing to the progression of CAD and CAD related complications to improve prognosis. Moreover, these results migt promote the understanding of molecular mechanisms and clinically related molecular targets for prognosis in CAD and CAD related complications and provide novel insight into the occurrence and adancement of CAD and CAD related complications.

## Acknowledgement

I thank Gualtiero Ivanoe Colombo, Centro Cardiologico Monzino IRCCS, Immunology and Functional Genomics, Via Carlo Parea, 4, Milano, Italy, very much, the author who deposited their NGS dataset GSE202625, into the public GEO database.

## Ethical approval

This article does not contain any studies with human participants or animals performed by any of the authors.

## Informed consent

No informed consent because this study does not contain human or animals participants.

## Availability of data and materials

The datasets supporting the conclusions of this article are available in the GEO (Gene Expression Omnibus) (https://www.ncbi.nlm.nih.gov/geo/) repository. [(GSE202625) https://www.ncbi.nlm.nih.gov/geo/query/acc.cgi?acc=GSE202625]

## Consent for publication

Not applicable.

## Author Contributions

B. V. - Writing original draft, and review and editing

C. V. - Software and investigation

